# Myeloid-derived β-hexosaminidase is essential for neuronal health and lysosome function: implications for Sandhoff disease

**DOI:** 10.1101/2024.10.21.619538

**Authors:** Kate I. Tsourmas, Claire A. Butler, Nellie E. Kwang, Zachary R. Sloane, Koby J. G. Dykman, Ghassan O. Maloof, Christiana A. Prekopa, Robert P. Krattli, Sanad M. El-Khatib, Vivek Swarup, Munjal M. Acharya, Lindsay A. Hohsfield, Kim N. Green

## Abstract

Lysosomal storage disorders (LSDs) are a large disease class involving lysosomal dysfunction, often resulting in neurodegeneration. Sandhoff disease (SD) is an LSD caused by a deficiency in the β subunit of the β-hexosaminidase enzyme (*Hexb*). Although *Hexb* expression in the brain is specific to microglia, SD primarily affects neurons. To understand how a microglial gene is involved in maintaining neuronal homeostasis, we demonstrated that β-hexosaminidase is secreted by microglia and integrated into the neuronal lysosomal compartment. To assess therapeutic relevance, we treated SD mice with bone marrow transplant and colony stimulating factor 1 receptor inhibition, which broadly replaced *Hexb*^-/-^ microglia with *Hexb*-sufficient cells. This intervention reversed apoptotic gene signatures, improved behavior, restored enzymatic activity and *Hexb* expression, ameliorated substrate accumulation, and normalized neuronal lysosomal phenotypes. These results underscore the critical role of myeloid-derived β- hexosaminidase in neuronal lysosomal function and establish microglial replacement as a potential LSD therapy.

## INTRODUCTION

Lysosomal storage disorders (LSDs) are a class of genetic diseases caused by deficiencies in lysosomal enzymes or membrane proteins, resulting in the accumulation of excess substrate that causes to lysosomal dysfunction and/or cell death^1^. Though individual LSDs are rare, LSDs have an overall frequency of approximately 1 in 5,000 live births^2^. The majority of LSDs are characterized by a progressive neurodegenerative phenotype with an infantile or early childhood onset. One such disorder is Sandhoff disease (SD), which is caused by a complete loss of the β-Hexosaminidase enzyme (Hexβ)^3,4^. Hexβ is a dimeric enzyme composed of either an alpha (HEXA) and beta (HEXB) subunit or two beta subunits, with the respective subunits encoded by the *Hexa* and *Hexb* genes^5^. In SD, a disruption to the *Hexb* gene results in the complete loss of functioning Hexβ enzyme and the inability to break down its substrates, namely GM2 ganglioside glycolipids and other glycolipids/glycoproteins, which accumulate in the central nervous system (CNS) and peripheral organs^6,7^. While some studies have linked dysregulated glycolipid metabolism to the pathogenesis of neurological disorders, it remains unclear how glycolipid accumulation causes neurodegeneration^8,9^. In humans and *Hexb*^-/-^ mice, which closely recapitulate features of SD, the disease manifests as a rapidly progressive neurodegenerative disorder characterized by extensive neuroinflammation followed by mass neuronal apoptosis, severe motor and developmental impairments, and death by age four in humans and 18-20 weeks in mice^3,10,11^ At present, there is no available curative or disease-modifying treatment for SD.

Bone marrow transplant (BMT) has been shown to be an effective treatment for several LSDs and has been investigated as a potential treatment strategy for SD^12,13^. However, in human patients with SD and the closely related gangliosidosis Tay-Sach’s disease (TSD), BMT has been ineffective and repeatedly failed to meaningfully extend lifespan^10,14,15^. BMT has shown limited efficacy in *Hexb*^-/-^ mice, reducing neuroinflammation and partially prolonging life; however, it ultimately failed to normalize lifespan or correct CNS pathology^11,16^. *Hexb*^-/-^ BMT-treated mice exhibit a reduction of GM2 ganglioside burden, but only in peripheral organs. This insufficiency may be linked to a failure to replace microglia, the primary myeloid cells of the CNS, using traditional BMT. Although studies have shown that BMT can reduce enzyme substrate accumulation in peripheral organs of many LSDs, including SD, the CNS has been notoriously difficult to correct in many cases^12^. This is likely attributable to the relatively higher rates of myeloid cell replacement in these organs compared to the nominal replacement rate of CNS myeloid cells with bone-marrow derived cells, ranging from <10 to ∼30%^17–21^. Microglia are heavily implicated in the development of SD and *Hexb* expression in the CNS has been reported highly specific, or exclusive, to microglia^11,22–26^. However, it remains unclear how deficits in a myeloid cell gene (i.e., loss of *Hexb*) result in the primarily neuronal pathology and cell death observed in the SD CNS. In this study, we sought to develop an approach that would allow us to better understand the relationship between myeloid *Hexb* expression and neuronal pathology in SD while improving upon the shortcomings of BMT and other treatment modalities with incomplete efficacy.

We and others have previously identified a means to reliably replace the microglial population with bone marrow-derived myeloid cells (BMDMs) via pharmacological inhibition of the colony stimulating factor 1 receptor (CSF1R) combined with BMT^19,27–29^. This approach results in the broad and brain-wide replacement of microglia with BMDMs, achieving 70-99% replacement. Busulfan-based BMT + CSF1R inhibitor (CSF1Ri) approaches have recently been utilized to therapeutically replace microglia in other mouse models of neurodegenerative disease, including progranulin deficiency, experimental autoimmune encephalomyelitis, and Prosaposin deficiency, all with promising results^30–32^. Here, we employ a BMT + CSF1Ri-based microglial replacement strategy in the *Hexb*^-/-^ mouse model of SD and demonstrate that delivery of *Hexb-*expressing cells via myeloid cell replacement rescues neuron-related molecular and functional outcomes. In neurons, we observe reversed expression of apoptosis-associated genes, resolution of glycolipid/glycoprotein storage, clearance of accumulated lysosomal components, and reduction of vacuolization following microglial replacement with combined BMT + CSF1Ri treatment. Subsequent cell culture experiments reveal that microglia secrete enzymatically active Hexβ protein in a Ca^2+^-dependent, P2X7-mediated manner, and that neurons take up extracellular Hexβ protein and integrate it into the lysosomal compartment. These experiments not only provide evidence for a promising novel treatment strategy for SD and other CNS LSDs, but also indicate that myeloid-derived Hexβ is essential for neuronal health and lysosomal function.

## RESULTS

### Spatial transcriptomic analysis reveals broad genetic changes induced by loss of Hexb

To explore the molecular underpinnings of SD, we performed multi-plex single cell resolution *in situ* RNA analysis by spatial molecular imaging^33^ on *Hexb*^-/-^ mouse brains, in the first characterization of its kind for SD. Previous studies have shown that *Hexb*^-/-^ mice faithfully recapitulate features of human SD, including neuroinflammation/microglial activation, GM2 ganglioside accumulation, and severe motor decline^7^ (Fig. 1a). For this experiment, wildtype control and *Hexb*^-/-^ mice (n=3 per group) were sacrificed at 16 weeks, a humane endpoint at which *Hexb*^-/-^ mice present severe motor phenotypes. Fixed brains were sectioned sagittally at 10µm, imaged with rRNA, Histone, DAPI, and GFAP markers for cell segmentation, and analyzed for 1,000 genes using the Nanostring CosMx Spatial Molecular Imager platform (316 total FOVs, ∼52 FOVs per brain section) (Fig. 1a, b; examples of cell segmentation in Extended Data Fig. 1a). A spatial transcriptomic approach is advantageous in its ability to identify brain regions more affected by disease, while also offering a high percentage of cell capture (∼90%) and relative reduction in sampling bias in comparison to single-cell RNA sequencing (RNA-seq) approaches. With this approach, we captured 196,533 cells with a mean transcript count of ∼800 transcripts per cell. Unbiased cell clustering identified 39 transcriptionally distinct clusters (Fig. 1c; Extended Data Fig. 1b). Clusters were annotated with a combination of automated and manual approaches: 1) label annotations from the Allen Brain Atlas single-cell RNA-seq reference dataset (for cortex and hippocampus) were projected onto our spatial transcriptomics dataset^34^, and 2) cluster identities were further refined via manual annotation based on gene expression of known marker genes and location in XY space (Fig. S2, S3). We identified 14 clusters of excitatory neurons, five clusters of inhibitory neurons, six astrocyte clusters, two myeloid clusters, four oligodendrocyte clusters, one oligodendrocyte precursor (OPC) cluster, three vasculature-associated clusters, two endothelial clusters, and two uncategorized (other) clusters. Projecting cell subclusters in XY space shows clear separation between anatomical regions and cortical layers (Fig. 1c, Extended Data Fig. 1c).

**Fig. 1:**
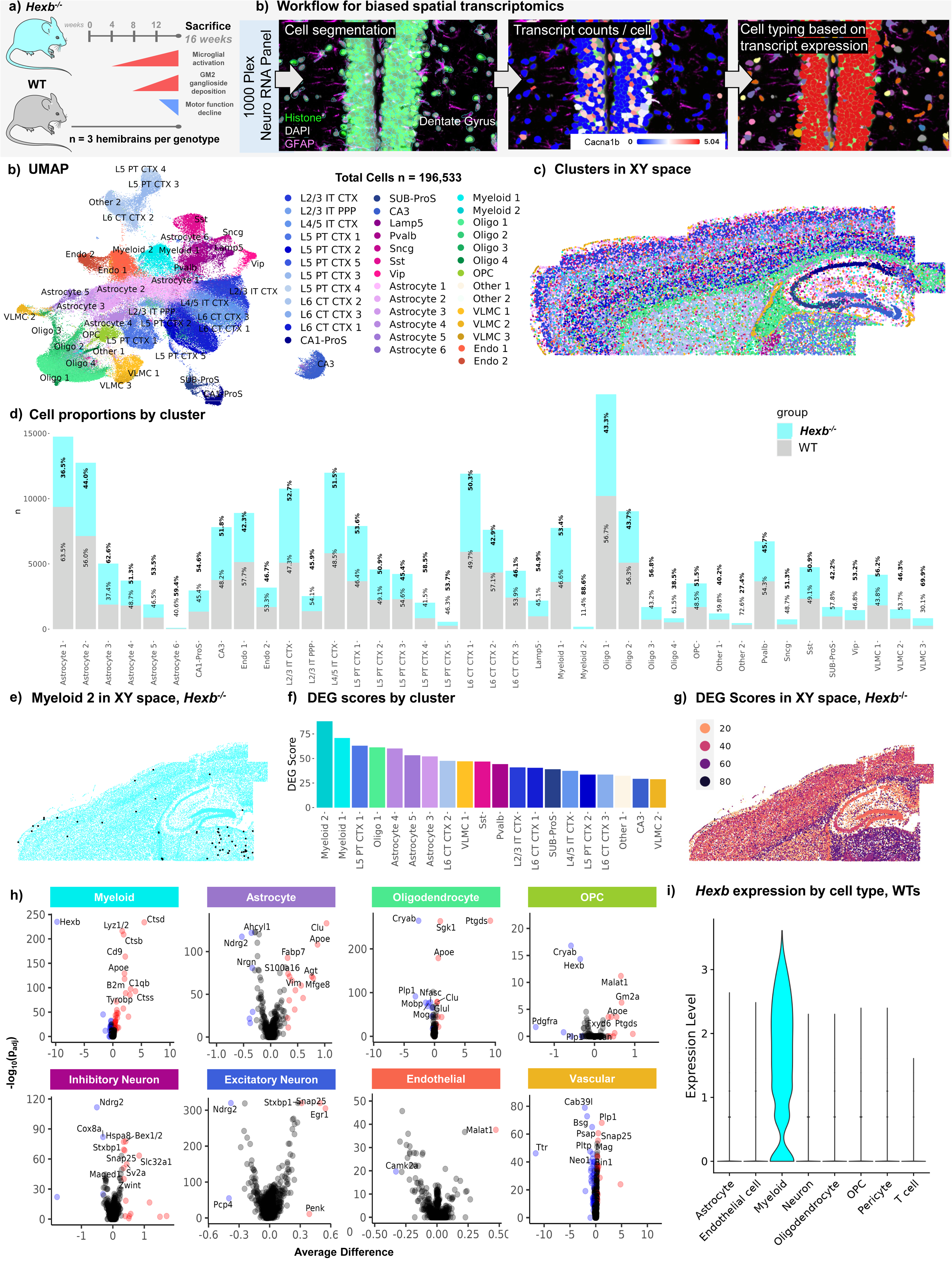
Spatial transcriptomic analysis of the SD mouse brain identifies disease-associated gene expression signatures. (a) Timeline of symptom progression in *Hexb^-/-^* Sandhoff disease model mice up to point of sacrifice at 16 weeks (n=3/genotype, *Hexb^-/-^*and wildtype (WT) control). Microglial/myeloid activation begins at ∼4 weeks, accumulation of GM2 ganglioside glycolipid can be detected ∼8 weeks, and motor deterioration begins ∼12 weeks. (b) Experimental workflow for targeted 1000- plex single-cell spatial transcriptomics. Fields of view (FOVs) were selected in cortex, corpus callosum, hippocampus, and upper regions of caudate and thalamus of each sagittal section, then imaged with DNA, rRNA, Histone, and GFAP markers for cell segmentation. Transcript counts for each gene were acquired per cell. (c) Uniform Manifold Approximation and Projection (UMAP) of 196,533 cells across 6 brains. Clustering at 1.0 resolution yielded 39 clusters, which were annotated with a combination of automated and manual approaches with reference to Allen Brain Atlas singe-cell RNA-seq cell types, gene expression, and anatomical location in space. (c) 39 clusters plotted in XY space. (d) Bar graph of proportions of cell counts by subcluster per genotype. (e) Myeloid 2 subcluster (black) overlaid above representative *Hexb^-/-^*brain plotted in XY space. (f) Descending bar graph of top 20 subclusters with highest differentially expressed gene (DEG) scores. Following differential gene expression analysis, DEG score was calculated per subcluster by summing the absolute value of the log2 fold change values for all DEGs between *Hexb^-/-^* and WT control with a padj value below 0.05. (g) Projection of subclusters colored by DEG score in XY space in representative *Hexb^-/-^* brain. (h) Volcano plots of DEGs between *Hexb^-/-^* and WT control for each broad cell type. (i) Violin plot of *Hexb* transcript counts in cell types demonstrating myeloid-specific expression.

Analysis of the distribution of cell counts within each cluster by genotype revealed a shift in the glial cell populations in *Hexb*^-/-^ mice (Fig. 1d). Interestingly, *Hexb*^-/-^ mice exhibited a high proportion of cells in the Myeloid 2 subcluster (88.6%) compared to WT mice, indicating a near-exclusive presence of this cell type in the *Hexb*^-/-^genotype. The Myeloid 2 subcluster is enriched in genes *H2-Aa*, *Cd74*, *H2-Ab1*, *Lyz1/2*, and *Tyrobp,* which have been associated with monocyte/monocyte identity and regulation of the immune response in monocytes^35–38^. In line with this, infiltrating monocytes/macrophages have previously been reported in small quantities in *Hexb*^-/-^ brains^39^. Plotting of the Myeloid 2 subcluster in XY space indicates that the cells are localized to the thalamus and throughout the cortex (Fig. 1e). *Hexb*^-/-^ mice also showed a larger proportion of cells in the Astrocyte 3 subcluster, with a lower proportion of cells in the Astrocyte 1 and Oligodendrocyte 4 subclusters. Interestingly, the Astrocyte 3 subcluster is enriched with disease-associated or reactive astrocyte genes (*Gfap*, *Aqp4*, *Vim*, *Clu*, *Gja1*)^40–44^. It should be noted that there were no major differences in the overall number of astrocytes, oligodendrocytes, or myeloid cells (Extended Data Fig. 1d). Surprisingly, we also observed no notable reductions in neuronal subclusters counts or overall neuronal counts in the *Hexb*^-/-^ brain in comparison to WT, indicative of a lack of overt neuronal loss by 16 weeks in SD mice (Extended Data Fig. 1d). These data suggest that at the 16-week time point, *Hexb* deficiency has an impact on glial populations without affecting neuronal counts.

Next, we performed differential gene expression (DGE) analysis on all cell clusters to assess gene expression changes associated with loss of *Hexb* (Extended Data Fig. 4). Each cluster was then assigned a differentially expressed gene (DEG) score (Fig. 1f; Extended Data Fig. 3d). DEG score is a measure of the magnitude of gene expression changes between two groups within each cellular subcluster. Subcluster DEG scores were calculated by summing the absolute log2 fold change values of all genes with significant (padj < 0.05) differential expression patterns between *Hexb^-/-^* and WT. This metric allowed us to assess broad alterations in cellular subtypes caused by *Hexb* insufficiency in comparison to WT (Fig. 1f). Myeloid 1 and 2 subclusters had the highest DEG scores, indicating that these cell populations are the most impacted by *Hexb* deficiency. To visualize DEGs across major CNS cell types, we next combined subclusters into broad cell types (i.e., Astrocyte clusters 1-6 were placed in the Astrocyte broad cell type category). DGE analysis of all myeloid subclusters revealed, as expected, a marked downregulation of *Hexb*, accompanied by an upregulation in several genes associated with inflammation: cathepsins (*Ctsb, Ctsd, Ctss*), immune activation (*B2m, Tyrobp*), and macrophage-associated genes (*Lyz1/2, C1qb*)^45–49^ (Fig. 1h). DGE analysis of oligodendrocytes, which included a subcluster with a high DEG score (Oligo 1), identified several genes associated with CNS inflammatory/stress response (*Ptgds, Sgk, Cryab*) and demyelination (*Mog, Mobp, Plp1)*^50–53^ (Fig. 1i); oligodendrocyte expression of *Ptgds,* in particular, has been shown to induce neuronal apoptosis^54^. Astrocyte subclusters 3-5 also exhibited high DEG scores. Plotting astrocyte DEGs, we found a downregulation in a homeostatic astrocyte gene (*Ndrg2)* and upregulation in markers associated with astrocyte activation and neurotoxicity (*Clu*, *Apoe*, *Fabp7, S100a6, Vim*)^40,42,43,55–60^.

The most affected neuronal subclusters in *Hexb^-/-^* compared to WT in terms of DEG score were the Layer 5 PT CTX 1 and Layer 6 CT CTX 2 clusters, both excitatory neuronal subtypes, followed by the two largest inhibitory subclusters (somatostatin [Sst], parvalbumin [Pvalb]). Plotting top DEGs of both excitatory and inhibitory neurons revealed an upregulation of genes associated with development and synaptic function (*Snap25, Stxbp1)*^61,62^. Excitatory neurons displayed fewer significant DEGs than inhibitory neurons; however, they also exhibited alterations in genes associated with apoptosis. We observe a strong upregulation in early growth response • (*Egr1*), a gene previously implicated in orchestrating neuronal apoptosis and modulating expression of stress-responsive transcription factor EB (TFEB), a master regulator of lysosomal biogenesis and autophagy; *Egr1* has also been shown to be upregulated under conditions of lysosomal dysfunction^63–65^. We also note a downregulation of Purkinje cell protein 4 (*Pcp4*), a gene that is decreased in various neurodegenerative diseases and linked to apoptosis^66^. Inhibitory neurons also exhibited a litany of DEGs associated with perturbed neurotransmission and apoptotic processes (*Sv2a, Slc32a1, Bex1/2, Zwint*, *Maged1)* and cellular stress and metabolic processes (*Hsp8a, Cox8a); Hsp8a* is also a key regulator of lysosome activity and autophagy^67–74^. Although we did not observe gross changes in neuronal counts at this stage of disease in *Hexb*^-/-^ mice, these neuronal gene expression changes are notable in their indication of broad neuronal dysregulation and the initiation of apoptotic processes. The selected endpoint thus may have captured the state of the SD CNS shortly preceding overt neuronal loss, as *Hexb*^-/-^ mice generally survive to 18-20 weeks and disease course is rapid.

We next plotted all subcluster DEG scores in XY space to visualize broad gene expression changes spatially and identify region-specific vulnerabilities (Fig. 1g). The regions populated by cells with the highest DEG scores were the thalamus and corpus callosum. Cells throughout the cortex had higher DEG scores than those of the hippocampus and caudoputamen. These region-specific effects align well with previous results from human SD patients and mice, which report white matter neurodegeneration, thalamic hyperintensities/hyperdensities, and cortical atrophy with relative sparing of the caudate^75–77^. Notably, many of the observed gene expression changes between *Hexb^-/-^* and WT mice closely aligned with DEGs previously identified in datasets derived from human SD and TSD patients, including *Ptgds*, *Vim*, *Apoe, Clu, Ctsb, Nrgn,* and *Mbp*^78^. Our DGE analysis provides evidence that CNS cell types of various lineages are affected by SD. Upregulation of genes associated with reactivity in glial cells may contribute to the apoptotic signatures detected in *Hexb^-/-^* neurons. Understanding how differing cell types interact and contribute to neurodegeneration in SD is of great interest to understanding disease pathogenesis and uncovering potential therapeutic opportunities.

Finally, we assessed *Hexb* expression levels in various cell subtypes in WT animals. Interestingly, despite the established role of Hexβ in maintaining neuronal health, we detected *Hexb* transcripts exclusively in myeloid cells of WT animals (Fig. 1i). Very few transcripts were detected in other cell types, including astrocytes, endothelial cells, neurons, oligodendrocytes, OPCs, pericytes, or T cells. Our identification of myeloid-specific *Hexb* expression is in agreement with previous reports of transcript and protein expression patterns, which show specific expression of *Hexb* in microglia in the CNS^25,26^. These data collectively identify the myeloid population as a particularly significant cell type in the SD brain, highlighting it as a promising target for therapeutic intervention.

### Combined BMT + CSF1Ri treatment leads to functional rescue, broad microglial replacement, and normalization of myeloid morphology in Hexb-deficient mice

Given the high potential of a myeloid cell-based therapeutic target for SD, we next sought to replace *Hexb* deficient microglia in *Hexb*^-/-^ mice with *Hexb* sufficient BMDMs from WT donors and assess the viability of microglial replacement as a treatment for SD. Pre-pathological (4-6 weeks of age) *Hexb*^-/-^ mice and WT control mice were treated with BMT by total body irradiation at a dose of 9 Gy and subsequent retro-orbital injection of bone marrow cells (Fig. 2a). Bone marrow cells were isolated from sex-matched CAG-EGFP mice, allowing for visual tracking of donor cells based on GFP expression. We hypothesized that CAG-EGFP donor cells would also allow for normalized *Hexb* expression in the brain. Analysis for chimerism revealed that BMT resulted in an average blood (granulocyte) and bone marrow (hematopoietic stem cell) chimerism rate of ∼95-99%, with no notable differences between genotypes or treatment paradigms (Extended Data Fig. 5a-c). Following BMT, one group of mice was then placed on control diet (WT BMT n=10, *Hexb*^-/-^ BMT n=11). Another group underwent a 2-week post-irradiation recovery period before being treated with the CSF1R inhibitor diet PLX5622 at a dose of 1200 ppm for 7 days to induce widespread microglial depletion. Following this, the inhibitor was withdrawn and the group returned to a control diet, which we have previously show results in efficient replacement of microglia with BMDMs following head irradiation^19^ (WT BMT + CSF1Ri n=10, *Hexb*^-/-^ BMT + CSF1Ri n=10). Untreated mice were also included to serve as controls (*Hexb*^-/-^ control n=10, WT control n=10).

**Fig. 2:**
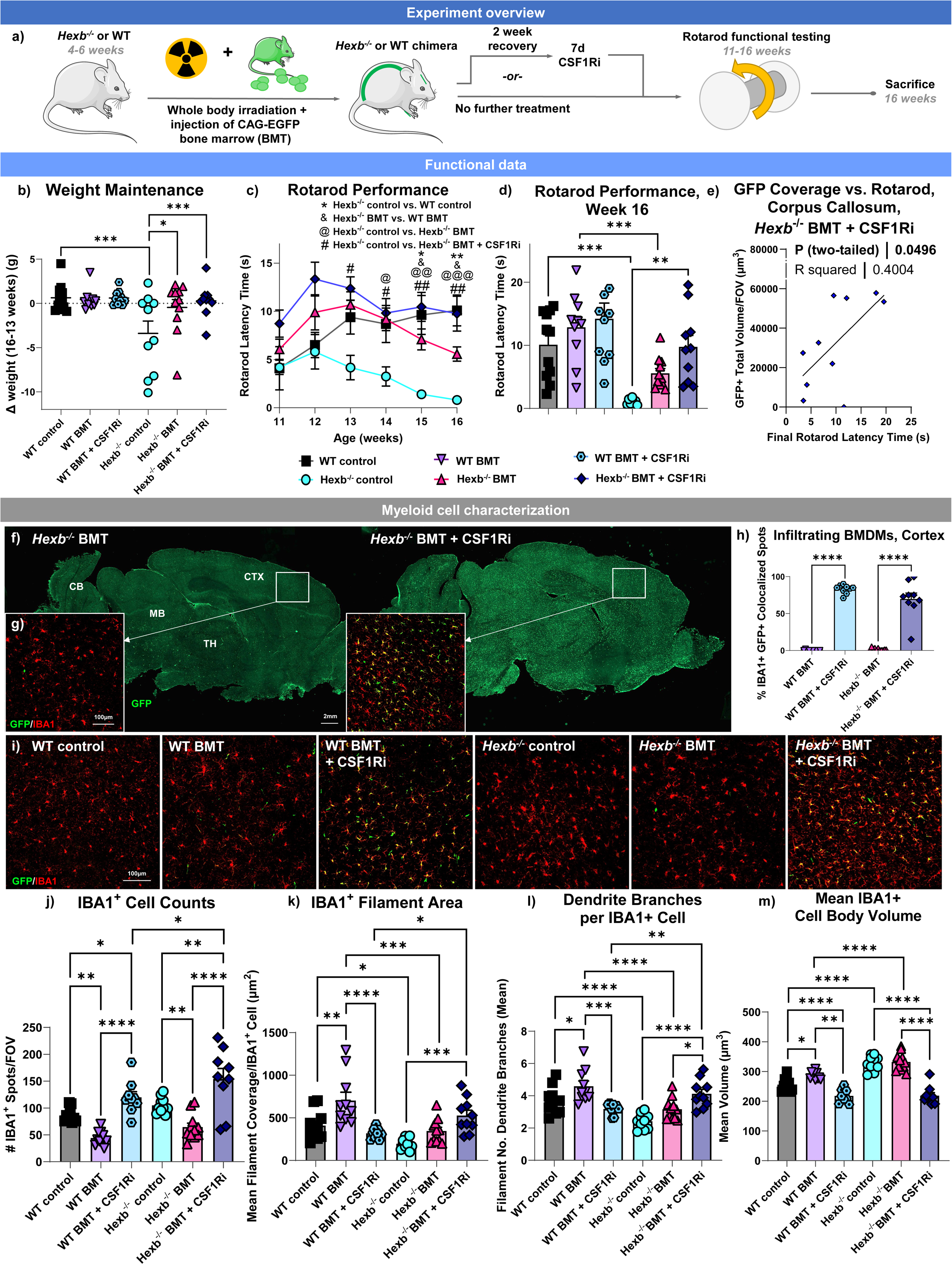
M**i**croglial **replacement in SD leads to functional rescue and normalization of microglial morphology.** (a) Schematic of treatment paradigm. WT and *Hexb*^-/-^ mice were split into 3 groups: untreated control, bone marrow transplant (BMT), and bone marrow transplant plus colony stimulating factor 1 inhibitor treatment (BMT + CSF1Ri). Mice underwent functional testing with the accelerating Rotarod task and were sacrificed at 16 weeks. (b) Categorical scatter plot of change in weight in grams in WT, BMT-treated WT, BMT + CSF1Ri-treated WT, *Hexb*^-/-^, BMT- treated *Hexb*^-/-^, and BMT + CSF1Ri-treated *Hexb*^-/-^ mice between week of sacrifice and week 13 (week 16 weight – week 13 weight). (c) Line graph of average latency-to-fall time in seconds in WT control, *Hexb*^-/-^ control, *Hexb*^-/-^ BMT, and *Hexb*^-/-^ BMT + CSF1Ri groups on Rotarod task per week from 11 to 16 weeks of age. From week 13, groups compared by repeated measures ANOVA with Tukey’s post-hoc testing. Symbols indicate significant differences between *Hexb*^-/-^ control and WT control (*), *Hexb*^-/-^ BMT-treated and WT BMT-treated (&), *Hexb*^-/-^ BMT-treated and *Hexb*^-/-^ control (@), and *Hexb*^-/-^ BMT + CSF1RI and *Hexb*^-/-^ control (#) mice. (d) Scattered bar plot of final week (week 16) Rotarod latency-to-fall time in WT, BMT-treated WT, BMT + CSF1Ri- treated WT, *Hexb*^-/-^, BMT-treated *Hexb*^-/-^, and BMT + CSF1Ri-treated *Hexb*^-/-^ mice. (e) Scatterplot with line of best fit of final week (week 16) Rotarod latency-to-fall time (x axis) versus total green fluorescent protein (GFP, green)^+^ staining volume in upper corpus callosum (y axis) in BMT + CSF1Ri-treated *Hexb*^-/-^ mice. Demonstrates significant (p = 0.0456) positive correlation between corpus callosum GFP^+^ volume and final Rotarod score. (f) Representative 10x whole brain images of sagittal sections from *Hexb*^-/-^ BMT and *Hexb*^-/-^ BMT + CSF1Ri mice immunolabeled for GFP (green), demonstrating CNS infiltration of CAG-EGFP donor-derived cells. CTX, cortex; MB, midbrain; CB, cerebellum; MB, midbrain; TH, thalamus. (g) Representative confocal images of cortex in *Hexb*^-/-^ BMT- treated and *Hexb*^-/-^ BMT + CSF1Ri-treated mice immunolabeled for GFP (green) and myeloid cell marker IBA1 (red), showing colocalization (yellow). (h) Bar graph of quantification of percentage of IBA1^+^ cells with colocalized GFP^+^ per FOV in cortex images from BMT-treated WT, BMT + CSF1Ri-treated WT, BMT-treated *Hexb*^-/-^, and BMT + CSF1Ri-treated *Hexb*^-/-^ mice, indicating ratio of myeloid cells with bone marrow-derived myeloid cell (BMDM) identity. Two-way ANOVA with Sidak’s post-hoc test. (i) Representative confocal images of cortex from WT, BMT-treated WT, BMT + CSF1Ri-treated WT, *Hexb*^-/-^, BMT-treated *Hexb*^-/-^, and BMT + CSF1Ri-treated *Hexb*^-/-^ mice immunolabeled for GFP (green) and myeloid cell marker IBA1 (red). (j-m) Bar graphs of quantification of cortex images from WT, BMT-treated WT, BMT + CSF1Ri- treated WT, *Hexb*^-/-^, BMT-treated *Hexb*^-/-^, and BMT + CSF1Ri-treated *Hexb*^-/-^ mice of (j) number of IBA1^+^ cells per FOV, (k) mean area covered by filaments of individual IBA1^+^ cells in FOV, (l) mean number of branches per individual IBA1^+^ cell in FOV, and (m) mean cell body volume excluding filaments per IBA1^+^ cell in FOV. Data are represented as mean ± SEM (n=10-11); groups compared by two-way ANOVA with Tukey’s post-hoc test to examine biologically relevant interactions unless otherwise noted; *p < 0.05, **p < 0.01, ***p < 0.001, ****p < 0.0001).

To assess the efficacy of these treatment strategies on functional readouts of disease progression, mice were weighed every other day and motor function was assessed on a weekly basis using the accelerating Rotarod (Fig. 2b-e) task. We observe that *Hexb*^-/-^ mice exhibit a significant change in weight between 13 and 16 weeks of age compared to WT mice (Fig. 2b). Both *Hexb*^-/-^ BMT and *Hexb*^-/-^ BMT + CSF1Ri mice lost significantly less weight by week 16 compared to *Hexb*^-/-^ control mice. On the accelerating Rotarod task, *Hexb*^-/-^ control mice showed a steady decline in motor performance, and all *Hexb*^-/-^ control mice were unable to stay on the Rotarod for any amount of time by week 16 (Fig. 3b, c). Both *Hexb*^-/-^ BMT and *Hexb*^-/-^ BMT + CSF1Ri mice significantly outperformed *Hexb*^-/-^ controls on the Rotarod task following onset of motor deterioration in controls (Fig. 2c-d). However, *Hexb*^-/-^ BMT mice had progressively declining latency-to-fall times over course of testing and had significantly shorter durations than WT BMT control mice in weeks 15 and 16. By contrast, the *Hexb*^-/-^ BMT + CSF1Ri group had stable performance in later weeks and had greater mean differences from *Hexb*^-/-^ controls than *Hexb*^-/-^ BMT mice (Fig. 2c, d). *Hexb*^-/-^ BMT + CSF1Ri mice also did not significantly differ in latency-to-fall time in comparison to WT BMT + CSF1Ri controls at any time point. Additionally, four *Hexb*^-/-^ BMT mice died prematurely or required humane euthanasia at or before week 16, in comparison to three mice in the *Hexb*^-/-^ control group and only one mouse in the *Hexb*^-/-^ BMT + CSF1Ri group. Overall, these data suggest that BMT + CSF1Ri leads to functional rescue, as seen by preservation of motor function and weight normalization in *Hexb*^-/-^ mice.

**Fig. 3:**
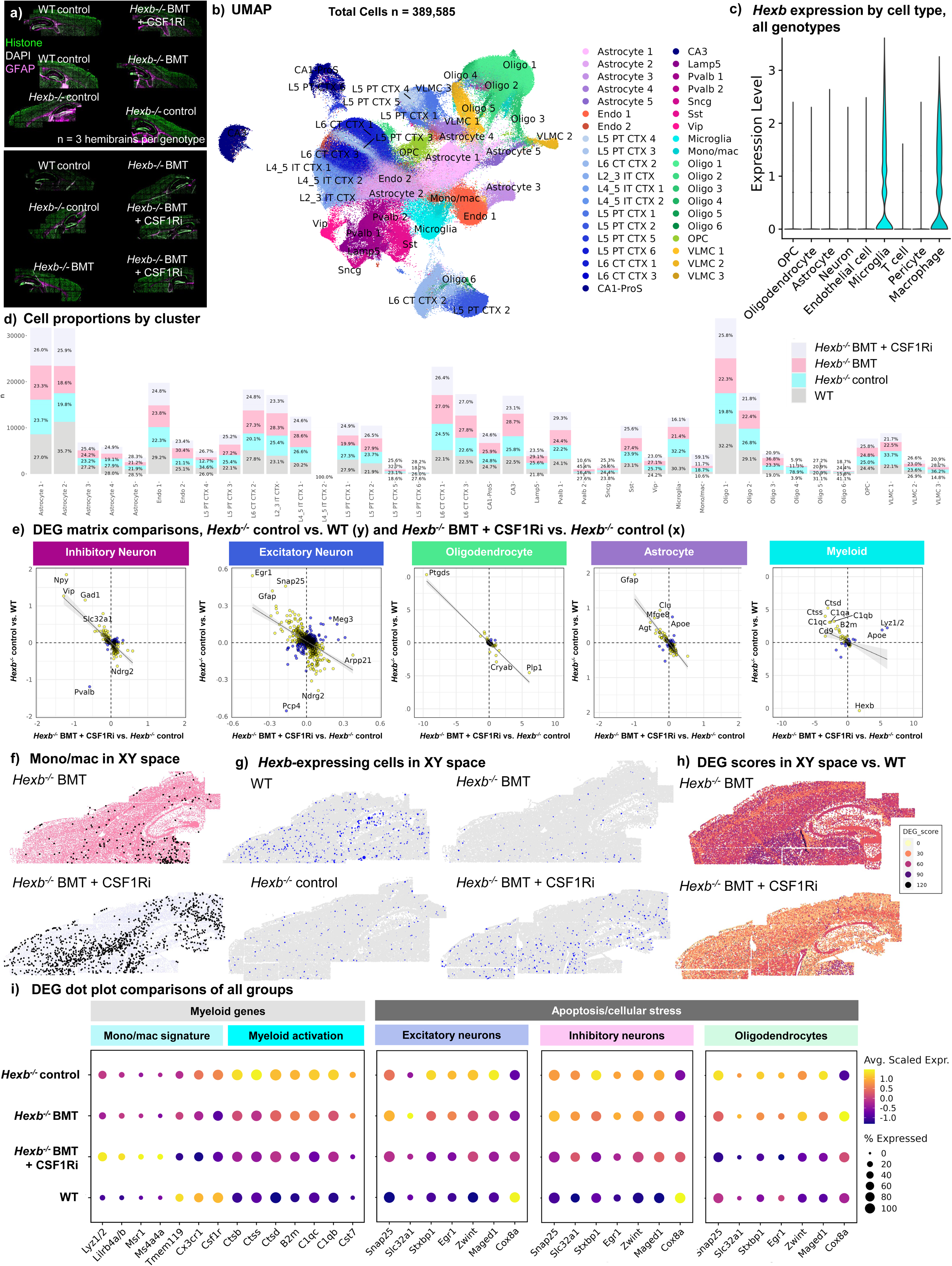
S**p**atial **transcriptomic analysis reveals reversal of disease-associated genetic changes following microglial replacement in SD mice.** (a) Image of WT control, *Hexb*^-/-^ control, bone marrow transplant (BMT)-treated *Hexb*^-/-^, and BMT + colony-stimulating factor 1 receptor inhibitor (CSF1Ri)-treated *Hexb*^-/-^ groups (n=3/group) distributed across 2 slides for spatial transcriptomic analysis, imaged for cell segmentation markers histone (green), DAPI (grey), and GFAP (magenta). (b) Uniform Manifold Approximation and Projection (UMAP) of 389,585 cells across 12 brains. Clustering at 1.0 resolution yielded 38 clusters, which were annotated with a combination of automated and manual approaches with reference to Allen Brain Atlas singe-cell RNA-seq cell types, gene expression, and anatomical location in space. (c) Violin plot of *Hexb* transcript counts in cell types in all cells from all groups, demonstrating myeloid-specific *Hexb* expression. (d) Bar graph of proportions of cell counts by subcluster per group. (e) Comparison matrix scatterplot of the average difference in all significant genes (i.e., padj < 0.05) in inhibitory neurons, excitatory neurons, oligodendrocytes, astrocytes, and myeloid cells between *Hexb*^-/-^ control vs. WT control and BMT + CSF1Ri-treated *Hexb*^-/-^ vs. *Hexb*^-/-^ control. Inversely correlated genes (yellow) occur in opposite directions for each comparison, while directly correlated genes (blue) occur in the same direction for both comparisons. A linear regression line shows the relationship between the two comparisons. (f) The monocyte/macrophage (mono/mac) subcluster (black) overlaid above representative BMT- treated and BMT + CSF1Ri-treated *Hexb^-/-^* brains plotted in XY space. (g) *Hexb-*expressing cells (blue) plotted in XY space in representative brains from WT control, *Hexb*^-/-^ control, BMT-treated *Hexb*^-/-^, and BMT + CSF1Ri-treated *Hexb*^-/-^ mice. Cells were sized in accordance with *Hexb*- expression level, assessed by number of transcripts detected within each cell: cells with 0 transcripts were not plotted, cells with 1 detected transcript were plotted at a point size of 0.001, cells with 2 detected transcripts were plotted at a point size of 0.15, and cells with 3 or more detected transcripts were plotted at a point size of 0.3. (h) Projection of subclusters in XY space colored by DEG score calculated in comparison to WT controls in representative BMT-treated *Hexb^-/-^* and BMT + CSF1Ri-treated *Hexb^-/-^* brains. Following DGE analysis, DEG score was calculated using results of DGE analysis from treatment condition pairs (i.e., BMT-treated *Hexb^-/-^*vs. WT control, BMT + CSF1Ri-treated *Hexb^-/-^* vs. WT control) in each subcluster by summing the absolute value of the log2 fold change values for all DEGs identified between WT control and BMT-treated *Hexb^-/-^* or BMT + CSF1Ri-treated *Hexb^-/-^*with a padj value below 0.05. (i) Dot plot representing pseudo-bulked expression values across the four animal groups (WT control, *Hexb*^-/-^ control, *Hexb*^-/-^ BMT-treated, and *Hexb*^-/-^ BMT + CSF1Ri-treated) in genes related to monocytes/macrophage identity, myeloid cell activation, and apotosis and/or cellular stress in excitatory neurons, inhibitory neurons, and oligodendrocytes.

We next assessed the efficacy of our treatment strategies in inducing BMDM infiltration into the CNS by staining for GFP and IBA1, a marker for myeloid cells. BMT alone led to limited GFP^+^ cell deposition throughout the parenchyma, where the BMT + CSF1Ri group exhibited a broad influx of GFP^+^ cells throughout the parenchyma (Fig. 2f-h). Myeloid cell chimerism in the cortex, identified based on colocalized GFP and IBA1 staining, averaged ∼70-90% in BMT + CSF1Ri mice in comparison to near-zero colocalization after BMT alone (Fig. 2h). These observations are consistent with previous reports of BMT leading to minimal myeloid cell replacement in the parenchyma aside from perivascular and meningeal spaces, contrasting reports of significant infiltration induced by BMT + CSF1Ri^79,80,19^. Overall, these data demonstrate highly efficient and significant replacement of microglia with donor-derived BMDMs following BMT + CSF1Ri. There was no significant difference in replacement rates between any of the *Hexb*^-/-^and WT groups, indicating that loss of *Hexb* does not affect rates of myeloid cell replacement.

To assess whether BMDM engraftment levels in different brain regions coincided with improved motor performance in *Hexb*^-/-^ BMT + CSF1Ri mice, we performed correlation analyses between the final (week 16) Rotarod latency-to-fall time and GFP^+^ cell coverage in the cortex, cerebellum, forebrain, and white matter/corpus callosum. We detected a significant positive correlation between week 16 Rotarod performance and GFP^+^ coverage in the upper corpus callosum (Fig. 2e). This finding complements the region-specific vulnerability identified in the corpus callosum by spatial transcriptomic DGE analysis, as well as previous reports of white matter-specific neurodegeneration in the human SD CNS^76,81^. There was no significant correlation between final week Rotarod performance and total GFP^+^ cell coverage in the cortex, cerebellum, or entire forebrain (Extended Data Fig. 5d-f). These data suggest that moderate overall BMDM infiltration is sufficient to improve motor performance, and the presence of BMDMs in white matter regions is of particular importance. Overall, there is a clear relationship between the broad replacement of microglia with BMDMs and the significant functional rescue observed in the *Hexb*^-/-^ BMT + CSF1Ri group in comparison to *Hexb*^-/-^ controls, which BMT alone was not sufficient to produce.

Detailed profiling of myeloid cell morphology revealed several changes induced by loss of *Hexb* which were effectively reversed following microglial replacement. Staining for IBA1, a myeloid cell marker, revealed various morphological differences in *Hexb^-/-^* cells consistent with microglial activation, including a greater average cell count (Fig. 2j), decreased process (filament) length (Fig. 2k), decreased number of branches per cell (Fig. 2l), and increased cell body volume (Fig. 2m). The *Hexb*^-/-^ BMT group only significantly differed from controls in terms of cell count, with an overall loss of cells. However, the WT BMT group also demonstrated a significant loss of total IBA1^+^ cells, indicative of an irradiation-induced effect. By contrast, myeloid cells in the *Hexb*^-/-^ BMT + CSF1Ri group had significantly longer processes, more branches, and a lower average cell body volume than *Hexb*^-/-^ controls. These data suggest that infiltrating BMDMs induced by BMT + CSF1Ri appear less ameboid/activated compared to microglia in *Hexb*^-/-^ control brains. Ultimately, BMT + CSF1Ri treatment results in a 70-90% replacement of microglial cells with BMDMs, with associated changes in myeloid cell phenotypes expectant of microglial replacement with peripheral cells, indicating successful engraftment and a potential reduction of deleterious microglial activation in SD.

### Microglial replacement reverses genetic changes associated with Hexb deficiency

Having confirmed that microglial replacement via BMT and CSF1Ri leads to functional rescue in *Hexb*^-/-^ mice, we next utilized spatial transcriptomic analysis to examine whether the delivery of *Hexb*-sufficient myeloid cells to the CNS can reverse the SD-associated gene expression changes observed in *Hexb*-deficient mice. Three brains from the WT control group and each *Hexb*^-/-^ group (*Hexb*^-/-^ control, *Hexb*^-/-^ BMT, *Hexb*^-/-^ BMT + CSF1Ri) were sagittally sectioned at 10µm and imaged as described previously (632 total FOVs, ∼53 FOVs per brain section) (Fig. 3a). Here, 389,585 cells were captured with a mean transcript count of ∼800 transcripts per cell. Unbiased cell clustering identified 38 transcriptionally distinct subclusters, and clusters and cell types were annotated as described in Fig. 1 (Fig. 3b, Extended Data Fig. 6a).

We first compared *Hexb* transcript expression in myeloid cells in order to assess whether donor BM cells that engrafted the brains indeed expressed *Hexb*. We detected *Hexb* transcripts in both microglia and macrophage populations in the WT and *Hexb^-/-^* BMT + CSF1Ri mice, confirming the presence of *Hexb* transcripts in donor-derived cells (Fig. 3c, d). Assessing other broad cell types, we detected minimal *Hexb* transcripts in OPCs, oligodendrocytes, astrocytes, neurons, endothelial cells, T cells, and pericytes, as observed previously (Fig. 3c). This data reinforces a myeloid-specific *Hexb* expression pattern and identifies monocytes/macrophages as a cell type that can express *Hexb* within the CNS.

Analysis of cell cluster proportions revealed that the second-largest myeloid subcluster, identified as monocytes/macrophages (Mono/mac) by expression of canonical marker genes (high *Lyz1/2*, *H2-Aa*, *Cd74;* low *Tmem119*)^35–37,82^, was drastically expanded in the *Hexb^-/-^* BMT + CSF1Ri group. Plotting the Mono/mac subcluster in XY space in the *Hexb^-/-^* BMT + CSF1Ri brain showed numerous cells scattered throughout the parenchyma (Fig. 3f), a spatial pattern consistent with the location and distribution of BMDMs, as indicated by GFP staining. Aside from the Mono/mac subcluster and the vascular broad cell type, the number of cells within the broad cell types (Extended Data Fig. 6c) and cellular subclusters (Fig. 3d) were largely consistent between groups. These data indicate that BMT + CSF1Ri treatment induces the infiltration of putative BMDMs that express *Hexb* in the CNS.

To assess whether gene expression changes associated with loss of *Hexb* were reversed with microglial replacement, we performed DGE analysis on all subclusters and broad cell types (Extended Data Figs. 8-10) between experimental group pairs. DGE analyses revealed shifts in gene expression between *Hexb^-/-^* BMT + CSF1Ri and *Hexb^-/-^* control mice in many broad cell types (Extended Data Fig. 8a). To evaluate whether the specific genes altered by Hexb deficiency (either upregulated or downregulated in *Hexb^-/-^* control versus WT mice) were rescued or reversed by BMT and CSF1Ri treatment (comparing *Hexb^-/-^*BMT + CSF1Ri with *Hexb^-/-^*control mice), we generated comparison matrices to assess expression differences in these two pairs (Fig. 3e). We were especially interested in whether certain disease-associated genes would display reversed directionality, i.e., whether genes that were downregulated in *Hexb^-/-^* control mice vs. WT mice would be upregulated in *Hexb^-/-^* BMT + CSF1Ri vs. *Hexb^-/-^*control mice and vice versa. Comparison matrices revealed that many DEGs between *Hexb^-/-^* control and WT mice were significantly changed in the opposite direction between *Hexb^-/-^* BMT + CSF1Ri and *Hexb^-/-^* control mice. In neurons, DEGs with reversed directionality included genes associated with apoptosis and lysosomal dysfunction (*Hspa8, Egr1, Npy, Sgk*), neurodevelopment and synaptic function (*Snap25, Arpp21, Slc32a1, Gad1*), and immunomodulation (*Vip*)^51,61,65,68,73,83–86^. In oligodendrocytes, DEGs involved in apoptosis, cellular stress, and myelination (i.e., *Ptgds, Cryab, and Plp1)*^52–54^ that had previously been identified between *Hexb^-/-^* control and WT mice demonstrated a reversal of expression directionality in *Hexb^-/-^*BMT + CSF1Ri versus *Hexb^-/-^* control mice. In astrocytes, upregulated disease-associated DEGs *Clu, Mfge8,* and *Agt* in *Hexb^-/-^* control mice versus WT were downregulated in *Hexb^-/-^* BMT + CSF1Ri mice in comparison to *Hexb^-/-^* controls. Finally, numerous myeloid subcluster genes that were upregulated in *Hexb^-/-^* control versus WT mice (*Ctsd, Ctss, C1qa, C1qc, B2m, Cd9)* were then downregulated between *Hexb^-/-^* BMT + CSF1Ri and *Hexb^-/-^* control mice following replacement of microglia with BMDMs, indicating that BMT + CSF1Ri reverses disease-associated myeloid cell changes^46–49,87^. Myeloid cells in the *Hexb^-/-^* BMT + CSF1Ri group also had significantly elevated *Hexb* expression compared to *Hexb^-/-^*control mice. The reversal of directionality in disease-associated gene signatures observed with microglial replacement demonstrates the efficacy of this strategy in addressing SD-related phenotypes at the transcript level in glial cells and neurons.

We next sought to compare the efficacy of BMT + CSF1Ri treatment over BMT alone in reversing transcriptomic changes caused by *Hexb* deficiency. In contrast to the reversal in transcriptional changes observed in the BMT + CSF1Ri group, fewer broad cell type DEGs were reversed in directionality and/or were reversed to a lesser degree in terms of log2fold change or adjusted p value in *Hexb^-/-^* BMT mice in comparison to *Hexb^-/-^* controls (Extended Data Fig. 8b). Plotting *Hexb-*expressing cells in XY space showed, predictably, high levels of expression in WT controls with minimal/background *Hexb* expression in *Hexb^-/-^*controls, which appeared unchanged in *Hexb^-/-^* BMT mice (Fig. 3g). By contrast, the *Hexb^-/-^* BMT + CSF1Ri mice demonstrated a restoration of *Hexb-*expressing cells which mirrored the spatial localization of the Mono/mac subcluster. DGE analysis and DEG score calculation revealed higher DEG scores and greater overall deviation from WTs in *Hexb^-/-^* BMT mouse brains; plotting DEG scores in XY space revealed higher overall DEG scores throughout the brain in *Hexb^-/-^* BMT mice than *Hexb^-/-^* BMT + CSF1Ri mice when each group was compared to WT controls (Fig. 3h). By performing a pseudo- bulk analysis in each broad cell type for all four groups (Extended Data Figs. 10b-11e), we confirmated a BMDM signature in the myeloid cell population of the *Hexb^-/-^* BMT + CSF1Ri group only (upregulation of monocyte/macrophage genes *Lyz1/2, Lilrb4a/b, Msr1, Ms4a4a;* downregulation of microglial homeostatic genes *Tmem119, Cx3cr1, Csf1r)* (Fig. 3i). Myeloid activation genes were reduced in both *Hexb^-/-^*BMT groups in comparison to *Hexb^-/-^* controls, though to a greater extent in *Hexb^-/-^* BMT + CSF1Ri mice. Pseudobulk analysis also revealed that genes associated with apoptosis and cellular stress pathways demonstrated reversed directionality in the *Hexb^-/-^*BMT + CSF1Ri group versus *Hexb^-/-^* controls in excitatory neurons, inhibitory neurons, and oligodendrocytes. These genes were largely unchanged in *Hexb^-/-^* BMT groups in comparison to *Hexb^-/-^* controls, demonstrating a failure of BMT alone to reverse genetic indicators of apoptotic processes. These data demonstate similarity between WT mice and *Hexb^-/-^* BMT + CSF1Ri and greater divergence from WT mice in *Hexb^-/-^* BMT mice. Overall, BMT is not sufficient to reverse the majority of gene expression changes associated with loss of *Hexb.* These findings underscore the importance of CSF1Ri-based microglial replacement in correction of disease-associated gene expression changes in neurons, myeloid cells, oligodendrocytes, and astrocytes in *Hexb^-/-^* mice.

In sum, spatial transcriptomic analysis reveals that several disease-associated gene signatures in *Hexb^-/-^* mice can be reversed with microglial replacement following BMT + CSF1Ri treatment. Numerous DEGs identified between *Hexb^-/-^* and WT mice were subsequently reversed in directionality between *Hexb^-/-^* BMT + CSF1Ri-treated mice and *Hexb^-/-^* controls, including genes related to apoptosis, myelination/demyelination, cellular stress response, inflammatory response, and endo-lysosomal function. We also observed a restoration of *Hexb* expression with the introduction of BMDMs to the *Hexb^-/-^*CNS. The ability of microglial replacement to correct SD-associated changes at the molecular level further underscores the potential of this strategy to treat disease.

### Proteomic analysis demonstrates reversal of disease-associated expression changes in neurons and myeloid cells following microglial replacement

To further understand the effects of *Hexb* loss and microglial replacement, we performed spatial proteomic analysis using the CosMx Spatial Molecular Imager (Fig. 4a). We utilized a multi-plex 67-protein mouse neuroscience panel on four 10μm sagittal brain sections from WT control mice and all *Hexb*^-/-^ groups (*Hexb*^-/-^ control, *Hexb*^-/-^ BMT, *Hexb*^-/-^ BMT + CSF1Ri). This technique allows for detailed analysis based on protein markers while maintaining the original structure of the tissue. The panel contains markers relevant to inflammation, lysosomal function, and neurodegenerative disease. A total of 1,199,879 cells were identified and imaged for expression of protein markers. Cell segmentation was automated based on DAPI, histone, and GFAP markers, with clear separation even in densely packed regions such as the dentate gyrus (Fig. 4a). Cells were sorted into subtypes based on marker expression, and plotting in XY space demonstrated accurate identification (Fig. 4b). We identified seven neuronal subsets as well as astrocytes, neuroepithelial cells, microglia, vascular cells, and oligodendrocytes.

**Fig. 4:**
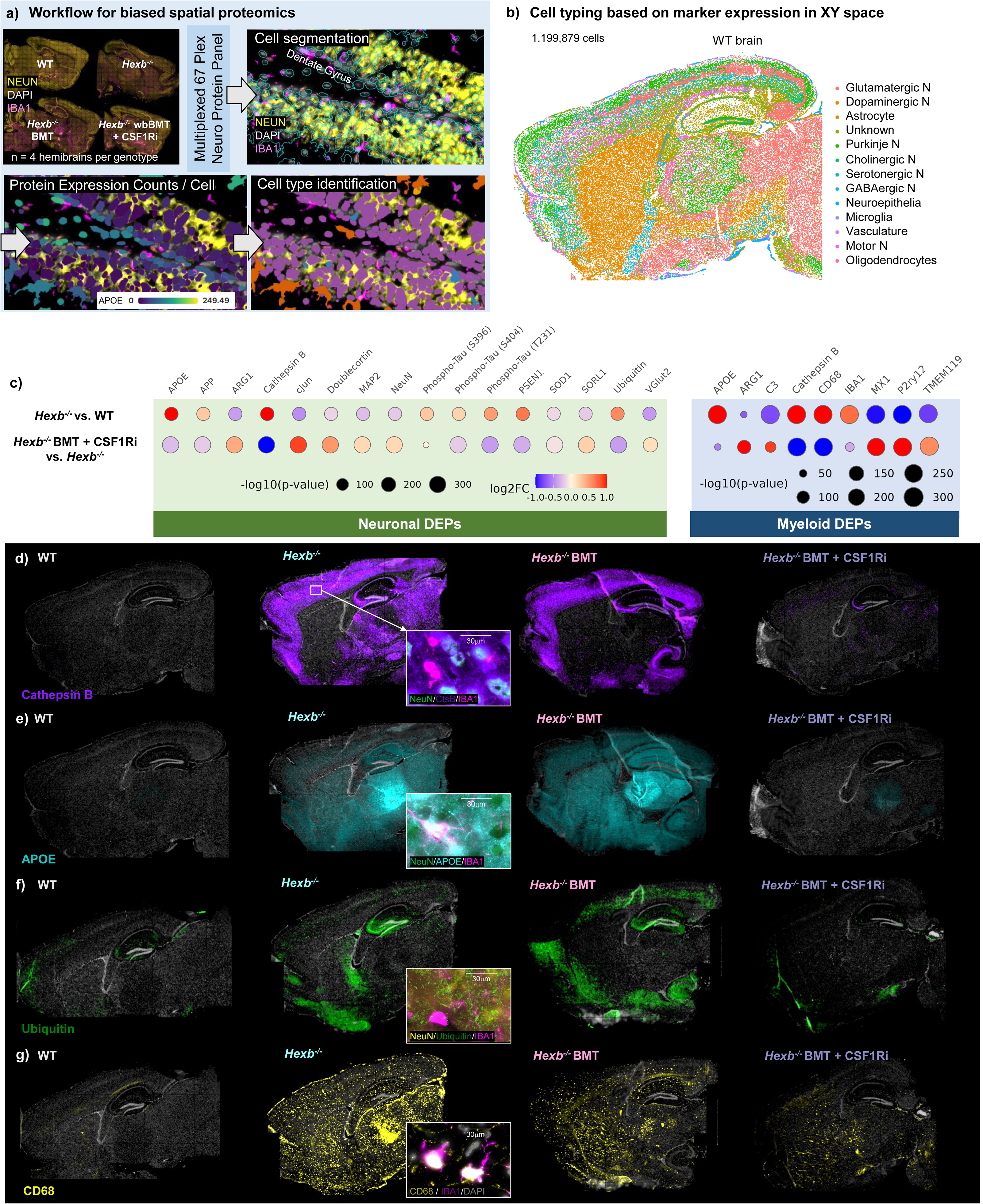
S**p**atial **proteomic analysis identifies disease-associated protein expression patterns in the SD mouse brain which are reversed with microglial replacement.** (a) Workflow for targeted 67-plex single-cell spatial proteomics. Fields-of-view (FOVs) are imaged with cell segmentation markers GFAP, NEUN, RPS6, and IBA1. Protein abundance is determined by quantification of fluorescently-labelled oligos bound to proteins within each cell. Cell types are identified using the CELESTA algorithm, which classifies cells based using expression of marker proteins. (b) Cell types plotted in XY space in representative WT control brain. 1,199,876 cells were captured across the four groups (WT control, *Hexb*^-/-^ control, BMT-treated *Hexb*^-/-^, and BMT + CSF1Ri-treated *Hexb*^-/-^ [n=4/group]). CELESTA cell classification yielded 13 cell types, which were plotted in space to confirm accurate identification. (c) Bubble plots of differentially expressed proteins (DEPs) of interest between pairs *Hexb^-/-^*control vs. WT control, and BMT + CSF1Ri- treated *Hexb^-/-^* vs. *Hexb^-/-^* control in neurons and myeloid cells. Dots are sized by p value (-log10p value) and colored by average difference (log2 fold change, red indicating increased expression, blue indicating decreased expression) of each DEP. (d-g) Representative whole brain images of WT control, *Hexb*^-/-^ control, BMT-treated *Hexb*^-/-^, and BMT + CSF1Ri-treated *Hexb*^-/-^ brains and expanded insets showing cellular marker colocalization of proteins (d) Cathepsin B (purple), colocalization with NeuN^+^ neurons (green) and not IBA1^+^ myeloid cells (magenta); (e) alipoprotein e (APOE, cyan), colocalization with both NeuN^+^ neurons (green) and IBA1^+^ myeloid cells (magenta); (f) Ubiquitin (green), colocalization with NeuN^+^ neurons (yellow) and not IBA1^+^ myeloid cells (magenta); (g) CD68 (yellow), colocalization with IBA1^+^ myeloid cells (magenta) with DAPI (grey) illustrating the rescue of pathological and lysosomal phenotypes by combined BMT and CSF1Ri treatment.

To assess how loss of *Hexb* affects expression of various proteins, especially those associated with lysosomal-endosomal function in the murine brain, we next performed differential protein expression (DPE) analysis between all groups in all cellular subsets (Extended Data Fig. 12b-f). We were particularly interested in differentially expressed proteins (DEPs) in neurons and myeloid cells after identifying disease-associated gene expression signatures in these cell types, and whether protein expression changes between *Hexb*^-/-^ and WT mice were reversed between *Hexb*^-/-^ BMT + CSF1Ri in comparison to *Hexb*^-/-^ controls (Fig. 4c). Neurons and microglia/myeloid cells from *Hexb^-/-^* mice both had significantly higher expression of several proteins associated with dysregulation of the endosomal-lysosomal system compared to WT cells, including APOE and cathepsin B^88–91^. Cathepsin B protein was prominent in NeuN^+^ neurons specifically, and expression was visible throughout the cortex, subiculum, dentate gyrus, pyramidal neurons of the hippocampus, and white matter striations of the thalamus (Fig. 4d). APOE protein was widespread and did not appear to colocalize with any particular cell type (Fig. 4e). Neurons from *Hexb*^-/-^ mice also exhibited elevated expression of several proteins associated with neurodegenerative diseases and/or lysosomal dysfunction in comparison to WT controls, such as amyloid precursor protein (APP), several species of phosphorylated tau, Presenilin 1 (PSEN1), and ubiquitin^88,92–96^. Proteins associated with normal neuronal health and development, such as c-Jun, doublecortin, and MAP2 were downregulated in *Hexb*^-/-^ mice versus WT controls; when dysregulated, many of these proteins have also been associated with apoptosis^97–99^. Microglia/myeloid cells in *Hexb^-/-^* mice exhibited significantly elevated expression of CD68, a lysosomal marker linked to microglial/myeloid cell activation^100^. In line with myeloid cell activation, we observe colocalization of CD68 with IBA1^+^ myeloid cells in *Hexb^-/-^* brains, including in the pia mater layer of the meninges (Fig. 4g). We also observed elevated APOE and CD68 deposition in the thalamus of *Hexb^-/-^* mouse brains. These data are in agreement with the region-specific effects identified by spatial transcriptomic analysis and data from human SD patients^75–77^. Microglia/myeloid cells in the *Hexb^-/-^* brain also showed marked reductions in homeostatic microglial proteins P2RY12 and TMEM119 in comparison to cells from WT control mice^82,101^. These findings provide further insight into the various myeloid and neuronal cell disruptions when myeloid *Hexb* expression is lost, manifesting as lysosomal abnormalities, neuronal dysregulation, and polarization of microglia from homeostatic to activated phenotypes.

We were next interested in whether microglial replacement via BMT + CSF1Ri led to reversal of protein expression changes associated with loss of *Hexb*. Indeed, all DEPs identified in neurons and myeloid cells between *Hexb^-/-^* control mice and WT controls exhibited reversed directionality and significant differences in expression between *Hexb^-/-^* BMT + CSF1Ri and *Hexb^-/-^* controls (Fig. 4c). Visually, expression of notable DEPs cathepsin B, APOE, ubiquitin, and CD68 was partially reduced in *Hexb^-/-^* BMT-treated mice (Fig. 4d-g). In the *Hexb*^-/-^ BMT group, the Cathepsin B phenotype was only corrected in white matter striations in the thalamus (Fig. 4d). By contrast, the overexpression of these proteins was completely or near-completely eliminated in the *Hexb^-/-^* BMT + CSF1Ri group. These data complement the findings from spatial transcriptomic analysis and demonstrate the BMT + CSF1Ri-induced microglial replacement can correct disease-associated protein expression patterns relevant to myeloid activation, lysosomal abnormalities, and neurodegenerative pathways. Furthermore, these data indicate that BMT + CSF1Ri improves upon the partial reductions in disease-associated protein expression achieved by BMT alone.

### CNS pathological changes associated with loss of Hexb are rescued following combined BMT and CSF1Ri treatment

Previous studies have shown that BMT prolongs lifespan and slows functional deterioration in *Hexb*^-/-^ mice, but fails to prevent disease pathology, especially in neurons (i.e., brain glycolipid storage)^11,16,102^. Having identified a reversal of disease-associated gene signatures and protein expression patterns in mice treated with BMT + CSF1Ri, we next sought to investigate the efficacy of combined BMT and CSF1Ri treatment in ameliorating CNS pathological changes in *Hexb*^-/-^ mice. We first performed Periodic Acid Schiff (PAS) staining, a detection method for glycolipids/glycoproteins. In line with prior reports^103^, *Hexb*^-/-^ and BMT- treated *Hexb*^-/-^ mice exhibit numerous PAS^+^ deposits throughout the brain parenchyma, which are consistent with the shape and size of neurons, and absent in WT animals (Fig. 5a-c). We observe a significant reduction in PAS^+^ staining in BMT-treated compared to control *Hexb*^-/-^ mice, indicating that BMT does partially reduce glycolipid storage, but does not resolve this pathology (Fig. 5c). Notably, PAS^+^ deposits were undetectable in BMT + CSF1Ri *Hexb*^-/-^ mice, indicating that replacement of *Hexb*-deficient microglia with *Hexb*-sufficient BMDMs can fully rescue the pathological accumulation of glycolipids in the murine SD brain.

**Fig. 5:**
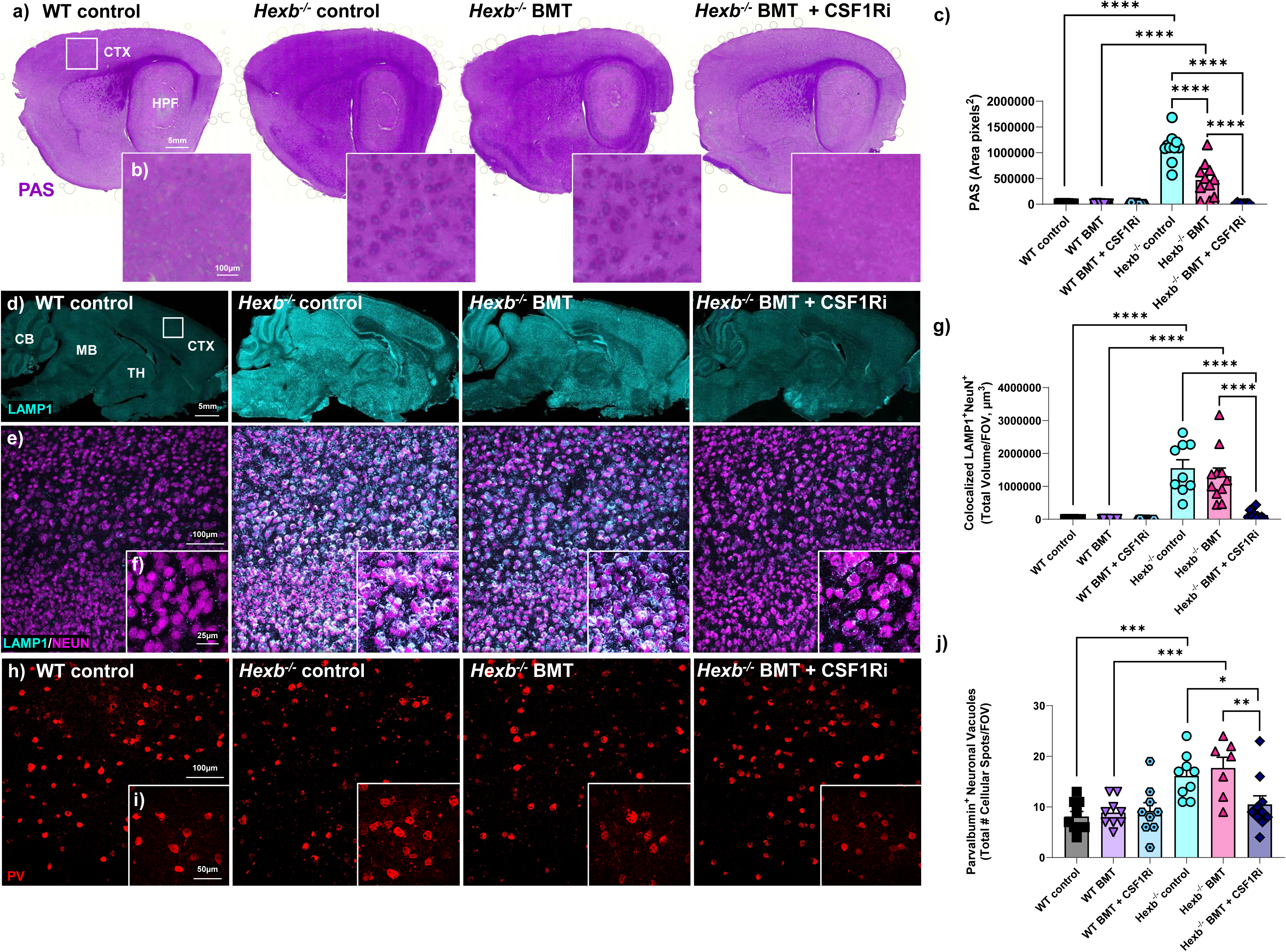
B**r**ain **pathological changes in neurons associated with loss of Hexb are rescued following combined BMT and CSF1Ri treatment.** (a) Representative whole brain sagittal sections and (b) 10x brightfield images of the cortex stained for Periodic acid Schiff (PAS, purple), a method to detect glycolipids, in the brains of wildtype (WT), *Hexb*^-/-^, bone marrow transplant (BMT)-treated *Hexb*^-/-^, and BMT + colony-stimulating factor 1 receptor inhibitor (CSF1Ri)-treated *Hexb*^-/-^ mice. (c) Bar graph of quantification of PAS staining in the cortex of WT, BMT-treated WT, BMT + CSF1Ri-treated WT, *Hexb*^-/-^, BMT-treated *Hexb*^-/-^, and BMT + CSF1Ri-treated *Hexb*^-/-^ mice illustrating the rescue of pathological glycolipid accumulation by combined BMT and CSF1Ri treatment. (d) Representative whole brain images of sagittal sections stained for lysosomal- associated membrane protein 1 (LAMP1, cyan), a marker for lysosomes, in WT, *Hexb*^-/-^, BMT- treated *Hexb*^-/-^, and BMT + CSF1Ri-treated *Hexb*^-/-^ mice. CTX, cortex; HPF, hippocampal formation; CB, cerebellum; MB, midbrain; TH, thalamus. (e) Representative immunofluorescence confocal images of LAMP1 (cyan) and NeuN (magenta), a marker for neurons, staining in the cortex of WT, *Hexb*^-/-^, BMT-treated *Hexb*^-/-^, and BMT + CSF1Ri-treated *Hexb*^-/-^ mice. Insert (f) represents a higher resolution confocal image highlighting the co-localization (white) of LAMP1^+^ staining within NeuN^+^ neurons in *Hexb*^-/-^ and BMT-treated *Hexb*^-/-^ mice. (g) Bar graph of quantification of co-localized LAMP1^+^ and NeuN^+^ staining in confocal images of the cortex of WT, BMT-treated WT, BMT + CSF1Ri-treated WT, *Hexb*^-/-^, BMT-treated *Hexb*^-/-^, and BMT + CSF1Ri- treated *Hexb*^-/-^ mice. (h) Representative immunofluorescence confocal images of the cortex in WT, *Hexb*^-/-^, BMT-treated *Hexb*^-/-^, and BMT + CSF1Ri-treated *Hexb*^-/-^ mice stained for parvalbumin (PV, red). Inset (i) represents a higher resolution confocal images within cortex in WT, *Hexb*^-/-^, BMT-treated *Hexb*^-/-^ and BMT + CSF1Ri-treated *Hexb*^-/-^ mice showing the presence of enlarged holes or vacuoles within PV^+^ cells in the cortex of *Hexb*^-/-^ and BMT-treated *Hexb*^-/-^ brains. (j) Bar graph of quantification of vacuoles within PV^+^ neurons in confocal images of cortex of WT, BMT- treated WT, BMT + CSF1Ri-treated WT, *Hexb*^-/-^, BMT-treated *Hexb*^-/-^, and BMT + CSF1Ri-treated *Hexb*^-/-^ mice. Data are represented as mean ± SEM (n=10-11; groups compared by two-way ANOVA with Tukey post hoc testing; *p < 0.05, ** p < 0.01, *** p < 0.001, ****p < 0.0001).

Next, we assessed the effects of *Hexb* deficiency, BMT, and microglial replacement on lysosomal alterations in neurons by co-staining for LAMP1, a marker for lysosomes and autophagic organelles, and NeuN, a marker for neurons. Immunostaining revealed extensive LAMP1^+^ accumulation (Fig. 5d) that colocalized with neurons (Fig. 5e-f) in *Hexb*^-/-^ mouse brains, indicative of a disruption in the endosomal-lysosomal system in murine SD; in line with this, previous studies have shown that LAMP1 is degraded by Hexβ^37–41^. Here, we show that BMT alone did not significantly reduce LAMP1^+^ staining in the *Hexb*^-/-^ brain. However, BMT + CSF1Ri treatment led to a drastic and significant reduction in LAMP1^+^ staining *Hexb*^-/-^, demonstrating near- (Fig. 5d-f). These findings provide evidence that microglial replacement can resolve abnormal lysosomal phenotypes within neurons in *Hexb*^-/-^ mice.

Having identified that several inhibitory neuronal subsets are affected by *Hexb* deficiency during spatial transcriptomics analysis, we next screened for morphological abnormalities and cell loss in parvalbumin (PV) neurons, a marker for the largest inhibitory neuronal subtype, in *Hexb*^-/-^ mice. We did not observe a significant loss in the number of cortical NeuN^+^ or PV^+^ cells in *Hexb*^-/-^ mice, but we did detect neuronal abnormalities: PV^+^ neurons in the *Hexb*^-/-^ control group demonstrated unusual puncta within the cell body, indicative of vacuolization (Fig. 5h,i). These abundant vacuoles were consistent with previous reports in SD and have previously been identified as enlarged, dysfunctional lysosomes^23,104,105^. Vacuoles were also present in the *Hexb*^-/-^ BMT group, but significantly reduced in the *Hexb*^-/-^ BMT + CSF1Ri group, which did not differ from WT BMT + CSF1Ri controls (Fig. 5j). Altogether, we demonstrate correction of several neuronal pathologies and abnormalities with microglial replacement in the SD CNS, reiterating the therapeutic potential of this treatment paradigm over traditional BMT approaches and suggesting that infiltrating *Hexb-*sufficient BM-derived myeloid cells can improve neuronal pathology in SD.

### BMT alleviates pathological hallmarks outside of the CNS

As SD is not limited to the CNS, we next profiled the consequences of our treatments outside of the CNS to assess the total-body efficacy of *Hexb*-sufficient myeloid cell replacement. We first performed histological analysis of the liver, an organ which exhibits high accumulation of GM2 gangliosides and other glycolipids in SD. Staining for GFP identified prominent deposition of donor bone marrow-derived GFP^+^ cells in the livers of all BMT groups (Fig. 6a, b). There were no significant differences in GFP^+^ cell counts between BMT alone and BMT + CSF1Ri in either WT or *Hexb^-/-^* mice. Having shown that liver myeloid cells were replaced following BMT treatment, we next stained for LAMP1 to assess endosomal-lysosomal abnormalities. Here, we observed a significant increase in LAMP1^+^ staining in the livers of *Hexb*^-/-^ control mice compared to WT mice, as in the CNS, which was abolished in both the *Hexb*^-/-^ BMT and *Hexb*^-/-^ BMT + CSF1Ri groups (Fig. 6c, d). Finally, we performed a PAS stain and found PAS^+^ deposits throughout the liver parenchyma in *Hexb*^-/-^ control animals, which were eliminated in both BMT and BMT + CSF1Ri groups (Fig. 6e, f). Together, these findings indicate that BMT alone is sufficient to improve pathological hallmarks in the SD liver, in alignment with previous reports^102^.

**Fig. 6:**
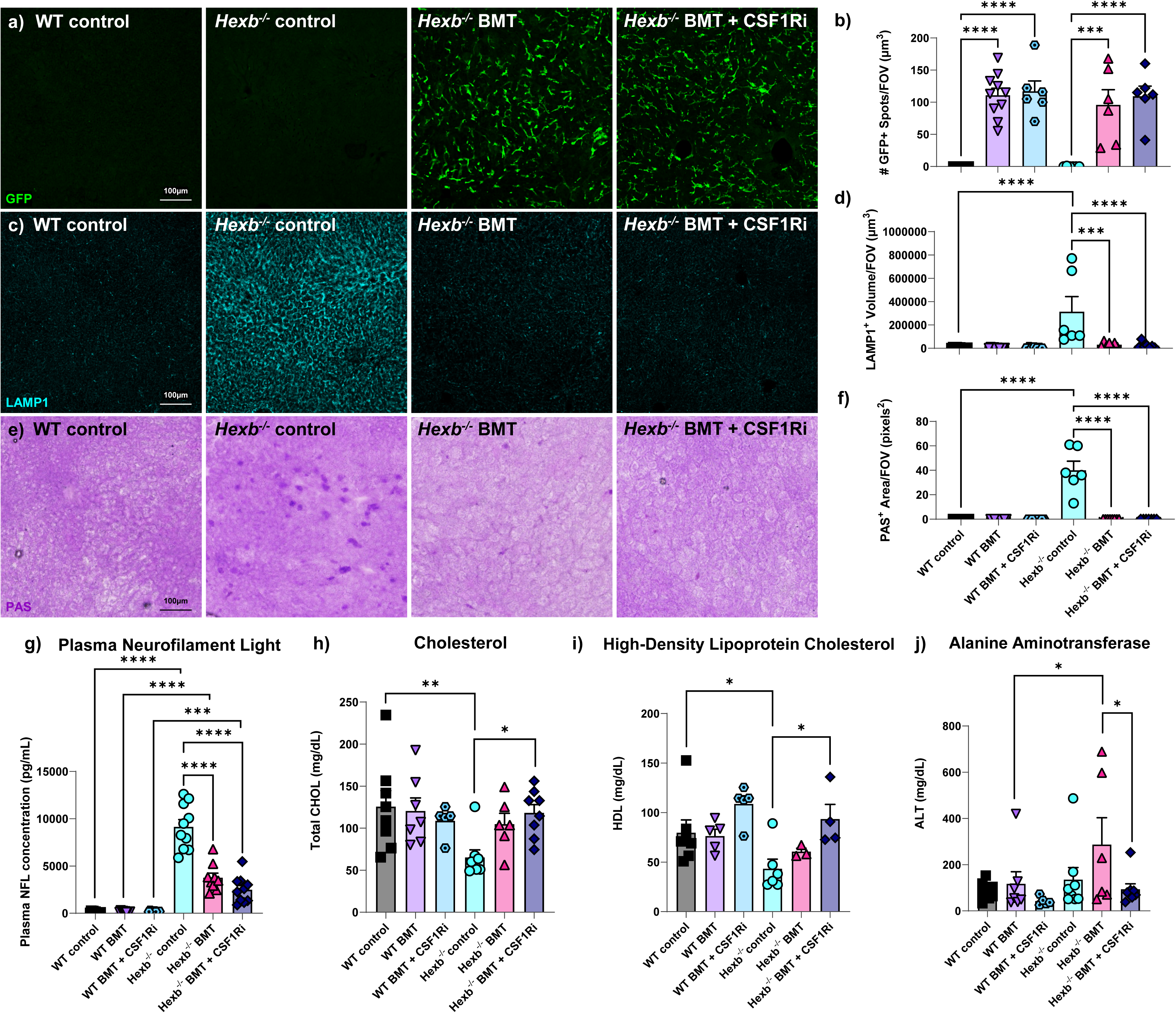
P**e**ripheral **changes associated with loss of Hexb are rescued with BMT.** (a) Representative confocal images of liver sections from WT, *Hexb*^-/-^, bone marrow transplant (BMT)-treated *Hexb*^-/-^, and BMT + colony-stimulating factor 1 receptor inhibitor (CSF1Ri)-treated mice immunolabeled for green fluorescent protein (GFP, green). (b) Bar graph of quantification of GFP^+^ cells (spots) in liver images from WT, BMT-treated WT, BMT + CSF1Ri-treated WT, *Hexb*^-/-^, BMT-treated *Hexb*^-/-^, and BMT + CSF1Ri-treated *Hexb*^-/-^ mice showing engraftment of cells derived from CAG-EGFP bone marrow donors with BMT treatment. (c) Representative confocal images of liver sections from WT, *Hexb*^-/-^, bone BMT-treated *Hexb*^-/-^, and BMT + CSF1Ri-treated *Hexb*^-/-^ mice immunolabeled for lysosomal-associated membrane protein 1 (LAMP1, cyan). d) Bar graph of quantification of LAMP1^+^ volume in liver images from WT, BMT-treated WT, BMT + CSF1Ri-treated WT, *Hexb*^-/-^, BMT-treated *Hexb*^-/-^, and BMT + CSF1Ri-treated *Hexb*^-/-^ mice. (e) Expanded and cropped whole-liver brightfield images stained for Periodic acid Schiff (PAS, purple), a method to detect glycolipids, in WT, *Hexb*^-/-^, bone BMT-treated *Hexb*^-/-^, and BMT + CSF1Ri-treated *Hexb*^-/-^ mice. (f) Bar graph of quantification of PAS staining in imagrs of the liver of WT, BMT-treated WT, BMT + CSF1Ri-treated WT, *Hexb*^-/-^, BMT-treated *Hexb*^-/-^, and BMT + CSF1Ri-treated *Hexb*^-/-^ mice illustrating the rescue of pathological glycolipid accumulation by BMT. (g) Measurement of plasma neurofilament light (NfL) in WT, BMT-treated WT, BMT + CSF1Ri-treated WT, *Hexb*^-/-^, BMT-treated *Hexb*^-/-^, and BMT + CSF1Ri-treated *Hexb*^-/-^ mice. (h) Measurement of total plasma cholesterol (CHOL) concentration in WT, BMT-treated WT, BMT + CSF1Ri-treated WT, *Hexb*^-/-^, BMT-treated *Hexb*^-/-^, and BMT + CSF1Ri-treated *Hexb*^-/-^ mice. (i) Measurement of total plasma cholesterol (CHOL) concentration in WT, BMT-treated WT, BMT + CSF1Ri-treated WT, *Hexb*^-/-^, BMT-treated *Hexb*^-/-^, and BMT + CSF1Ri-treated *Hexb*^-/-^ mice. Data are represented as mean ± SEM (n=6-8, livers; n=5-11, plasma); groups compared by two-way ANOVA with Tukey’s post-hoc test to examine biologically relevant interactions unless otherwise noted; *p < 0.05, **p < 0.01, ***p < 0.001, ****p < 0.0001). HDL measurement was unable to be obtained in some samples due to high heme content in plasma.

In addition to immunohistochemical analysis of the liver, we also collected blood plasma to assess the levels of neurofilament light chain (NfL), a well-established biomarker of neurodegeneration that correlates with axonal damage^106^. Previous studies have shown that NfL is increased in human SD patients^107^. Here, we demonstrate that *Hexb*^-/-^ control mice display significanctly higher concentrations of plasma NfL than WT mice, signifying axonal damage (Fig. 6g). Notably, NfL was significantly reduced in both *Hexb*^-/-^ BMT and *Hexb*^-/-^ BMT + CSF1Ri-treated mice compared to *Hexb^-/-^*mice, indicative of a reduction in axonal damage in both treatment contexts. Piccolo multi-chemistry analysis of plasma also demonstrated a significant alteration in several circulating lipids/enzymes. In comparison to WT controls, plasma from *Hexb*^-/-^ control mice exhibited significantly lower concentrations of total cholesterol (Fig. 6h) and high density lipoprotein (HDL) cholesterol (Fig. 6i), often referred to as “good” cholesterol. Both cholesterol abnormalities were ameliorated with BMT + CSF1Ri treatment in *Hexb*^-/-^ BMT + CSF1Ri mice compared to *Hexb*^-/-^ control mice (Fig. 6h,i). Interestingly, we observed a significant elevation in the concentration of alanine aminotransferase (ALT), a liver enzyme which increases in blood plasma following acute liver injury^108^, in *Hexb*^-/-^ BMT mice in comparison to WT BMT mice; this was significantly reduced in *Hexb*^-/-^ BMT + CSF1Ri mice (Fig. 6j). This elevation in ALT was not present in any other groups. Overall, we report significant normalization in the concentrations of several plasma biomarkers of disease with BMT-based treatments in *Hexb*^-/-^ mice. These results highlight the benefits of a total-body intervention in SD, rather than a CNS-specific treatment strategy which would not address SD-related pathology in other organ systems.

### Myeloid-derived Hexβ is restored with microglial replacement and is secreted from microglia in a Ca^2+^dependent, P2X7-mediated manner and subsequently taken up by neurons

Following confirmation that BMDMs engrafted in the murine SD CNS are able to resolve substrate accumulation and lysosomal abnormalities within neurons, we were interested in whether Hexβ activity was restored the brains of treated mice. While many lysosomal enzymes are only active within the lysosome, previous studies have indicated that Hexβ is enzymatically active outside of the it, including in the extracellular space^109–111^. We therefore utilized a Hexβ activity assay^112^ to asses enzyme activity in two protein fractions acquired from frozen brain hemispheres from WT control, *Hexb*^-/-^ control, *Hexb*^-/-^ BMT, and *Hexb*^-/-^ BMT-treated mice. We first homogenized pulverized brains in a high-salt, detergent-free buffer and collected supernatant to extract salt-soluble proteins while minimizing cell lysis. Typically, efficient dissolution of the cell membrane requires a detergent^113^; therefore, the salt-soluble fraction is likely enriched for extracellular proteins. We then resuspended the pellet in a detergent-containing buffer to lyse cells and extract total protein from the tissue. We detected Hexβ in both fractions in WT mice (Fig. 7b, c). Upon assessing activity in each *Hexb*^-/-^ group, we found minimal activity in both fractions from *Hexb*^-/-^ control brains, with no significant increase in *Hexb*^-/-^ BMT-treated brains in either fraction. We also did not observe a significant difference between *Hexb*^-/-^ control mice and *Hexb*^-^

**Fig. 7:**
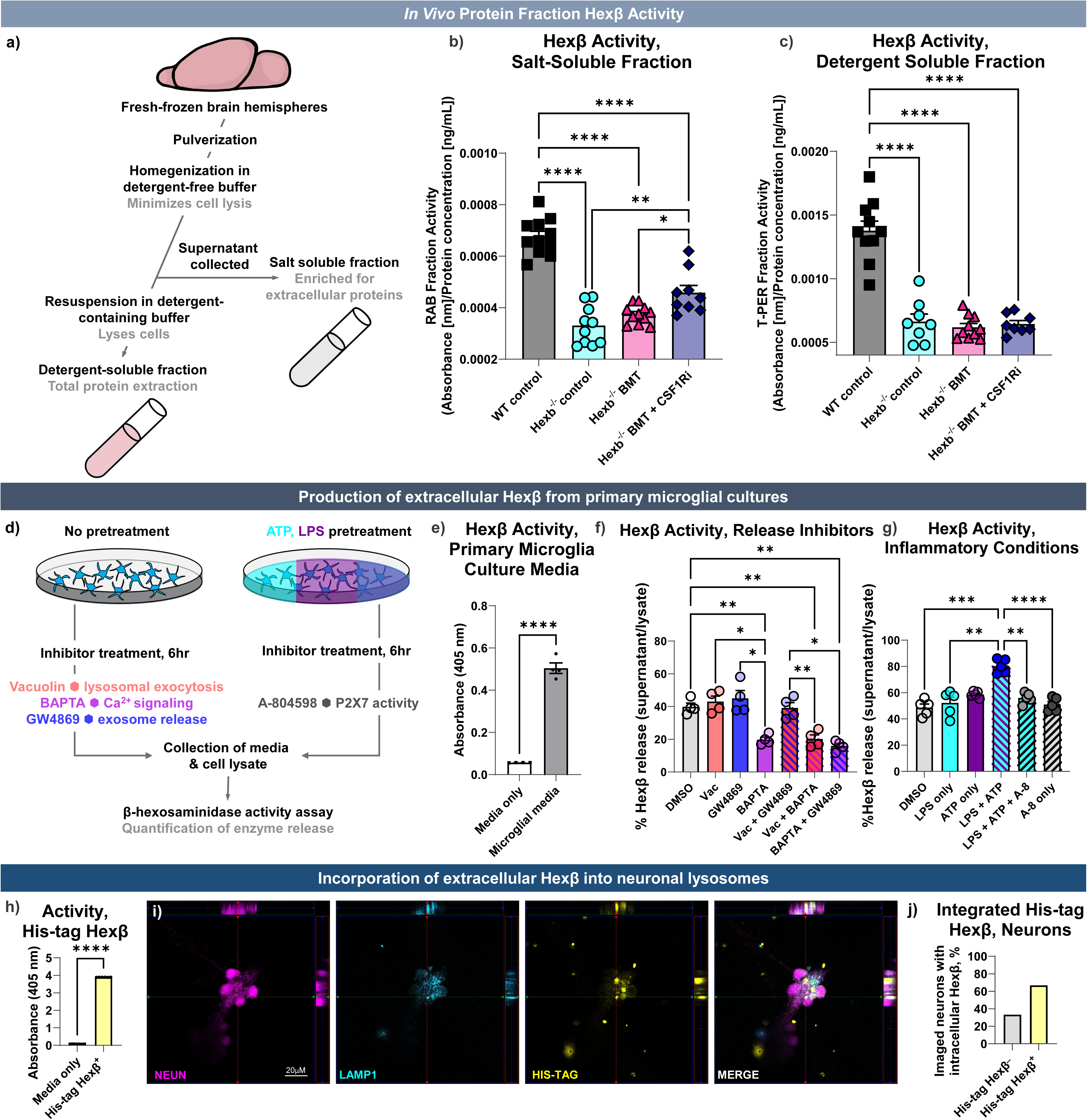
Hexβ is restored in an exracellular-enriched brain protein fraction in *Hexb^-/-^* mice treated microglial replacement, and is secreted by microglia and taken up by neurons *in vitro* (a) Schematic of protein fraction collection. Pulverized fresh-frozen hemispheres from WT, *Hexb*^- /-^, bone marrow transplant (BMT)-treated *Hexb*^-/-^, and BMT + colony-stimulating factor 1 receptor inhibitor (CSF1Ri)-treated mice were homogenized in a high-salt, detergent-free buffer to limit cell lysis and enrich for extracellular proteins. Supernatant was collected as the salt-soluble fraction. The pellet was then resuspended in a detergent-containing buffer to lyse cells and supernatant was collected as the detergent-soluble fraction. (b) Bar graph of absorbance values from β- hexosaminidase (Hexβ) enzymatic activity assay normalized to protein concentration in reassembly buffer (RAB) salt-solube protein fraction in WT, *Hexb*^-/-^, bone BMT-treated *Hexb*^-/-^, and BMT + CSF1Ri-treated *Hexb*^-/-^ mice. (c) Bar graph of Hexβ activity normalized to protein concentration in Total Protein Extraction Reagent (T-PER) buffer detergent-solube protein fraction in WT, *Hexb*^-/-^, bone BMT-treated *Hexb*^-/-^, and BMT + CSF1Ri-treated *Hexb*^-/-^ mice. (d) Schematic of *in vitro* primary microglial experiments. For inhibitor experiments, cultured primary microglia derived from mouse neonates were incubated with inhibitors of lysosomal exocytosis (Vacuolin), calcium (Ca^2+^) signaling (BAPTA), or lysosomal exocytosis (GW4869) for 6 hours. For lipopolysaccharide (LPS) and adenoside triphosphate (ATP) experiments, cells were primed with LPS for 3 hours, incubated with an inhibitor of the P2X7 purinergic receptor (A-804598) for 10 minutes, and treated with ATP for 20 minutes. Hexβ activity in media and cell lysate was then assed using a Hexβ enzymatic activity assay. (e) Hexβ activity assay measured by absorbance value in culture media only and culture media collected from primary microglial cultures demonstrating *in vitro* secretion of Hexβ from microglia. Groups compaired using an unpaired Student’s T test. (f) Bar graph of Hexβ release measured as a ratio of Hexβ activity in supernatant (cell culture media) normalized to Hexβ activity in cell lysate in cultured primary microglia treated with dimethyl sulfoxide (DMSO, control), vacuolin, GW869, BAPTA, vacuolin + GW869, vacuolin + BAPTA, or BAPTA + GW869. (g) Bar graph of Hexβ release measured as a ratio of Hexβ activity in supernatant (cell culture media) normalized to Hexβ activity in cell lysate in cultured primary microglia treated with DMSO (control), LPS, ATP, LPS + ATP, LPS + ATP + A-804598, or A- 804598 alone. (h) Bar graph Hexβ activity assay measured by absorbance value in media only and media containing his-tagged recombinant Hexβ protein, demonstrating that the his-tagged Hexβ protein is enzymatically active. Groups compaired using an unpaired Student’s T test. (i) Confocal images of mouse hippocampal neurons treated with media containing his-tagged Hexβ protein immunolabeled for neurons (NeuN, magenta), lysosomal-associated membrane protein 1 (LAMP1, cyan), his-tagged Hexβ protein (HIS-TAG, yellow), and a merged image showing orthogonal x/z and z/y projections at top and right of image showing colocalization of LAMP1^+^ and HIS-TAG^+^ stainging within NeuN^+^ neurons (white). (j) Bar graph representing the percentage of neurons with intracellular incorporation of his-tagged Hexβ protein as identified by orthogonal imaging of HIS-TAG staining within NeuN^+^ neurons. Shows percentage of imaged neurons without intracellular his-tagged Hexβ staining and neurons with intracellular his-tagged Hexβ staining. Data are represented as mean ± SEM (n=10-11, protein fractions; n=4-5, *in vitro* activity assay; n=12-13, neuronal cultures); groups compared by two-way ANOVA with Tukey’s post-hoc test to examine biologically relevant interactions unless otherwise noted; *p < 0.05, **p < 0.01, ***p < 0.001, ****p < 0.0001).

^/-^ BMT + CSF1Ri mice in the detergent-soluble fraction (Fig. 7c). However, in the salt-soluble fraction, we observed significantly increased Hexβ activity in the *Hexb*^-/-^ BMT + CSF1Ri group in comparison to both *Hexb*^-/-^ control and *Hexb*^-/-^ BMT groups. These data indicate that microglial replacement in *Hexb*^-/-^ mice partially restores Hexβ activity in the salt-soluble, extracellular enriched fraction. This finding further highlights the potential efficacy of a microglial replacement approach for treating SD in reconstituting enzyme in the CNS, while also demonstrating that full enzyme reconstitution to WT levels is not necessary to correct pathological hallmarks.

Given the restoration of activity in an extracellularly-enriched protein fraction and the ability of engrafted BMDMs to correct a litany of disease-associated neuronal phenotypes, we theorized that myeloid cells may be supporting neuronal function through secretion of Hexβ. Thus, we first sought to identify whether microglia release enzymatically active Hexβ protein *in vitro*. To accomplish this, cortices of 3 to 5-day old neonatal wildtype pups were collected, dissociated, and incubated for 14 days to generate a primary cell culture of mixed glial cells. Following this, microglia were isolated via gentle shaking and plated for 48 hours before collection of the supernatant and cell lysate (Fig. 7d). Using the Hexβ activity assay, we found that Hexβ activity was present in the supernatant media collected from primary microglial cultures, which was not present in media alone (Fig. 7e). This result indicates that microglia passively secrete enzymatically active Hexβ under homeostatic conditions *in vitro*.

To further explore the mechanism of Hexβ release from myeloid cells, we investigated several of the main pathways of lysosomal enzyme secretion, including lysosomal exocytosis, exosome release, and calcium-mediated intracelluar pathways^114,115^. We incubated primary microglial cultures with inhibitors of each pathway (vacuolin, lysosomal exocytosis; GW4869, exosome release; BAPTA-AM, calcium signalling)^116–118^ for 6 hours and assessed Hexβ activity (Fig. 7f). Vacuolin and GW4869-treated microglia did not exhibit significantly reduced Hexβ release in comparison to control cells, nor did cells treated with both vacuolin and GW4869. These data indicate that microglial Hexβ release *in vitro* is not driven by lysosomal exocytosis or exosome release. However, we found that treatment with BAPTA-AM significantly reduced the release of active Hexβ by microglia compared to controls, with BAPTA-AM-treated microglia exhibiting a >50% reduction in activity. Combined treatment with BAPTA and vacuolin or GW4869 did not further reduce Hexβ release in comparison to BAPTA alone. These findings suggest that Hexβ is passively secreted by microglia in a calcium-dependent manner independent of lysosomal exocytosis or exosome release.

Considering that inflammatory and/or pathological conditions increase the secretion of other lysosomal enzymes, we next hypothesized that inflammation-mimicking conditions would elicit increased release of Hexβ in cultured microglia^119^. To simulate inflammatory conditions, we incubated cells with lipopolysaccharide (LPS), adenosine triphosphate (ATP), or a combination of both. LPS is frequently used to induce acute inflammation both *in vitro* and *in vivo;* it activates immune cells via activation of toll-like receptor 4 (TLR4), inducing release of inflammatory cytokines^120^. ATP accumulates in the extracellular space in inflammatory conditions, is released by damaged and/or dying cells, and can act as a damage-associated molecular pattern to induce an inflammatory response^121^. Neither LPS or ATP-treated microglia exhibited increased Hexβ release in comparison to untreated control cells (Fig. 7g). However, cells incubated with a combination of both ATP and LPS demonstrated significantly higher levels of Hexβ release than control and LPS-treated cells. These data suggest that the combination of LPS priming subsequent and exposure to ATP, which mimicks physiological inflammatory conditions, is important for the increased release of Hexβ from microglia; this is consistent with previous reports regarding other lysosomal enzymes^121–123^.

A key mediator of inflammation in microglia is the ATP-sensitive P2X7 purinergic receptor, which acts as a scavenger receptor in microglial phagocytosis in the absence of stimulation^124^. Activation of P2X7 by ATP and more potent analogs causes the influx of calcium and leads to microglial activation, cytokine release, and lysosomal destabilization/leakage^125–130^. Given the efficacy of calcium-chelating BAPTA-AM in blunting Hexβ release, we theorized that increased Hexβ secretion induced by ATP + LPS treatment may be mediated by the P2X7 receptor. To test this hypothesis, we primed microglia with LPS for 3 hours, and then pre-treated cultured microglia with P2X7 inhibitor A-804598 for 10 minutes before adding exogeneous ATP for 20 minutes. As predicted, P2X7 inhibition significantly reduced Hexβ release in comparison to cells treated with LPS + ATP alone (Fig. 7g). Hexβ release from cells treated with LPS + ATP + A-804598 did not significantly differ from that of control cells. A-804598 alone without ATP or LPS did not decrease Hexβ release in comparison to untreated control cells. These data indicate that the increased release of Hexβ by microglia following inflammation-mimicking LPS + ATP treatment is mediated by the P2X7 receptor, but secretion of Hexβ under homeostatic/non-inflammatory conditions is P2X7 independent.

Having established that microglia secrete enzymatically active Hexβ via Ca^2+^- and P2X7- mediated mechanisms, we were next interested in the capacity of wildtype neurons to take up Hexβ from the extracellular space. To accomplish this, we acquired his-tagged recombinant mouse Hexβ protein and first confirmed its enzymatic activity using the Hexβ activity assay to assure physiological relevance (Fig. 7h). Dissociated E18 hippocampal neurons were then plated and cultured for one week, before incubation in media containing 10μg of his-tagged Hexβ for 24 hours. Neurons were fixed and subsequently stained with NeuN, His-tag, and LAMP1 antibodies to identify neurons, Hexβ protein, and lysosomal membranes, respectively. Using confocal imaging, we observed that NeuN^+^ neurons incubated with his-tagged Hexβ showed integration of the protein into the cell bodies of ∼67% of total neurons imaged (Fig. 7i, j; 12-13 neurons/neuron clusters imaged per treatment condition). The integration of his-tagged Hexβ *in vitro* indicates that neurons have the capacity to take up extracellular Hexβ. Additionally, we observed colocalized staining between LAMP1 and anti-his within NeuN^+^ cell bodies (Fig. 7i). This colocalization indicates that following uptake into neurons, the Hexβ protein is localized to the lysosomal compartment. These data suggest that neurons are capable of taking up extracellular Hexβ and integrating it into the lysosome *in vitro.* Taken together, our findings provide evidence that myeloid-derived Hexβ can be taken up into neurons, identifying a potential mechanism for the neuronal correction which is observed following microglial replacement in the SD CNS.

## DISCUSSION

Our data demonstrates robust correction of SD phenotypes in *Hexb*^-/-^ model mice following BMT-based microglial replacement. We show that microglia and CNS-engrafted macrophages/monocytes are the only cell types that express *Hexb* in the wildtype CNS, then demonstrate that replacement of *Hexb-*deficient microglia with *Hexb-*sufficient BMDMs leads to the normalization of myeloid cell morphology, reversal of disease-associated changes in gene and protein expression, clearance of enzymatic substrate pathology and lysosomal abnormalities in the brain. We also provide *in vitro* data which identifies the capacity of 1. microglia to release Hexβ and 2. neurons to take up Hexβ and integrate it into the lysosomal compartment. Taken together, these data reveal a critical role for myeloid-derived Hexβ in SD and, by extension, normal neuronal function. While our data and previous studies demonstrate that BMT alone is not sufficient to correct pathological hallmarks of SD in the CNS, our CSF1Ri-mediated approach to induce broad peripheral cell engraftment drastically improves outcomes following BMT.

In addition to demonstrating correction of the murine SD CNS, we also provide the first single-cell spatial transcriptomic and proteomic datasets from brain sections at an advanced disease stage in the *Hexb*^-/-^ mouse model of SD. We found that *Hexb*^-/-^ mice display a strong disease-associated gene expression signature in myeloid cells, oligodendrocytes, astrocytes, and neurons, with the latter demonstrating activation of apoptotic pathways and perturbations in neurotransmission and cellular metabolism. Many of the noted differentially expressed genes aligned with genes previously reported in RNA sequencing dataset derived from human SD and TSD patients, underscoring the strength of the model in recapitulating human disease. We also identified region-specific vulnerabilities which are highly consistent with reports from human SD patients, namely in the thlamaus, white matter tracts, and cortex. Notably, loss of *Hexb* was associated with marked gene expression changes in the myeloid cell population, which was also the only cell type found to express *Hexb* in wildtype brains.

While the mechanism by which ganglioside accumulation causes neurodegeneration is not fully understood, it is clear that microglial activation and peripheral macrophage infiltration are important aspects of the pathogenesis of SD. These myeloid phenotypes precede neurodegeneration in *Hexb*^-/-^ mice^11^, and deletion of neuroinflammatory factors such as tumor necrosis factor-α (TNF-α) and macrophage-inflammatory protein 1α (MIP-1 α/CCL3) has been shown to reduce neurodegeneration and slightly extend the lifespan of *Hexb*^-/-^ mice without reducing neuronal GM2 ganglioside burden^39,131^. These observations have led to speculation that microglial dysregulation caused by loss of *Hexb* and subsequent activation is the driving factor in disease progression in SD. However, evidence from a number of experimental treatment inquiries suggests otherwise. In one study utilizing *Hexb*^-/-^ mice, conditional expression of human Hexβ protein exclusively in neurons extended lifespan and substantially improved neuropathology in *Hexb*^-/-^ mice, despite no reduction in microglial or astrocyte activation^132^. This finding suggests that the restoration of neuronal Hexβ is of greater importance than reducing inflammation in ameliorating disease phenotypes. Studies utilizing viral-vector based gene therapy to treat disease in various animal models of SD have demonstrated similar efficacy, even though they have shown mixed results in terms of reducing microglial activation. The viral vector-induced expression of *Hexb* in neurons in *Hexb*^-/-^ mice, sheep, primates, and felines has been highly effective in reducing pathology and extending lifespan; notably, these viruses do not infect microglia^105,133–138^. Finally, recent work in *Hexb*^-/-^ mice and other LSD model mice indicates that a neuron-intrinsic mechanism drives cell death and disease progression as a result of lysosomal dysfunction; this pathway does not depend on microglial involvement^9^. As a whole, these findings suggest that microglial activation/neuroinflammation alone is not sufficient to explain the neuronal pathology and neurodegeneration observed in SD. In fact, it is clear that Hexβ protein in neurons plays an important role in maintaining cellular health and lysosomal function, despite a confirmed lack of *Hexb* transcript expression in neurons themselves. It is possible that the critical relationship between neurons and microglia in SD is one of enzyme provision rather than inflammation.

Our data strongly supports a relationship between myeloid-derived Hexβ and regulation/restoration of neuronal health. The exclusive expression of *Hexb* transcripts in microglia and BMDMs in our spatial transcriptomic datasets corroborates previous findings identifying it as a myeloid-specific gene. The engraftment of wild type donor-derived BMDMs in the CNS corrected numerous neuron-specific disease signatures identified in *Hexb*^-/-^ mice, including expression of genes related to cellular stress and apoptosis, accumulation of enzymatic substrate, and phenotypes associated with lysosomal dysfunction. In the context of results from other treatment modalities and previous reports regarding *Hexb* expression, our findings suggest that myeloid cells are likely the souce of functional Hexβ in the homeostatic CNS, which is necessary for neuronal health and lysosome function.

We next sought to determine if—and how—cultured microglia secrete Hexβ. Recent work has revealed that CNS myeloid cells can secrete lysosomal enzymes in a manner which affects neuronal health^139,140^. In line with this, our *in vitro* data demonstrate that microglia also secrete enzymatically active Hexβ, building upon previous reports from cultured murine microglia and human-derived monocytes^109,110,114^. Our experiments also show that neurons are capable of taking up extracellular Hexβ protein and integrating it into the lysosomal compartment. Inhbitors of lysosomal exocytosis and exosome release did not reduce *in vitro* Hexβ secretion, but treatment with BAPTA-AM, a calcium chelator, lowered Hexβ secretion by ∼50%. Under homeostatic conditions, the calcium-dependent secretion of Hexβ is likely to be a result of escape from the *trans*-Golgi network. In this network, lysosomal enzymes, including Hexβ, are sorted to the lysosome by the mannose-6 phosphate (M6P) pathway, but up to 40% of enzyme escapes and is instead secreted into the extracellular space^12,141,142^. Secreted enzyme can then be taken up by surrounding cells via M6P receptors expressed on the cell surface^143,144^. In this study, we also show that inflammation-mimicking conditions (LPS + ATP) increase Hexβ secretion, which is abolished by inhibition of the purinergic receptor P2X7, implicating this receptor in increased enzyme secretion under pathological conditions. These *in vitro* experiments provide insight into the potential mechanism(s) by which myeloid cell Hexβ release plays a role in neuronal function and, by extension, how neuronal lysosomal abnormalities may be corrected following BMT + CSF1Ri in *Hexb*^-/-^ mice. Further research is needed to identify the exact secretory pathway in both contexts and confirm relevance *in vivo*.

Our BMT + CSF1Ri approach offers several advantages over other therapeutic interventions, including substrate reduction, enzyme replacement and gene therapies. Previous attempts at artificially rebalancing enzyme-substrate concentrations to treat disease have had mixed results. Therapies directed at reducing enzymatic substrate, though effective in other LSDs, only resulted in a partial delay of disease progression in SD model mice, and minimal human patient improvement^145–147^. Moreover, enzyme replacement therapy is limited by difficulties in accessing the CNS and lack of feasible delivery routes^148,149^. Gene therapy, another promising avenue for the long-term treatment of SD, has also had major drawbacks in *Hexb*^-/-^ animal models and other disease contexts. A primate study which achieved successful Hexβ reconstitution in the CNS unfortunately also reported heavy neurotoxicity, and gene therapy as a whole is presently limited by safety concerns, immunogenicity, and difficulty in accessing the CNS^150–152^. Our study highlights that CNS-engrafted *Hexb-*expressing cells have the potential to reconstitute enzyme in a sustained, physiologically relevant manner and provide long term reduction of substrate. We accomplish this by combining BMT with CSF1R inhibition, which has potential for immediate clinical translation. While BMT once carried significant risk, it has seen major advances in safety and efficacy over recent decades and dramatic increases in long-term survival such that it is now considered the gold standard in various conditions^12,153,154^. However, there are still challenges to utilizing BMT. Perhaps the largest barrier in using BMT to treat CNS conditions is the limited access to the brain parenchyma and failure to correct the CNS. In the present study, by following myeloablative conditioning (irradiation) with CSF1Ri treatment and withdrawal, we are able to overcome this barrier and induce the broad influx of BM-derived cells into the CNS in a mouse model of SD. Importantly, CSF1R inhibitors are also already an approved class of drug for the treatment of Tenosynovial giant cell tumor, further emphasizing the translatability of this strategy^155^.

Though BMDMs perform many of the same immunological functions as microglia, they are not a perfect substitute. Microglia are highly specialized to the environment and demands of the CNS, and BMDMs maintain a distinct transcriptional and phenotypic identity when engrafted in the brain^156–159^. Infiltration of activated monocytes/macrophages in pathological contexts also has deleterious effects^19,160–162^. However, the consequences of having BMDMs engrafted in the CNS are secondary to the potential benefits in a context as severe as SD, especially with no available treatment and other experimental treatments limited by safety concerns. An optimal approach may involve a combination of BMT with administration of induced pluripotent stem cell (iPSC)-derived microglia, a growing field of inquiry^163–166^; however, administration of iPSC-derived microglia alone is unlikely to address the periphery, an important consideration in SD as demonstrated by our observation of correctible liver pathology. Head irradiation also comes with several drawbacks including cognitive and synaptic deficits and microglial activation, though CSF1Ri-mediated microglial depletion has been shown to ameliorate these effects^19,167–169^. Alternative myeloablative conditioning regimes such as busulfan treatment are also compatible with CSF1Ri to induce BMDM infiltration^30–32^; however, these approaches carry their own drawbacks, especially in pediatric administration, including neurotoxicity and other neurocognitive effects^170^. Future advances in the safety and tolerability of BMT will make this approach more widely applicable in pateints. Overall, our approach harnesses a commonly utilized clinical practice in BMT and the innate properties of myeloid cells to deliver Hexβ to correct the SD CNS, with BMDMs replacing microglia as the putative cellular source of Hexβ . Further research and refinement of this approach to mitigate the present limitations will improve its viability and enhance a strategy that could be applied in other neurodegenerative LSDs and a litany of additional CNS conditions in the future.

## ACKNOWLEDGMENTS

This work was supported by the National Institutes of Health (NIH) under awards: R01NS083801 (NINDS), R01 AG081599 (NIA), RF1AG056768 (NIA), and U54AG054349 (NIA Model Organism Development and Evaluation for Late-onset Alzheimer’s Disease [MODEL-AD]) to K.N.G, T32NS082174 (NINDS) to K.I.T, and T32NS121727 (NINDS) to N.K. We acknowledge the support of the Flow Cytometry Core in the Sue & Bill Gross Stem Cell Research Center.

## METHODS

### Compounds

PLX5622 was provided by Plexxikon Inc. and formulated in AIN-76A standard chow by Research Diets Inc. at 1200 ppm.

### Animals

All animal experiments were performed in accordance with animal protocols approved by the Institutional Animal Care and Use Committee (IACUC) at the University of California, Irvine, an American Association for Accreditation of Laboratory Animal Care (AAALAC)-accredited institution, and were conducted in compliance with all relevant ethical regulations for animal testing and research.

*Mice:* All mice were obtained from The Jackson Laboratory. We utilized B6;129S-*Hexb^tm1Rlp^*/J mice in this study, which harbor a loss-of-function mutation in the *Hexb* gene, described in detail previously^7^ (strain #002914). Heterozygous breeding pairs were used to generate *Hexb*^-/-^ mice and WT littermate controls. For BMT, bone marrow cells were isolated from sex-matched CAG-EGFP donor mice (strain #006567). Animals were housed in autoclaved individual ventilated cages (SuperMouse 750, Lab Products, Seaford, DE) containing autoclaved corncob bedding (Envigo 7092BK 1/8” Teklad, Placentia, CA) and two autoclaved 2” square cotton nestlets (Ancare, Bellmore, NY) plus a LifeSpan multi-level environmental enrichment platform. Ad libitum acces to water (acidified to pH2.5–3.0 with HCl then autoclaved) and food (LabDiet Mouse Irr 6F; LabDiet, St. Louis, MO) was provided. Cages were changed every 2 weeks with a maximum of five animals per cage. Room temperature was maintained at 72 ± 2°F with ambient room humidity (average 40–60% RH, range 10 – 70%). Light cycle was 12h light / 12h dark, with lights on at 06:30h and off at 18:30h. Animals were assigned to treatment groups by randomization and was balanced for sex. Experimenters were blinded to genotype and treatment group during behavioral testing and analysis of histological data.

*Genotyping:* Genotyping for the *Hexb* mutation was performed using two primer sets to amplify both the wildtype (Forward 5’-ATT TTA AAA TTC AGG CCT CGA-3’ and Reverse 5’-CAT TCT GCA GCG GTG CAC GGC) and *Hexb* mutant (5’-CAT AGC GTT GGC TAC CCG TGA-3’) sequences using cycling conditions from The Jackson Laboratory and a JumpStart taq antibody (mouse stock #002914, JumpStart A7721-200TST MilliporeSigma, Burlington, MA).

### Animal treatments

*Bone marrow transplant*: 4-6 week-old mice were irradiated with 9 Gy (XRAD 320 irradiator, Precision X-ray, North Branford, CT), anesthetized with isoflurane (5% induction and 2% maintenance isoflurane, vol/vol) and reconstituted via retroorbital injection with 2 × 10^6^ whole bone marrow cells isolated from CAG-EGFP donor mice in 50μL of sterile saline solution. The irradiator was equipped with a hardening filter (0.75mm Sn + 0.25mm Cu + 1.5mm Al; HVL = 3.7mm Cu, half value layer) to eliminate low energy X-rays. X-ray irradiation was delivered at a rate of 1.10 Gy/min. Following transplant, mice were transferred to sterile cages with autoclaved bedding and water, supplemented with DietGel® (76A formulation, one half cup/cage at point of cage transfer, ClearH2O, INC., Westbrook, ME) and fed with Uniprim® antibiotic supplement diet (Envigo Bioproducts, Madison, WI) for 14 days to support recovery and prevent opportunistic infection.

*Microglial depletion*: Following a two week recovery period, mice were fed ad libitum with PLX5622 at a dosage of 1200 ppm (to eliminate microglia) or vehicle (control) for 7 days. Mice were then returned to vivarium diet.

### Behavioral monitoring

*Rotarod task:* Motor function was monitored on a weekly basis from 11-16 weeks of age using an accelerating Rotarod (Ugo Basile, Gemonio, Italy). Each mouse was placed on the rotarod beam while it was stationary, then acceleration was initiatied. The Rotarod apparatus automatically tracked the duration between initiation of acceleration and mouse falling to the base of the apparatus. A total of five consecutive trials were performed per mouse each week and trial times were averaged for each mouse. If a mouse was unable to maintain position on the stationary beam and fell to the base prior to initiation of acceleration for a trial, a score of zero was manually entered.

*Weight monitoring*: Mice were weighed every other day starting at 13 weeks of age until the point of sacrifice to assess condition and progression to humane endpoint defined as a loss of 20% of original body weight. If mice reached humane endpoint prior to 16 weeks of age, mice were sacrificed and tissue was collected as described below. Mice which did not reach 16 weeks of age were not included in the final dataset or analyses.

### Tissue preparation for histology

Mice were euthanized by CO2 inhalation at 16 weeks of age or upon reaching the humane endpoint, depending on which was reached first. Mice were then transcardially perfused with 1X phosphate buffered saline (PBS). Brain hemispheres were divided along the midline, and the left lobe of the liver was cut in half. One hemisphere of each brain and one half of each liver were fresh-frozen on dry ice and stored at -80°C, while the other hemisphere and liver half were fixed in 4% paraformaldehyde (PFA) in PBS (Thermo Fisher Scientific, Waltham, MA) for 48 hr at 4°C for immunohistochemical analysis, then cryoprotected in 30% sucrose + 0.05% sodium azide. PFA-fixed brain halves were then embedded in optimal cutting temperature media (OCT; Tissue-Tek, Sakura Fintek, Torrance, CA) and sectioned into either 10 μm or 35μm sagittal slices using a cryostat (CM1950, LeicaBiosystems, Deer Park, IL). 10μm sections were mounted directly on slides for immunohistochemistry. 35μm sections were washed three times with fresh 1X phosphate-buffered saline (PBS) to remove excess OCT before transferring to a 1x PBS + 30% glycerol + 30% ethylene glycol solution for storage at -20°C. PFA-fixed liver halves were sectioned into 35μm slices using a Leica SM2000R freezing microtome and sections were stored in 1x PBS + 30% glycerol + 30% ethylene glycol at -20°C. Brains and livers were protected from light to maintain GFP fluorescence.

### Flow cytometry

At the time of sacrifice, bone marrow and whole blood were harvested and analyzed by flow cytometry for hematopoietic stem cell and granulocyte chimerism. Bone marrow/hematopoietic stem cells were extracted from femurs and tibia by flushing with ice cold PBS. Whole blood/granulocytes were collected in EDTA via cardiac puncture following CO2 euthanasia. Samples were centrifuged at 1250rpm for 5 minutes. Supernatant was discarded, then samples were incubated with 1 mL of 1x ACK Lysing Buffer (A1049201, Gibco, Waltham, MA) for 1 minute at RT, protected from light. Reaction was quenched with 9 mL of ice cold PBS, and cells were again centrifuged at 1250rpm for 5 minutes. Supernatant was discarded, and pellet was resuspended in 1mL PBS. Finally, samples were centrifuged at 2400rpm for 5 minutes, supernatant was discarded, and pellet was reconstituted in 225µL of PBS. Cells were then stained for flow cytometric analysis as previously described^19^ with the following surface antibodies purchased from Biolegend (San Diego) and diluted in PBS at 1:200 unless otherwise noted: CD34-eFlour660 (1:50, #50-0341-80, eBioscience), Sca-1-AF700 (1:100, #108141), Ter119-PE/Cy5 (#116209), ckit/CD117-PE/Cy7 (#25-1171-81, eBioscience), CD150/SLAM-PerCP-eFlour710 (#46-1502-82, eBioscience), CD11b-APC (#101212), Gr1-AF700 (#108422), CD45-APC/Cy7 (#103116), NK1.1-PE (#108707), and CD27-APC/Cy7 (#124226). Flow cytometry analysis was performed using a BD LSRFortessa X20 Benchtop Flow Cytometer (BD Biosciences, Franklin Lakes, NJ), and data was analyzed in BD FACSDiva and FCS Express software.

### Plasma lipid and neurofilament light chain measurement

Blood plasma was collected at time of sacrifice and analyzed using the Piccolo® blood chemistry analyzer from Abaxis (Union City, CA) following manufacturer instructions. Plasma samples were thawed from -80°C one at a time and diluted 1:1 with distilled water (ddH2O), then 100 μl of diluted sample was loaded onto a Piccolo lipid plus panel plate (#07P0212, Abaxis). Various lipid parameters including total cholesterol (CHOL), high-density lipoprotein (HDL), non-HDL cholesterol (nHDLc), triglycerides, low-density lipoprotein (LDL), and very low-density lipoprotein (vLDL) were analyzed and plotted. Some parameters could not able to be assessed in samples with high heme content. Lipid and general chemistry controls (#07P0401, Abaxis) were utilized to verify accuracy and reproducibility of the measurements. Quantitative biochemical analysis of plasma neurofilament light chain (NfL) was performed with the R-Plex Human Neurofilament L Assay (K1517XR-2; Meso Scale Discovery).

### Histology

*Immunohistochemistry:* Fluorescent immunohistochemical labelling followed a standard indirect technique as described previously^171^. Briefly: free-floating sections underwent a series of washes at room temperature in 1x PBS three times in for 10 minutes, 5 minutes, and 5 minutes. Sections were then immersed in blocking serum solution (5% normal goat serum with 0.2% Triton X-100 in 1X PBS) for 1 hour at room temperature, followed by overnight incubation at 4°C in primary antibodies diluted to concentrations described below with blocking solution. Finally, sections were incubated, covered, with fluorescent secondary antibodies for 1hr, followed by 3 washes in 1X PBS prior to mounting on microscope slides and coverslipping with Fluoromount-G with or without DAPI (0100–20 and 0100–01; SouthernBiotech, Birmingham, AL).

Brain and liver sections were stained with combinations of antibodies against ionized calcium-binding adapter molecule 1 (IBA1, 1:1000; #019–19741, Wako, Osaka, Japan), green fluorescent protein (GFP, 1:200; ab13970, Abcam, Waltham, MA), neuronal nuclei (NeuN, 1:1000; Ab104225; Abcam), lysosome-associated membrane protein 1 (LAMP1, 1:200; Ab25245, Abcam), and parvalbumin (Pvalb, 1:500; MAB1572, Millipore, Burlington, MA). Whole brain and liver images were captured with a Zeiss Axio Scan Z1 Slidescanner using a 10× 0.45 NA Plan-Apo objective. High resolution fluorescent images of brain sections were captured using a Leica TCS SPE-II confocal microscope and LAS-X software. One 20X Z-stack (2 μm step interval, within a depth of 35-40 μm) field-of-view (FOV) per brain region was captured per mouse, and max projections of 63X Z-stacks were used for representative images where indicated. Liver sections were imaged using a Zeiss LSM 900 Airyscan 2 microscope and Zen image acquisition software (Zen Blue, Carl Zeiss, White Plains, NY). All images were collected using a 20x / NA 0.8 lens, and one Z-stack image (within a depth of 35-40 μm) per mouse/sex/group was acquired in each liver. Airyscan processing of all channels and z-stack images was performed in Zen software and Bitplane Imaris Software was used for quantification of 20x confocal images.

*Periodic Acid Schiff:* Free-floating brain and liver sections underwent three 1x PBS washes as described above. Sections were there placed in a 0.5% periodic acid solution diluted in millipure water and incubated for 5 minutes. Sections were then briefly washed in ddH2O 3 times for 1.5 minutes each. Sections were then incubated for 15 minutes in Schiff’s reagent (3952016, MilliporeSigma, Burlington, MA) and washed in tap water 4 times for 1 minute and 15 seconds each and washed once briefly in ddH2O. Sections were mounted and coverslipped as above. 10x brightfield images were captured with an Olympus BX60F5 microscope (Hachioji-shi, Tokyo, Japan) with an attached Nikon camera (DS-Fi3; Shinagawa-ku, Tokyo, Japan) using NIS-Elements D 5.30.05 64-bit. ImageJ analysis software was used for quantification of brightfield images.

### Image analysis

*Imaris:* Confocal images were quantified using the spots and surfaces modules in Imaris v9.7 software. Volumetric measurements (i.e., GFP^+^ staining volume, IBA1^+^ microglia volume) were acquired automatically utilizing the surfaces module in confocal images of livers and cortex, corpus callosum, and cerebellum brain regions. Quantitative comparisons between experimental groups were always carried out in simultaneously stained sections. For whole-brain GFP quantification, the spots algorithm was used to automatically quantify GFP^+^ cells in a defined region of identical size for each brain that included the midbrain and forbrain, but not the hindbrain. For microglial morphology, microglial branching and filament area was assessed using the filaments module.

*ImageJ:* Brightfield images were converted to 8-bit gray-scale and quantified using Fiji ImageJ. The threshold feature was adjusted and used to distinguish signal from background before percent coverage was measured. Standardized limits for deposit size and circularity were applied to each image to further distinguish signal from background. To quantify PAS percent area coverage, the whole FOV of each 10x cortical image was analyzed and PAS^+^ deposits identified by thresholding were reported in pixel coverage over the total image.

### Data analysis and statistics

Statistical analysis was performed with GraphPad Prism software (v.10.0.1). To compare two groups, the unpaired Student’s t test was used. To compare unpaired groups, a one-way ANOVA with Tukey’s post hoc test was used. To compare paired groups with repeated measures for identical subjects over time (i.e. weekly Rotarod testing), a repeated measures ANOVA with Tukey’s post-hoc test was used. To compare condition-paired groups, a two-way ANOVA with Tukey’s post hoc test (3 groups, 2 conditions) or Sidak’s test (2 groups, 2 conditions) was used. For all analyses, statistical significance was defined by a p value below 0.05. All bar graphs are represented as group mean ± SEM, significance is expressed as follows: *p < 0.05, **p < 0.01, ***p < 0.001, ****p<0.0001. n represents the number of mice within each group.

### Spatial transcriptomic & proteomic analysis

*Section preparation:* One day prior to experiment, PFA-fixed brain hemispheres were embedded in optimal cutting temperature (OCT) compound (Tissue-Tek, Sakura Fintek, Torrance, CA), and 10um sagittal sections were cut using a cryostat (CM1950, LeicaBiosystems, Deer Park, IL). Six hemibrains were mounted onto VWR Superfrost Plus slides (Avantor, 48311–703) and kept at −80°C overnight. For *Hexb*^-/-^ BMT groups and the WT control group, n=3 mice per experimental condition were utilized (wild-type control, *Hexb*^-/-^ control, *Hexb*^-/-^ BMT, *Hexb*^-/-^ BMT + CSF1Ri) for transcriptomics and proteomics. When selecting representative brains, we considered BMDM infiltration levels from both *Hexb*^-/-^ BMT groups, choosing brains with similar total forebrain GFP^+^ staining to group averages. Tissue was processed in accordance with the Nanostring CosMx fresh-frozen slide preparation manual for RNA and protein assays (NanoString University).

*Slide treatment, RNA, day 1:* Slides were removed from -80°C and baked at 60°C for 30 min. Slides were then processed for CosMx: three 1X washes PBS for 5 minutes each, 4% sodium dodecyl sulfate (SDS; CAT#AM9822) for 2 minutes, three 1X PBS washes for 5 minutes each, 50% ethanol for 5 minutes, 70% ethanol for 5 minutes, and two washes with 100% ethanol for 5 minutes each before allowing slides to air dry for 10 minutes at room temperature. Antigen retrieval was performed using a pressure cooker maintained at 100°C for 15 min in preheated 1X CosMx Target Retrieval Solution (Nanostring, Seattle, WA). Slides were then transferred to DEPC-treated water (CAT#AM9922) and washed for 15 seconds, incubated in 100% ethanol for 3 minutes, and air dried at room temperature for 30 minute. Slides were incubated with digestion buffer (3 μg/mL Proteinase K in 1X PBS; Nanostring) for tissue permeabilization, then washed 2 times in 1X PBS for 5 minutes. Fiducials for imaging were diluted to 0.00015% in 2X SSC-T and incubated on slides for 5 minutes. Following fiducial treatment, slides were protected from light at all times. Tissues were then post-fixed with 10% neutral buffered formalin (NBF; CAT#15740) for 1 minute, washed twice with NBF Stop Buffer (0.1M Tris-Glycine Buffer, CAT#15740) for 5 minutes each, and washed with 1x PBS for 5 minutes. Next, NHS-Acetate (100 mM; CAT#26777) mixture was applied to each slide and incubated at room temperature for 15 minutes. Slides were washed twice with 2X SSC for 5 minutes each. Slides were then incubated for 16–18 hours in a hybridization oven at 37°C with a modified 1000-plex Mouse Neuroscience RNA panel (Nanostring) for *in situ* hybridization with the addition of an rRNA segmentation marker.

*Slide treatment, RNA, day 2:* Following *in situ* hybridization, slides were washed twice in pre- heated stringent wash solution (50% deionized formamide [CAT#AM9342], 2X saline-sodium citrate [SSC; CAT#AM9763]) at 37°C for 25 minutes each, then washed twice in 2X SSC for 2 minutes each. Slides were then incubated with DAPI nuclear stain for 15 minutes, washed with 1X PBS for 5 minutes, incubated with GFAP and histone cell segmentation markers for 1 hour, and washed three times in 1X PBS for 5 minutes each. Flow cells were adhered to each slide to create a fluidic chamber for spatial imaging. Slides were loaded into and processed automatically with the CosMx instrument. Approximately 300 fields of view (FOVs) were selected on each slide, capturing hippocampal, corpus callosum, upper thalamic, upper caudate, and and cortical regions for each section. Slides were imaged for approximated 7 days and data were automatically uploaded to the Nanostring AtoMx online platform. Pipeline pre-processed data was exported as a Seurat object for analysis with R 4.3.1 software.

*Side treatment, protein, day 1:* Slides were removed from -80°C and baked at 60°C for 30 min, then washed three times with 1X Tris Buffered Saline with Tween (TBS-T; CAT#J77500.K2) for 5 minutes each. Antigen retrieval was performed using a pressure cooker held at 80°C in pre-heated Tris-EDTA buffer (10 mM Tris Base [CAT#10708976001], 1 mM EDTA solution, 0.05% Tween 20, pH 9.0) for 7 minutes. Following antigen retrieval, slides were allowed to cool to room temperature for 5 minutes, then washed three times in 1X TBS-T for 5 minutes each. Slides were incubated with Buffer W (Nanostring) for 1 hour at room temperature. Slides were then incubated for 16-18 hours at 4°C with the CosMx 64-plex protein panel and segmentation markers (GFAP, IBA1, NEUN, and S6).

*Side treatment, protein, day 2:* Following incubation, slides were washed three times with 1X TBS- T for 10 minutes each, then washed with 1X PBS for 2 minutes. Fiducials for imaging were diluted to 0.00005% in 1X TBS-T and incubated on the slide for 5 minutes. Slides were the washed with 1X PBS for 5 minutes, incubated in 4% PFA for 15 minutes, and washed three times with 1X PBS for 5 minutes each. Slides were incubated with DAPI nuclear stain for 10 minutes, then washed twice wuth 1X PBS for 5 minutes each. Slides were then incubated with 100 mM NHS-Acetate for 15 minutes and washed with 1X PBS for 5 minutes. Flow cells were adhered to each slide to create a fluidic chamber for spatial imaging. Slides were loaded into and processed automatically with the CosMx instrument. Approximately 600 FOVs were selected per slide, capturing each full section. Slides were imaged for ∼6 days data were automatically uploaded to the Nanostring AtoMx online platform. Pipeline pre-processed data was exported as a Seurat object for analysis with R 4.3.1 software.

*Spatial transcriptomics data analysis:* Spatial transcriptomics datasets were processed as previousy described^172^. Principal component analysis (PCA) and uniform manifold approximation and projection (UMAP) analysis were performed to reduce the dimensionality of the dataset and visualize clusters in space. Unsupervised clustering at 1.0 resolution yielded 39 clusters for the WT control versus *Hexb*^-/-^ control dataset and 38 clusters for the dataset which incuded WT controls, *Hexb*^-/-^ controls, *Hexb*^-/-^ BMT, and *Hexb*^-/-^ BMT + CSF1Ri. Clusters Clusters were annotated with a combination of automated and manual approaches: 1) label annotations from the Allen Brain Atlas single-cell RNA-seq reference dataset (for cortex and hippocampus) were projected onto our spatial transcriptomics dataset^34^, and 2) cluster identities were further refined via manual annotation based on gene expression of known marker genes and location in XY space. Cell proportion plots were generated by first plotting the number of cells in each broad cell type, then scaling to 1. normalized percentages for each group, calculated by dividing the number of cells in a given cell type-group pair by the total number of cells in that group, and 2. dividing by the sum of the proportions across the cell type to account for differences in sample sizes. Differential gene expression analysis per cell type between groups was performed on scaled expression data using MAST to calculate the average difference^173^, defined as the difference in log-scaled average expression between the two groups for each broad cell type. DEG scores were calculated between group pairs for each subcluster by summing the absolute log2 fold change values of all genes with statistically significant gene (i.e., padj < 0.05) differential expression patterns between two groups. Data visualizations were generated using ggplot2 3.4.4^174^.

### Biochemical analysis of salt-soluble and detergent soluble protein fractions

*Fractionation:* Protein fractions were obtained from fresh frozen hemispheres which were first pulverized using a Bessman Tissue Pulverizer. Samples were then homogenized in 10μL high salt reassembly buffer per 1 mg of sample (RAB buffer; C752K77; Thermo Fisher; 100 mM MES, 1 mM EGTA, 0.5 mM MgSO4, 750 mM NaCl, 20 mM NaF, 1 mM Na3VO4, pH = 7.0).

Homogenates were centrifuged at 44,000g for 20 minutes and the supernatant was collected as the salt-soluble fraction. The pellet was then resuspended in 10uL of detergent-containing Tissue Protein Extraction Reagent (T-PER 25 mM bicine and 150 mM sodium chloride (pH 7.6); Life Technologies, Grand Island, NY) per 1 mg of original sample to gently extract total protein, then centrifuged at 44,000g for 1 hour. Supernatant was then collected as the detergent-soluble fraction. Fractions were analyzed using the Hexβ activity assay as described below and values were normalized to protein concentration for each sample.

*Protein quantification:* Total protein in salt-soluble and detergent-soluble fractions was quantified using the Pierce™ 660 nm Protein Assay Kit (#22662, Thermo Fisher, Waltham, MA). BSA standards were created for RAB and T-PER-extracted samples by diluting kit-supplied protein assay standards in extraction media 1:1. Fractions were removed from -80°C and thawed on ice. A 1:5 dilution of each sample was created by dilution in extraction media. Diluted BSA standards and 10 μL of diluted samples were loaded onto 96-well plates in triplicate, then 150 μL of Pierce Reagent was added to each well with a multichannel pipette using reverse pipetting. Plates were agitated on a plate shaker to 1 minute, then incubated at room temperature for 5 minutes before absorbances were read on a 96-well colorimetric and fluorescent microplate reader. Average absorbances were calculated for each sample, and protein concentration was determined using the standard curve of each plate.

### Primary glial cultures from neonatal mice

Primary mixed microglia-astrocyte cultures were generated as previously described^175^. Whole brains were extracted from neonatal 3- to 5-day-old mice and cortical tissue was cut into small pieces before digestion with trypsin. Trypsin was quenched using glia media (DMEM supplemented with 10% performance plus heat inactivated serum [#10082147; Thermo Fisher, Waltham, MA] and 1% penicillin/streptomycin [P4333-100ML; Sigma Aldrich, Saint Louis, MO]) and tissue was dissociated by pipetting up and down 20 times with a 1000-μL tip a total of 6 times. Following dissociation, the tissue was centrifuged at 150g for 7 minutes with slow start-stop at room temperature. Cells were resuspended in fresh glia media and filtered using 100-μm cell strainers (#352360; Falcon, Abilene, TX), followed by 40-μm strainers (#352340; Falcon). Finally, the cells were reconstituted with 10 mL of glia media and placed in T-75 cm2 flasks. Flasks were precoated with 0.002% poly-lysine (P4707-50ML; Sigma-Aldrich) for at least 30 minutes at room temperature. After 24 hours, debris was removed by gentle tapping of flasks and removal of media, and 20 mL of fresh media was added to the cell cultures. After 14 days in vitro, mixed microglia-astrocyte cultures were used for experiments.

### Primary microglia monoculture

Primary microglia were removed from mixed microglia-astrocyte culture by gentle shaking as described previously^175^. Microglia were seeded at 20,000/150μl on pre-coated 0.002% poly-lysine 96 well plates in 1:2 (conditioned: fresh) media and left to adhere for 48 hours.

### Cell treatments and β-hexosaminidase activity assay

The Hexβ acitivity assay was run as previously described^112^. Cells were washed with 1x PBS followed by 150μl of fresh media and treated with DMSO (vehicle), 400nM vacuolin-1 (673000, Millipore Sigma), 10μM BAPTA-AM (A1076, Millipore Sigma) and 20μM GW4869 (D1692, Millipore Sigma) for 6h. For P2X7 inhibition experiments, cells were treated with 1μg/ml LPS (L4130, Millipore Sigma) for 3h, followed by 10μM of P2X7 inhibitor, A-804598 (A16066, Tocris) for 10 min and 1mM ATP (A0157, TCI) for 20 min. The media (supernatant) was collected, and cells were lysed in 150μl of 1x M-PER supplemented with protease and phosphatase inhibitors for 20 min, on ice. Following lysis, samples were centrifuged at 15,000g for 10 min. The β- hexosaminidase assay was performed in a 96-well plate by mixing 50 μL of 2 mg/mL 4-nitrophenyl N-acetyl-β-D-galactosaminide (N9003, Millipore Sigma) in 0.1 M citrate buffer (pH 4.5) with 75 μL of supernatant or cell lysate and incubating for 1 hour at 37° C. Following 1h incubation, 100μl of 0.2M borate buffer (pH 9.8) was added to stop the reaction. The plate was read at 405nm using an absorbance plate reader. Percentage values were obtained by dividing the reading from the supernatant with that of the cell lysate.

### E18 hippocampal neuron cultures and imaging

Dissociated E18 hippocampal neurons were purchased from Brain Bits by Transnetyx (C57EDHP). 50,000 cells were plated on (50μg/ml) poly-D-lysine (A3890401, Thermofisher) pre-coated glass bottom plates (P35G-1.5-14-C, Mattek) in NbActiv1 media (Neurobasal/B27/Glutamax). Half media swaps were completed every 3-4 days with NbActiv1 media without Glutamax supplemented. Neurons were left for a minimum of 7 days before experiments were conducted. 10μg of recombinant C terminal 6x his-tagged mouse Hexβ was added to neurons for 24h. Following incubation, cells were washed 3 x 5 min each with 1X PBS. Neurons were fixed with 4% PFA for 15 min at room temperature. Following fixation, cells were washed a further 3 x 5 min each before adding blocking buffer (5% normal goat serum with 0.02% triton-x 100 in 1X PBS) for 1 hour at room temperature with gentle shaking. Cells were then incubated with primary antibodies overnight at 4°C. Primary antibodies used were rat anti-mouse LAMP1 (Ab25245, ABCAM), rabbit anti-6x his-tag (MA5-33032, Invitrogen), and mouse anti-mouse NeuN (MA5, 33103, Invitrogen). Following incubation, cells were washed 3 times with 1X PBS for 5 minutes each and incubated with secondary antibodies Alexa Fluor 633 (A21094, Thermofisher), Alexa Fluor 555 (A21422, Thermofisher) and Alexa Fluor 488 (A11034, Thermofisher) for 1 hour at room temperature with gentle shaking. Finally, cells were washed 3 times with 1X PBS for 5 minutes and imaged using LSM 900 (Carl Zeiss) 10 × 0.45 NA air objective and 4× digital zoom. 12-13 neurons were imaged per treatment condition.

---

Extended Data Fig. 1: Spatial transcriptomic cell segmentation and expanded data visualization, *Hexb^-/-^* vs. WT.

(a) Representative images demonstrating cell segmentation in cortex, choroid plexus/ventricle, corpus callosum, and dentate gyrus. Cells were imaged with rRNA (not shown), Histone, DAPI, and GFAP markers and segmented automatically. (b) Uniform Manifold Approximation and Projection (UMAP) of 39 clusters split by group. (c) 39 clusters plotted in XY space in all 6 brains from WT control and *Hexb*^-/-^ control brains (n=3/group). (d) Bar graph of proportions of cell counts by broad cell type per group.

Extended Data Fig. 2: Heatmap of top 5 marker genes for all spatial transcriptomics subclusters, *Hexb^-/-^* vs. WT.

(a) Heatmap of top 5 marker genes for each subcluster. DGE analysis was performed between each subcluster compared to all other subclusters to identify top 5 genes enriched in each subcluster.

Extended Data Fig. 3: Subcluster annotation information and DEG scores for all clusters identified by spatial transcriptomics, *Hexb^-/-^*vs. WT.

(a) Uniform Manifold Approximation and Projection (UMAP) of 39 clusters showing transcript expression of canonical marker genes for different broad cell types (purple = low, yellow = high).

(b) Bar graph of DEG scores in all subclusters.

Extended Data Fig. 4: Volcano plots for all spatial transcriptomics subclusters, *Hexb^-/-^*vs. WT.

(a) Volcano plots of DEGs between *Hexb^-/-^* and WT control for all cellular subclusters.

Extended Data Fig. 5: Confirmation of successful BMT by flow cytometry and correlation analysis of regional peripheral cell infiltration versus final week Rotarod score.

(a) Schematic of sample collection for assessment of donor chimerism. Whole bone marrow and whole blood were collected from chimeric mice at point of sacrifice. Red blood cells (RBCs) were lysed to enrich for hematopoietic stem cells (HSC; Ter119^−^CD27^+^ckit^+^Sca^+^CD150^+^CD34^−^ cells) In bone marrow and granulocytes (CD45^+^NK1.1^−^CD11b^+^GR1/Ly6G^+^ cells) in blood. (b) Representative flow cytometry plot showing gating strategy for granulocytes. (c) Bar graph of percent donor chimerism of all BMT (BMT-treated WT, BMT + CSF1Ri-treated WT, *Hexb*^-/-^, BMT- treated *Hexb*^-/-^, and BMT + CSF1Ri-treated *Hexb*^-/-^) mice. Donor chimerism was assessed by % green fluorescent protein (GFP)^+^ cells (GFP^+^ cells/total cells). (d-e) Scatterplots with line of best fit of final week (week 16) Rotarod latency-to-fall time (x axis) versus total GFP^+^ staining volume (y axis) in 20x confocal FOVs from (d) somatosensory cortex and (e) cerebellum in BMT + CSF1Ri-treated *Hexb*^-/-^ mice. (f) Scatterplot with line of best fit of final week (week 16) Rotarod latency-to-fall time (x axis) versus number of green fluorescent (GFP)^+^ spots/cells (y axis) in forebrain portion of whole brain sagittal scans of BMT + CSF1Ri-treated *Hexb*^-/-^ mice.

Extended Data Fig. 6: Expanded spatial transcriptomic data visualization, WT and all *Hexb^- /-^* groups. (a) Uniform Manifold Approximation and Projection (UMAP) of 38 clusters split by genotype and treatment condition group. (b) 38 clusters plotted in XY space in all 12 brains from WT control, *Hexb*^-/-^ control, BMT-treated *Hexb*^-/-^, and BMT + CSF1Ri-treated *Hexb*^-/-^ brains (n=3/group). (d) Bar graph of proportions of cell counts by broad cell type per group.

Extended Data Fig. 7: Heatmap of top 5 marker genes for all spatial transcriptomics subclusters, WT and all *Hexb^-/-^* groups. (a) Heatmap of top 5 marker genes for each subcluster. DGE analysis was performed between each subcluster compared to all other subclusters to identify top 5 genes enriched in each subcluster.

Extended Data Fig. 8: Spatial transcriptomics broad cell type differentially expressed genes, treated *Hexb^-/-^* versus *Hexb^-/-^*control. Volcano plots of DEGs between (a) BMT + CSF1Ri-treated *Hexb^-/-^* and *Hexb^-/-^* control and (b) BMT-treated *Hexb^-/-^* and *Hexb^-/-^* control for each broad cell type.

Extended Data Fig. 9: Spatial transcriptomics broad cell type differentially expressed genes, treated *Hexb^-/-^* versus WT*^-^* control. Volcano plots of DEGs between (a) BMT + CSF1Ri- treated *Hexb^-/-^* and WT*^-^* control and (b) BMT-treated *Hexb^-/-^* and WT control for each broad cell type.

Extended Data Fig. 10: Spatial transcriptomics broad cell type differentially expressed genes, BMT-treated *Hexb^-/-^* versus BMT-treated *Hexb^-/-^*, and broad cell type pseudobulk analysis, WT and all *Hexb^-/-^* groups. (a) Volcano plots of DEGs between BMT-treated *Hexb^-/-^* and BMT + CSF1Ri-treated *Hexb^-/-^* for each broad cell type. (b-d) Dot plots representing pseudo- bulked expression values across the four animal groups (WT control, *Hexb*^-/-^ control, *Hexb*^-/-^ BMT-treated, and *Hexb*^-/-^ BMT + CSF1Ri-treated). Pseudo-bulk analysis was performed to identify the top DEGs in the (b) astrocyte, (c) endothelial cell, and (d) excitatory neuron broad cell types and plotted across the four animal groups.

Extended Data Fig. 11: Spatial transcriptomics broad cell type pseudobulk analysis, WT and all *Hexb^-/-^* groups. Dot plots representing pseudo-bulked expression values across the four animal groups (WT control, *Hexb*^-/-^ control, BMT-treated *Hexb*^-/-^, and BMT + CSF1Ri-treated *Hexb*^-/-^). Pseudo-bulk analysis was performed to identify the top DEGs in the (a) inhibitory neuron, (b) myeloid, (c) oligodendrocyte, (d) oligodendrocyte precursor cell, and (e) vascular broad cell types and plotted across the four animal groups.

Extended Data Fig. 12: Expanded spatial proteomics data visualization. (a) Projection of all brains (WT control, *Hexb*^-/-^ control, BMT-treated *Hexb*^-/-^, and BMT + CSF1Ri-treated *Hexb*^-/-^) in XY space. 4 brains per groups across 4 slides with 4 brains per slide. (b-f) Bubble plots of differentially expressed proteins (DEPs) of interest in broad cell types between pairs (b) *Hexb^-/-^* control vs. WT control, (c) BMT- treated *Hexb^-/-^* vs. *Hexb^-/-^* control, (d) BMT + CSF1Ri-treated *Hexb^-/-^* vs. *Hexb^-/-^* control, and (e) BMT + CSF1Ri-treated *Hexb^-/-^* vs. BMT-treated *Hexb^-/-^.* Dots are sized by p value (-log10p value) and colored by average difference (log2 fold change, red indicating increased expression, blue indicating decreased expression) of each DEP.

## REFERENCES

1. Platt, F. M., d’Azzo, A., Davidson, B. L., Neufeld, E. F. & Tifft, C. J. Lysosomal storage diseases. Nat. Rev. Dis. Primer 4, 1–25 (2018).

2. Fuller, M., Meikle, P. J. & Hopwood, J. J. Epidemiology of lysosomal storage diseases: an overview. in Fabry Disease: Perspectives from 5 Years of FOS (eds. Mehta, A., Beck, M. & Sunder-Plassmann, G.) (Oxford PharmaGenesis, Oxford, 2006).

3. Huang, J. Q. et al. Apoptotic cell death in mouse models of GM2 gangliosidosis and observations on human Tay-Sachs and Sandhoff diseases. Hum. Mol. Genet. 6, 1879–1885 (1997).

4. Conzelmann, E. & Sandhoff, K. Purification and characterization of an activator protein for the degradation of glycolipids GM2 and GA2 by hexosaminidase A. Hoppe. Seylers Z. Physiol. Chem. 360, 1837–1849 (1979).

5. Robinson, D. & Stirling, J. L. N-Acetyl-beta-glucosaminidases in human spleen. Biochem. J. 107, 321–327 (1968).

6. Srivastava, S. K. & Beutler, E. Hexosaminidase-A and hexosaminidase-B: studies in Tay-Sachs’ and Sandhoff’s disease. Nature 241, 463 (1973).

7. Sango, K. et al. Mouse models of Tay-Sachs and Sandhoff diseases differ in neurologic phenotype and ganglioside metabolism. Nat. Genet. 11, 170–176 (1995).

8. Itokazu, Y., Fuchigami, T. & Yu, R. K. Functional Impairment of the Nervous System with Glycolipid Deficiencies. Adv. Neurobiol. 29, 419–448 (2023).

9. Wang, A. et al. Innate immune sensing of lysosomal dysfunction drives multiple lysosomal storage disorders. Nat. Cell Biol. 26, 219–234 (2024).

10. Bley, A. E. et al. Natural history of infantile G(M2) gangliosidosis. Pediatrics 128, e1233–1241 (2011).

11. Wada, R., Tifft, C. J. & Proia, R. L. Microglial activation precedes acute neurodegeneration in Sandhoff disease and is suppressed by bone marrow transplantation. Proc. Natl. Acad. Sci. 97, 10954–10959 (2000).

12. Biffi, A. Hematopoietic Stem Cell Gene Therapy for Storage Disease: Current and New Indications. Mol. Ther. 25, 1155–1162 (2017).

13. Beck, M. Treatment strategies for lysosomal storage disorders. Dev. Med. Child Neurol. 60, 13–18 (2018).

14. Prasad, V. K. et al. Unrelated donor umbilical cord blood transplantation for inherited metabolic disorders in 159 pediatric patients from a single center: influence of cellular composition of the graft on transplantation outcomes. Blood 112, 2979–2989 (2008).

15. Jacobs, J. F. M., Willemsen, M. a. a. P., Groot-Loonen, J. J., Wevers, R. A. & Hoogerbrugge, P. M. Allogeneic BMT followed by substrate reduction therapy in a child with subacute Tay- Sachs disease. Bone Marrow Transplant. 36, 925–926 (2005).

16. Norflus, F., et al. Bone marrow transplantation prolongs life span and ameliorates neurologic manifestations in Sandhoff disease mice. J. Clin. Invest. 101, 1881–1888 (1998).

17. Beattie, L. et al. Bone marrow-derived and resident liver macrophages display unique transcriptomic signatures but similar biological functions. J. Hepatol. 65, 758–768 (2016).

18. Kennedy, D. W. & Abkowitz, J. L. Kinetics of Central Nervous System Microglial and Macrophage Engraftment: Analysis Using a Transgenic Bone Marrow Transplantation Model. Blood 90, 986–993 (1997).

19. Hohsfield, L. A., et al. Effects of long-term and brain-wide colonization of peripheral bone marrow-derived myeloid cells in the CNS. J. Neuroinflammation 17, 279 (2020).

20. Loeb, A. M., Pattwell, S. S., Meshinchi, S., Bedalov, A. & Loeb, K. R. Donor bone marrow- derived macrophage engraftment into the central nervous system of patients following allogeneic transplantation. Blood Adv. 7, 5851–5859 (2023).

21. Du, J., Yang, B., He, Y. & Rao, Y. Evaluate the efficiency of Mr BMT in macrophage replacement of peripheral organs. 2024.06.22.600056 Preprint at 10.1101/2024.06.22.600056 (2024).

22. Jeyakumar, M. et al. Central nervous system inflammation is a hallmark of pathogenesis in mouse models of GM1 and GM2 gangliosidosis. Brain 126, 974–987 (2003).

23. Kuil, L. E. et al. Hexb enzyme deficiency leads to lysosomal abnormalities in radial glia and microglia in zebrafish brain development. Glia 67, 1705–1718 (2019).

24. Allende, M. L. et al. Genetic defects in the sphingolipid degradation pathway and their effects on microglia in neurodegenerative disease. Cell. Signal. 78, 109879 (2021).

25. Masuda, T. et al. Novel Hexb-based tools for studying microglia in the CNS. Nat. Immunol. 21, 802–815 (2020).

26. Shah, S. et al. Microglia-Specific Promoter Activities of HEXB Gene. Front. Cell. Neurosci. 16, (2022).

27. Elmore, M. R. P. et al. Colony-Stimulating Factor 1 Receptor Signaling Is Necessary for Microglia Viability, Unmasking a Microglia Progenitor Cell in the Adult Brain. Neuron 82, 380– 397 (2014).

28. Hohsfield, L. A., et al. MAC2 is a long-lasting marker of peripheral cell infiltrates into the mouse CNS after bone marrow transplantation and coronavirus infection. Glia 70, 875–891 (2022).

29. Xu, Z., Zhou, X., Peng, B. & Rao, Y. Microglia replacement by bone marrow transplantation (Mr BMT) in the central nervous system of adult mice. STAR Protoc. 2, 100666 (2021).

30. Colella, P. et al. CNS-wide repopulation by hematopoietic-derived microglia-like cells corrects progranulin deficiency in mice. Nat. Commun. 15, 5654 (2024).

31. Mader, M. M.-D. et al. Myeloid cell replacement is neuroprotective in chronic experimental autoimmune encephalomyelitis. Nat. Neurosci. 27, 901–912 (2024).

32. Shibuya, Y. et al. Treatment of a genetic brain disease by CNS-wide microglia replacement. Sci. Transl. Med. 14, eabl9945 (2022).

33. He, S. et al. High-plex imaging of RNA and proteins at subcellular resolution in fixed tissue by spatial molecular imaging. Nat. Biotechnol. 40, 1794–1806 (2022).

34. Yao, Z. et al. A taxonomy of transcriptomic cell types across the isocortex and hippocampal formation. Cell 184, 3222–3241.e26 (2021).

35. Dang, D. et al. Computational Approach to Identifying Universal Macrophage Biomarkers. Front. Physiol. 11, 275 (2020).

36. Stables, M. J. et al. Transcriptomic analyses of murine resolution-phase macrophages. Blood 118, e192–e208 (2011).

37. Cross, M. & Renkawitz, R. Repetitive sequence involvement in the duplication and divergence of mouse lysozyme genes. EMBO J. 9, 1283–1288 (1990).

38. Monaghan, K. L., Zheng, W., Hu, G. & Wan, E. C. K. Monocytes and Monocyte-Derived Antigen-Presenting Cells Have Distinct Gene Signatures in Experimental Model of Multiple Sclerosis. Front. Immunol. 10, 2779 (2019).

39. Wu, Y.-P. & Proia, R. L. Deletion of macrophage-inflammatory protein 1 alpha retards neurodegeneration in Sandhoff disease mice. Proc. Natl. Acad. Sci. U. S. A. 101, 8425–8430 (2004).

40. Pekny, M., Wilhelmsson, U. & Pekna, M. The dual role of astrocyte activation and reactive gliosis. Neurosci. Lett. 565, 30–38 (2014).

41. Lan, Y.-L., Fang, D.-Y., Zhao, J., Ma, T.-H. & Li, S. A research update on the potential roles of aquaporin 4 in neuroinflammation. Acta Neurol. Belg. 116, 127–134 (2016).

42. Preman, P., Alfonso-Triguero, M., Alberdi, E., Verkhratsky, A. & Arranz, A. M. Astrocytes in Alzheimer’s Disease: Pathological Significance and Molecular Pathways. Cells 10, 540 (2021).

43. Habib, N. et al. Disease-associated astrocytes in Alzheimer’s disease and aging. Nat. Neurosci. 23, 701–706 (2020).

44. Kajiwara, Y. et al. GJA1 (connexin43) is a key regulator of Alzheimer’s disease pathogenesis. Acta Neuropathol. Commun. 6, 144 (2018).

45. Castranio, E. L. et al. Gene co-expression networks identify Trem2 and Tyrobp as major hubs in human APOE expressing mice following traumatic brain injury. Neurobiol. Dis. 105, 1– 14 (2017).

46. York, I. A. & Rock, K. L. Antigen processing and presentation by the class I major histocompatibility complex. Annu. Rev. Immunol. 14, 369–396 (1996).

47. Glynn, M. W. et al. MHCI negatively regulates synapse density during the establishment of cortical connections. Nat. Neurosci. 14, 442–451 (2011).

48. Nakanishi, H. Microglial cathepsin B as a key driver of inflammatory brain diseases and brain aging. Neural Regen. Res. 15, 25–29 (2020).

49. Nakanishi, H. Neuronal and microglial cathepsins in aging and age-related diseases. Ageing Res. Rev. 2, 367–381 (2003).

50. Ishizaka, M., Ohe, Y., Senbongi, T., Wakabayashi, K. & Ishikawa, K. Inflammatory stimuli increase prostaglandin D synthase levels in cerebrospinal fluid of rats. Neuroreport 12, 1161– 1165 (2001).

51. Schoenebeck, B. et al. Sgk1, a cell survival response in neurodegenerative diseases. Mol. Cell. Neurosci. 30, 249–264 (2005).

52. Zhang, J. et al. Progression of the role of CRYAB in signaling pathways and cancers. OncoTargets Ther. 12, 4129–4139 (2019).

53. Kuhn, S., Gritti, L., Crooks, D. & Dombrowski, Y. Oligodendrocytes in Development, Myelin Generation and Beyond. Cells 8, 1424 (2019).

54. Ragolia, L., Palaia, T., Frese, L., Fishbane, S. & Maesaka, J. K. Prostaglandin D2 synthase induces apoptosis in PC12 neuronal cells. Neuroreport 12, 2623–2628 (2001).

55. Flügge, G., Araya-Callis, C., Garea-Rodriguez, E., Stadelmann-Nessler, C. & Fuchs, E. NDRG2 as a marker protein for brain astrocytes. Cell Tissue Res. 357, 31–41 (2014).

56. Svirsky, S. et al. Neurogranin Protein Expression Is Reduced after Controlled Cortical Impact in Rats. J. Neurotrauma 37, 939–949 (2020).

57. Killoy, K. M., Harlan, B. A., Pehar, M. & Vargas, M. R. FABP7 up-regulation induces a neurotoxic phenotype in astrocytes. Glia 68, 2693–2704 (2020).

58. Hoyaux, D. et al. S100A6 overexpression within astrocytes associated with impaired axons from both ALS mouse model and human patients. J. Neuropathol. Exp. Neurol. 61, 736–744 (2002).

59. Fricker, M. et al. MFG-E8 Mediates Primary Phagocytosis of Viable Neurons during Neuroinflammation. J. Neurosci. 32, 2657–2666 (2012).

60. Arnaud, L. et al. APOE4 drives inflammation in human astrocytes via TAGLN3 repression and NF-κB activation. Cell Rep. 40, 111200 (2022).

61. Antonucci, F. et al. SNAP-25, a Known Presynaptic Protein with Emerging Postsynaptic Functions. Front. Synaptic Neurosci. 8, (2016).

62. Ichise, E. et al. Impaired neuronal activity and differential gene expression in STXBP1 encephalopathy patient iPSC-derived GABAergic neurons. Hum. Mol. Genet. 30, 1337–1348 (2021).

63. Itoh, M. et al. Lysosomal cholesterol overload in macrophages promotes liver fibrosis in a mouse model of NASH. J. Exp. Med. 220, e20220681 (2023).

64. Cesana, M. et al. EGR1 drives cell proliferation by directly stimulating TFEB transcription in response to starvation. PLOS Biol. 21, e3002034 (2023).

65. Xie, B. et al. Egr-1 transactivates Bim gene expression to promote neuronal apoptosis. J. Neurosci. Off. J. Soc. Neurosci. 31, 5032–5044 (2011).

66. Harashima, S., Wang, Y., Horiuchi, T., Seino, Y. & Inagaki, N. Purkinje cell protein 4 positively regulates neurite outgrowth and neurotransmitter release. J. Neurosci. Res. 89, 1519–1530 (2011).

67. Yu, H., Han, Y., Cui, C., Li, G. & Zhang, B. Loss of SV2A promotes human neural stem cell apoptosis via p53 signaling. Neurosci. Lett. 800, 137125 (2023).

68. Platzer, K. et al. De Novo Missense Variants in SLC32A1 Cause a Developmental and Epileptic Encephalopathy Due to Impaired GABAergic Neurotransmission. Ann. Neurol. 92, 958–973 (2022).

69. Foltz, G. et al. Genome-wide analysis of epigenetic silencing identifies BEX1 and BEX2 as candidate tumor suppressor genes in malignant glioma. Cancer Res. 66, 6665–6674 (2006).

70. Yang, L. et al. ZWINT: A potential therapeutic biomarker in patients with glioblastoma correlates with cell proliferation and invasion. Oncol. Rep. 43, 1831–1844 (2020).

71. Hallmann, K. et al. Loss of the smallest subunit of cytochrome c oxidase, COX8A, causes Leigh-like syndrome and epilepsy. Brain J. Neurol. 139, 338–345 (2016).

72. Arora, M., Kumari, S., Singh, J., Chopra, A. & Chauhan, S. S. Downregulation of Brain Enriched Type 2 MAGEs Is Associated With Immune Infiltration and Poor Prognosis in Glioma. Front. Oncol. 10, 573378 (2020).

73. Dukay, B., Csoboz, B. & Tóth, M. E. Heat-Shock Proteins in Neuroinflammation. Front. Pharmacol. 10, (2019).

74. Bonam, S. R., Ruff, M. & Muller, S. HSPA8/HSC70 in Immune Disorders: A Molecular Rheostat that Adjusts Chaperone-Mediated Autophagy Substrates. Cells 8, 849 (2019).

75. Saouab, R. et al. A case report of Sandhoff disease. Clin. Neuroradiol. 21, 83–85 (2011).

76. Beker-Acay, M., Elmas, M., Koken, R., Unlu, E. & Bukulmez, A. Infantile Type Sandhoff Disease with Striking Brain MRI Findings and a Novel Mutation. Pol. J. Radiol. 81, 86–89 (2016).

77. Calişkan, M., Ozmen, M., Beck, M. & Apak, S. Thalamic hyperdensity--is it a diagnostic marker for Sandhoff disease? Brain Dev. 15, 387–388 (1993).

78. Myerowitz, R. et al. Molecular pathophysiology in Tay-Sachs and Sandhoff diseases as revealed by gene expression profiling. Hum. Mol. Genet. 11, 1343–1350 (2002).

79. Cugurra, A. et al. Skull and vertebral bone marrow are myeloid cell reservoirs for the meninges and CNS parenchyma. Science 373, eabf7844 (2021).

80. Yang, Y. et al. Perivascular, but not parenchymal, cerebral engraftment of donor cells after non-myeloablative bone marrow transplantation. Exp. Mol. Pathol. 95, 7–17 (2013).

81. Jain, A., Kohli, A. & Sachan, D. Infantile Sandhoff’s disease with peripheral neuropathy. Pediatr. Neurol. 42, 459–461 (2010).

82. Bennett, M. L. et al. New tools for studying microglia in the mouse and human CNS. Proc. Natl. Acad. Sci. U. S. A. 113, E1738–1746 (2016).

83. Santos-Carvalho, A., Elvas, F., Alvaro, A. R., Ambrósio, A. F. & Cavadas, C. Neuropeptide Y receptors activation protects rat retinal neural cells against necrotic and apoptotic cell death induced by glutamate. Cell Death Dis. 4, e636 (2013).

84. Rehfeld, F. et al. The RNA-binding protein ARPP21 controls dendritic branching by functionally opposing the miRNA it hosts. Nat. Commun. 9, 1235 (2018).

85. Waschek, J. VIP and PACAP: neuropeptide modulators of CNS inflammation, injury, and repair. Br. J. Pharmacol. 169, 512–523 (2013).

86. Dicken, M. S., Hughes, A. R. & Hentges, S. T. Gad1 mRNA as a reliable indicator of altered GABA release from orexigenic neurons in the hypothalamus. Eur. J. Neurosci. 42, 2644–2653 (2015).

87. Keren-Shaul, H. et al. A Unique Microglia Type Associated with Restricting Development of Alzheimer’s Disease. Cell 169, 1276–1290.e17 (2017).

88. Nixon, R. A. Amyloid precursor protein and endosomal–lysosomal dysfunction in Alzheimer’s disease: inseparable partners in a multifactorial disease. FASEB J. 31, 2729–2743 (2017).

89. Nuriel, T. et al. The Endosomal–Lysosomal Pathway Is Dysregulated by APOE4 Expression in Vivo. Front. Neurosci. 11, 702 (2017).

90. Yadati, T., Houben, T., Bitorina, A. & Shiri-Sverdlov, R. The Ins and Outs of Cathepsins: Physiological Function and Role in Disease Management. Cells 9, 1679 (2020).

91. Ni, J., Lan, F., Xu, Y., Nakanishi, H. & Li, X. Extralysosomal cathepsin B in central nervous system: Mechanisms and therapeutic implications. Brain Pathol. 32, e13071 (2022).

92. Lee, J.-H. et al. Presenilin 1 maintains lysosomal Ca2+ homeostasis by regulating vATPase-mediated lysosome acidification. Cell Rep. 12, 1430–1444 (2015).

93. Zhou, Z. et al. The roles of amyloid precursor protein (APP) in neurogenesis, implications to pathogenesis and therapy of Alzheimer disease (AD). Cell Adhes. Migr. 5, 280–292 (2011).

94. Hanger, D. P., Anderton, B. H. & Noble, W. Tau phosphorylation: the therapeutic challenge for neurodegenerative disease. Trends Mol. Med. 15, 112–119 (2009).

95. Yan, K., Zhang, C., Kang, J., Montenegro, P. & Shen, J. Cortical neurodegeneration caused by Psen1 mutations is independent of Aβ. Proc. Natl. Acad. Sci. U. S. A. 121, e2409343121 (2024).

96. Schmidt, M. F., Gan, Z. Y., Komander, D. & Dewson, G. Ubiquitin signalling in neurodegeneration: mechanisms and therapeutic opportunities. Cell Death Differ. 28, 570– 590 (2021).

97. Ham, J., Eilers, A., Whitfield, J., Neame, S. J. & Shah, B. c-Jun and the transcriptional control of neuronal apoptosis. Biochem. Pharmacol. 60, 1015–1021 (2000).

98. Ayanlaja, A. A. et al. Distinct Features of Doublecortin as a Marker of Neuronal Migration and Its Implications in Cancer Cell Mobility. Front. Mol. Neurosci. 10, 199 (2017).

99. Mollinedo, F. & Gajate, C. Microtubules, microtubule-interfering agents and apoptosis. Apoptosis Int. J. Program. Cell Death 8, 413–450 (2003).

100. Lier, J., Streit, W. J. & Bechmann, I. Beyond Activation: Characterizing Microglial Functional Phenotypes. Cells 10, 2236 (2021).

101. Gómez Morillas, A., Besson, V. C. & Lerouet, D. Microglia and Neuroinflammation: What Place for P2RY12? Int. J. Mol. Sci. 22, 1636 (2021).

102. Jeyakumar, M. et al. Enhanced survival in Sandhoff disease mice receiving a combination of substrate deprivation therapy and bone marrow transplantation. Blood 97, 327–329 (2001).

103. Jeyakumar, M. et al. Delayed symptom onset and increased life expectancy in Sandhoff disease mice treated with N-butyldeoxynojirimycin. Proc. Natl. Acad. Sci. U. S. A. 96, 6388– 6393 (1999).

104. Ou, L. et al. A novel gene editing system to treat both Tay-Sachs and Sandhoff diseases. Gene Ther. 27, 226–236 (2020).

105. Walia, J. S. et al. Long-term correction of Sandhoff disease following intravenous delivery of rAAV9 to mouse neonates. Mol. Ther. J. Am. Soc. Gene Ther. 23, 414–422 (2015).

106. Bacioglu, M. et al. Neurofilament Light Chain in Blood and CSF as Marker of Disease Progression in Mouse Models and in Neurodegenerative Diseases. Neuron 91, 56–66 (2016).

107. Welford, R. W. D. et al. Plasma neurofilament light, glial fibrillary acidic protein and lysosphingolipid biomarkers for pharmacodynamics and disease monitoring of GM2 and GM1 gangliosidoses patients. Mol. Genet. Metab. Rep. 30, 100843 (2022).

108. Senior, J. R. Alanine aminotransferase: a clinical and regulatory tool for detecting liver injury-past, present, and future. Clin. Pharmacol. Ther. 92, 332–339 (2012).

109. Andersson, S. V., Sjögren, E. C., Magnusson, C. & Gierow, J. P. Sequencing, expression, and enzymatic characterization of beta-hexosaminidase in rabbit lacrimal gland and primary cultured acinar cells. Glycobiology 15, 211–220 (2005).

110. Dersh, D., Iwamoto, Y. & Argon, Y. Tay–Sachs disease mutations in HEXA target the α chain of hexosaminidase A to endoplasmic reticulum–associated degradation. Mol. Biol. Cell 27, 3813–3827 (2016).

111. Toyomitsu, E. et al. CCL2 promotes P2X4 receptor trafficking to the cell surface of microglia. Purinergic Signal. 8, 301–310 (2012).

112. Yanes, R. E. et al. Involvement of lysosomal exocytosis in the excretion of mesoporous silica nanoparticles and enhancement of the drug delivery effect by exocytosis inhibition. Small Weinh. Bergstr. Ger. 9, 697–704 (2013).

113. Shehadul Islam, M., Aryasomayajula, A. & Selvaganapathy, P. R. A Review on Macroscale and Microscale Cell Lysis Methods. Micromachines 8, 83 (2017).

114. Leoni, P. & Dean, R. T. Mechanisms of lysosomal enzyme secretion by human monocytes. Biochim. Biophys. Acta BBA - Mol. Cell Res. 762, 378–389 (1983).

115. Buratta, S. et al. Lysosomal Exocytosis, Exosome Release and Secretory Autophagy: The Autophagic- and Endo-Lysosomal Systems Go Extracellular. Int. J. Mol. Sci. 21, 2576 (2020).

116. Cerny, J. et al. The small chemical vacuolin-1 inhibits Ca(2+)-dependent lysosomal exocytosis but not cell resealing. EMBO Rep. 5, 883–888 (2004).

117. Tsien, R. Y. A non-disruptive technique for loading calcium buffers and indicators into cells. Nature 290, 527–528 (1981).

118. Essandoh, K. et al. Blockade of exosome generation with GW4869 dampens the sepsis- induced inflammation and cardiac dysfunction. Biochim. Biophys. Acta 1852, 2362–2371 (2015).

119. Zamyatnin, A. A., Gregory, L. C., Townsend, P. A. & Soond, S. M. Beyond basic research: the contribution of cathepsin B to cancer development, diagnosis and therapy. Expert Opin. Ther. Targets 26, 963–977 (2022).

120. Lien, E. et al. Toll-like receptor 4 imparts ligand-specific recognition of bacterial lipopolysaccharide. J. Clin. Invest. 105, 497–504 (2000).

121. Fiebich, B. L., Akter, S. & Akundi, R. S. The two-hit hypothesis for neuroinflammation: role of exogenous ATP in modulating inflammation in the brain. Front. Cell. Neurosci. 8, 260 (2014).

122. Chevriaux, A. et al. Cathepsin B Is Required for NLRP3 Inflammasome Activation in Macrophages, Through NLRP3 Interaction. Front. Cell Dev. Biol. **8**, 167 (2020).

123. Takenouchi, T. et al. The activation of P2X7 receptor induces cathepsin D-dependent production of a 20-kDa form of IL-1β under acidic extracellular pH in LPS-primed microglial cells. J. Neurochem. 117, 712–723 (2011).

124. Gu, B. J. & Wiley, J. S. P2X7 as a scavenger receptor for innate phagocytosis in the brain. Br. J. Pharmacol. 175, 4195–4208 (2018).

125. Campagno, K. E. & Mitchell, C. H. The P2X7 Receptor in Microglial Cells Modulates the Endolysosomal Axis, Autophagy, and Phagocytosis. Front. Cell. Neurosci. 15, (2021).

126. Egan, T. M. & Khakh, B. S. Contribution of calcium ions to P2X channel responses. J. Neurosci. Off. J. Soc. Neurosci. 24, 3413–3420 (2004).

127. Samways, D. S. K. et al. Quantifying Ca2+ Current and Permeability in ATP-gated P2X7 Receptors. J. Biol. Chem. 290, 7930–7942 (2015).

128. Donnelly-Roberts, D. L., Namovic, M. T., Han, P. & Jarvis, M. F. Mammalian P2X7 receptor pharmacology: comparison of recombinant mouse, rat and human P2X7 receptors. Br. J. Pharmacol. 157, 1203–1214 (2009).

129. Sekar, P., Huang, D.-Y., Hsieh, S.-L., Chang, S.-F. & Lin, W.-W. AMPK-dependent and independent actions of P2X7 in regulation of mitochondrial and lysosomal functions in microglia. Cell Commun. Signal. CCS 16, 83 (2018).

130. Adinolfi, E., et al. The P2X7 receptor: A main player in inflammation. Biochem. Pharmacol. 151, 234–244 (2018).

131. Abo-ouf, H. et al. Deletion of tumor necrosis factor-α ameliorates neurodegeneration in Sandhoff disease mice. Hum. Mol. Genet. 22, 3960–3975 (2013).

132. Kyrkanides, S. et al. Conditional expression of human β-hexosaminidase in the neurons of Sandhoff disease rescues mice from neurodegeneration but not neuroinflammation. J. Neuroinflammation 9, 186 (2012).

133. Cachón-González, M. B. et al. Effective gene therapy in an authentic model of Tay-Sachs- related diseases. Proc. Natl. Acad. Sci. U. S. A. 103, 10373–10378 (2006).

134. Cachón-González, M. B. et al. Gene transfer corrects acute GM2 gangliosidosis--potential therapeutic contribution of perivascular enzyme flow. Mol. Ther. J. Am. Soc. Gene Ther. 20, 1489–1500 (2012).

135. Beegle, J., Hendrix, K., Maciel, H., Nolta, J. A. & Anderson, J. S. Improvement of motor and behavioral activity in Sandhoff mice transplanted with human CD34+ cells transduced with a HexA/HexB expressing lentiviral vector. J. Gene Med. 22, e3205 (2020).

136. Niemir, N. et al. Intravenous administration of scAAV9-Hexb normalizes lifespan and prevents pathology in Sandhoff disease mice. Hum. Mol. Genet. 27, 954–968 (2018).

137. Sala, D. et al. Therapeutic advantages of combined gene/cell therapy strategies in a murine model of GM2 gangliosidosis. Mol. Ther. Methods Clin. Dev. 25, 170–189 (2022).

138. McCurdy, V. J. et al. Widespread correction of central nervous system disease after intracranial gene therapy in a feline model of Sandhoff disease. Gene Ther. 22, 181–189 (2015).

139. Kitchener, E. J. A., Dundee, J. M. & Brown, G. C. Activated microglia release β−galactosidase that promotes inflammatory neurodegeneration. Front. Aging Neurosci. 15, 1327756 (2024).

140. Allendorf, D. H. & Brown, G. C. Neu1 Is Released From Activated Microglia, Stimulating Microglial Phagocytosis and Sensitizing Neurons to Glutamate. Front. Cell. Neurosci. 16, 917884 (2022).

141. Čaval, T. et al. Targeted Analysis of Lysosomal Directed Proteins and Their Sites of Mannose-6-phosphate Modification. Mol. Cell. Proteomics MCP 18, 16–27 (2019).

142. Mahuran, D. J. Biochemical consequences of mutations causing the GM2 gangliosidoses. Biochim. Biophys. Acta BBA - Mol. Basis Dis. 1455, 105–138 (1999).

143. Hawkes, C. & Kar, S. The insulin-like growth factor-II/mannose-6-phosphate receptor: structure, distribution and function in the central nervous system. Brain Res. Rev. 44, 117– 140 (2004).

144. Gary-Bobo, M., Nirdé, P., Jeanjean, A., Morère, A. & Garcia, M. Mannose 6-phosphate receptor targeting and its applications in human diseases. Curr. Med. Chem. 14, 2945–2953 (2007).

145. Masciullo, M. et al. Substrate reduction therapy with miglustat in chronic GM2 gangliosidosis type Sandhoff: results of a 3-year follow-up. J. Inherit. Metab. Dis. 33 **Suppl 3**, S355–361 (2010).

146. Marshall, J. et al. Substrate Reduction Therapy for Sandhoff Disease through Inhibition of Glucosylceramide Synthase Activity. Mol. Ther. 27, 1495–1506 (2019).

147. Arthur, J. R., Lee, J. P., Snyder, E. Y. & Seyfried, T. N. Therapeutic effects of stem cells and substrate reduction in juvenile Sandhoff mice. Neurochem. Res. 37, 1335–1343 (2012).

148. Concolino, D., Deodato, F. & Parini, R. Enzyme replacement therapy: efficacy and limitations. Ital. J. Pediatr. 44, 120 (2018).

149. Tsuji, D. et al. Highly phosphomannosylated enzyme replacement therapy for GM2 gangliosidosis. Ann. Neurol. 69, 691–701 (2011).

150. Puhl, D. L., D’Amato, A. R. & Gilbert, R. J. Challenges of gene delivery to the central nervous system and the growing use of biomaterial vectors. Brain Res. Bull. 150, 216–230 (2019).

151. Mingozzi, F. & High, K. A. Immune responses to AAV in clinical trials. Curr. Gene Ther. 11, 321–330 (2011).

152. Golebiowski, D. et al. Direct Intracranial Injection of AAVrh8 Encoding Monkey β-N- Acetylhexosaminidase Causes Neurotoxicity in the Primate Brain. Hum. Gene Ther. 28, 510– 522 (2017).

153. Sharrack, B. et al. Autologous haematopoietic stem cell transplantation and other cellular therapy in multiple sclerosis and immune-mediated neurological diseases: updated guidelines and recommendations from the EBMT Autoimmune Diseases Working Party (ADWP) and the Joint Accreditation Committee of EBMT and ISCT (JACIE). Bone Marrow Transplant. 55, 283–306 (2020).

154. Platt, F. M. & Lachmann, R. H. Treating lysosomal storage disorders: Current practice and future prospects. Biochim. Biophys. Acta BBA - Mol. Cell Res. 1793, 737–745 (2009).

155. Tap, W. D. et al. Pexidartinib versus placebo for advanced tenosynovial giant cell tumour (ENLIVEN): a randomised phase 3 trial. Lancet Lond. Engl. 394, 478–487 (2019).

156. Bachiller, S. et al. Microglia in Neurological Diseases: A Road Map to Brain-Disease Dependent-Inflammatory Response. Front. Cell. Neurosci. 12, 488 (2018).

157. Colonna, M. & Butovsky, O. Microglia Function in the Central Nervous System During Health and Neurodegeneration. Annu. Rev. Immunol. 35, 441–468 (2017).

158. Hohsfield, L. A., et al. Subventricular zone/white matter microglia reconstitute the empty adult microglial niche in a dynamic wave. eLife 10, e66738 (2021).

159. Cronk, J. C. et al. Peripherally derived macrophages can engraft the brain independent of irradiation and maintain an identity distinct from microglia. J. Exp. Med. 215, 1627–1647 (2018).

160. Minogue, A. M. Role of infiltrating monocytes/macrophages in acute and chronic neuroinflammation: Effects on cognition, learning and affective behaviour. Prog. Neuropsychopharmacol. Biol. Psychiatry 79, 15–18 (2017).

161. Graves, M. C. et al. Inflammation in amyotrophic lateral sclerosis spinal cord and brain is mediated by activated macrophages, mast cells and T cells. Amyotroph. Lateral Scler. Mot. Neuron Disord. Off. Publ. World Fed. Neurol. Res. Group Mot. Neuron Dis. 5, 213–219 (2004).

162. Silvin, A., Qian, J. & Ginhoux, F. Brain macrophage development, diversity and dysregulation in health and disease. Cell. Mol. Immunol. 20, 1277–1289 (2023).

163. Douvaras, P. et al. Ready-to-use iPSC-derived microglia progenitors for the treatment of CNS disease in mouse models of neuropathic mucopolysaccharidoses. Nat. Commun. 15, 8132 (2024).

164. Chadarevian, J. P. et al. Engineering an inhibitor-resistant human CSF1R variant for microglia replacement. J. Exp. Med. 220, e20220857 (2023).

165. Hasselmann, J. & Blurton-Jones, M. Human iPSC-derived microglia: A growing toolset to study the brain’s innate immune cells. Glia 68, 721–739 (2020).

166. Abud, E. M. et al. iPSC-derived human microglia-like cells to study neurological diseases. Neuron 94, 278–293.e9 (2017).

167. Turnquist, C., Harris, B. T. & Harris, C. C. Radiation-induced brain injury: current concepts and therapeutic strategies targeting neuroinflammation. Neuro-Oncol. Adv. 2, vdaa057 (2020).

168. Magnuson, A., Mohile, S. & Janelsins, M. Cognition and Cognitive Impairment in Older Adults with Cancer. Curr. Geriatr. Rep. 5, 213–219 (2016).

169. Acharya, M. M. et al. Elimination of microglia improves cognitive function following cranial irradiation. Sci. Rep. 6, 31545 (2016).

170. Gabriel, M. et al. A Review of Acute and Long-Term Neurological Complications Following Haematopoietic Stem Cell Transplant for Paediatric Acute Lymphoblastic Leukaemia. Front. Pediatr. 9, 774853 (2021).

171. Spangenberg, E. et al. Sustained microglial depletion with CSF1R inhibitor impairs parenchymal plaque development in an Alzheimer’s disease model. Nat. Commun. 10, 3758 (2019).

172. Tran, K. M. et al. APOE Christchurch enhances a disease-associated microglial response to plaque but suppresses response to tau pathology. *bioRxiv* 2024.06.03.597211 (2024) doi:10.1101/2024.06.03.597211.

173. Finak, G. et al. MAST: a flexible statistical framework for assessing transcriptional changes and characterizing heterogeneity in single-cell RNA sequencing data. Genome Biol. 16, 278 (2015).

174. Wickham, H. Data Analysis. in *ggplot2: Elegant Graphics for Data Analysis* (ed. Wickham, H.) 189–201 (Springer International Publishing, Cham, 2016). doi:10.1007/978-3-319-24277-4_9.

175. Butler, C. A. et al. The Abca7V1613M variant reduces Aβ generation, plaque load, and neuronal damage. Alzheimers Dement. J. Alzheimers Assoc. 20, 4914–4934 (2024).

