## Extended Data Figures for "Myeloid-derived β-hexosaminidase is essential for neuronal health and lysosome function: implications for Sandhoff disease"

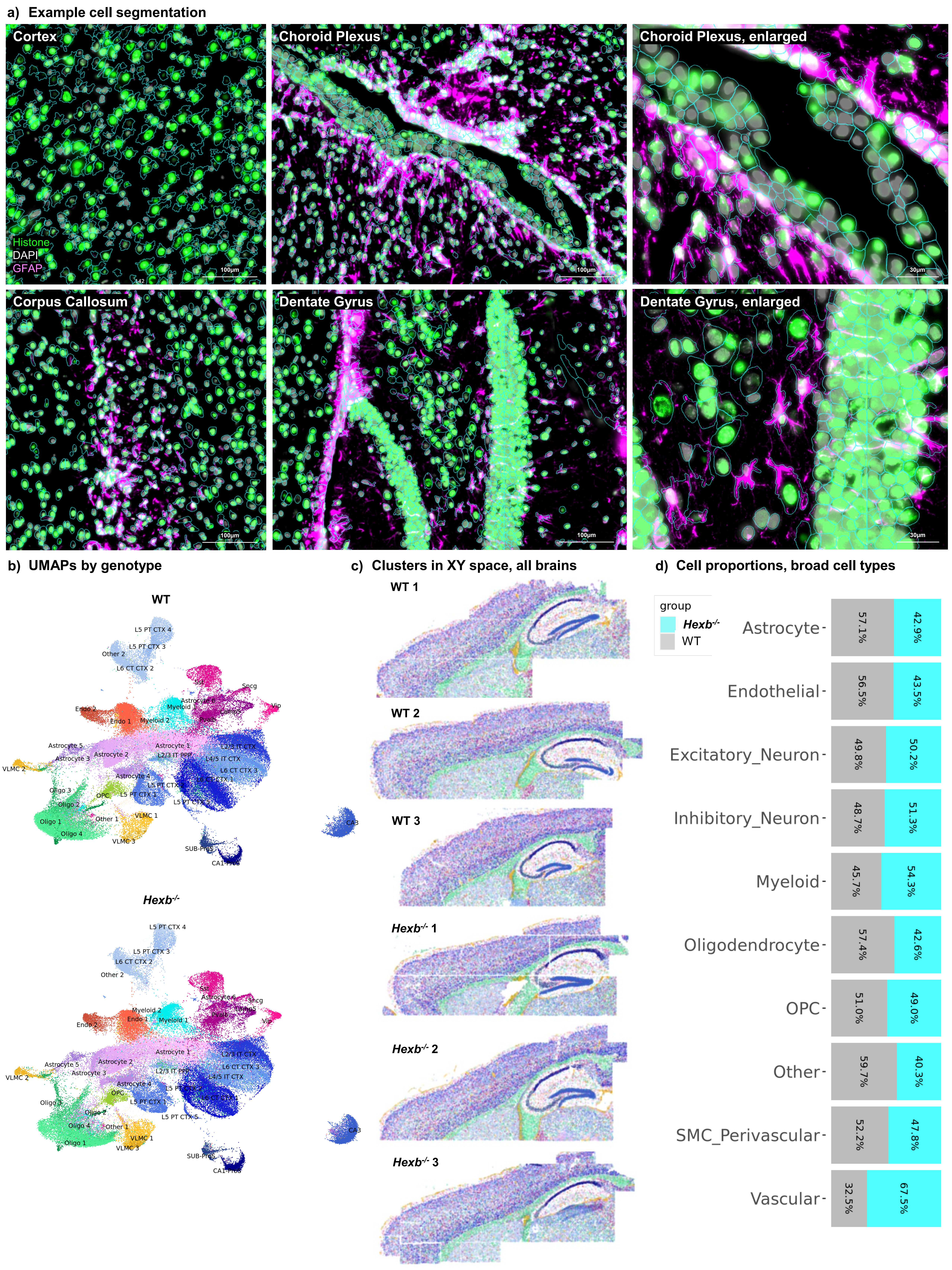
a) Example cell segmentation
Cortex
Choroid Plexus
Choroid Plexus, enlarged
Histone
DAPI
GFAP
100µm
30µm
Corpus Callosum
Dentate Gyrus
Dentate Gyrus, enlarged
b) UMAPs by genotype
WT
Hexb-/-
L5 PT CTX 4
L5 PT CTX 3
Other 2
L6 CT CTX 2
Sst
Sncg
Vip
Endo 2
Endo 1
Myeloid
Myeloid 2
Myeloid 1
Astrocyte 6
Lamp5
Pvalb
Astrocyte 5
Astrocyte 3
Astrocyte 2
Astrocyte 1
Astrocyte 4
L2/3 IT CTX
L2/3 IT PPP
L4/5 IT CTX
L6 CT CTX 3
L6 CT CTX 1
L5 PT CTX 2
L5 PT CTX 1
L5 PT CTX 5
VLMC 2
VLMC 1
VLMC 3
Oligo 1
Oligo 2
Oligo 3
Oligo 4
OPC
Other 1
SUB-Pros
CA1-Pros
CA3
c) Clusters in XY space, all brains
WT 1
WT 2
WT 3
Hexb-/- 1
Hexb-/- 2
Hexb-/- 3
d) Cell proportions, broad cell types
group
Hexb-/-
WT
Astrocyte 57.1% 42.9%
Endothelial 56.5% 43.5%
Excitatory_Neuron 49.8% 50.2%
Inhibitory_Neuron 48.7% 51.3%
Myeloid 45.7% 54.3%
Oligodendrocyte 57.4% 42.6%
OPC 51.0% 49.0%
Other 59.7% 40.3%
SMC_Perivascular 52.2% 47.8%
Vascular 32.5% 67.5%

**a) Top 5 marker genes per subcluster**

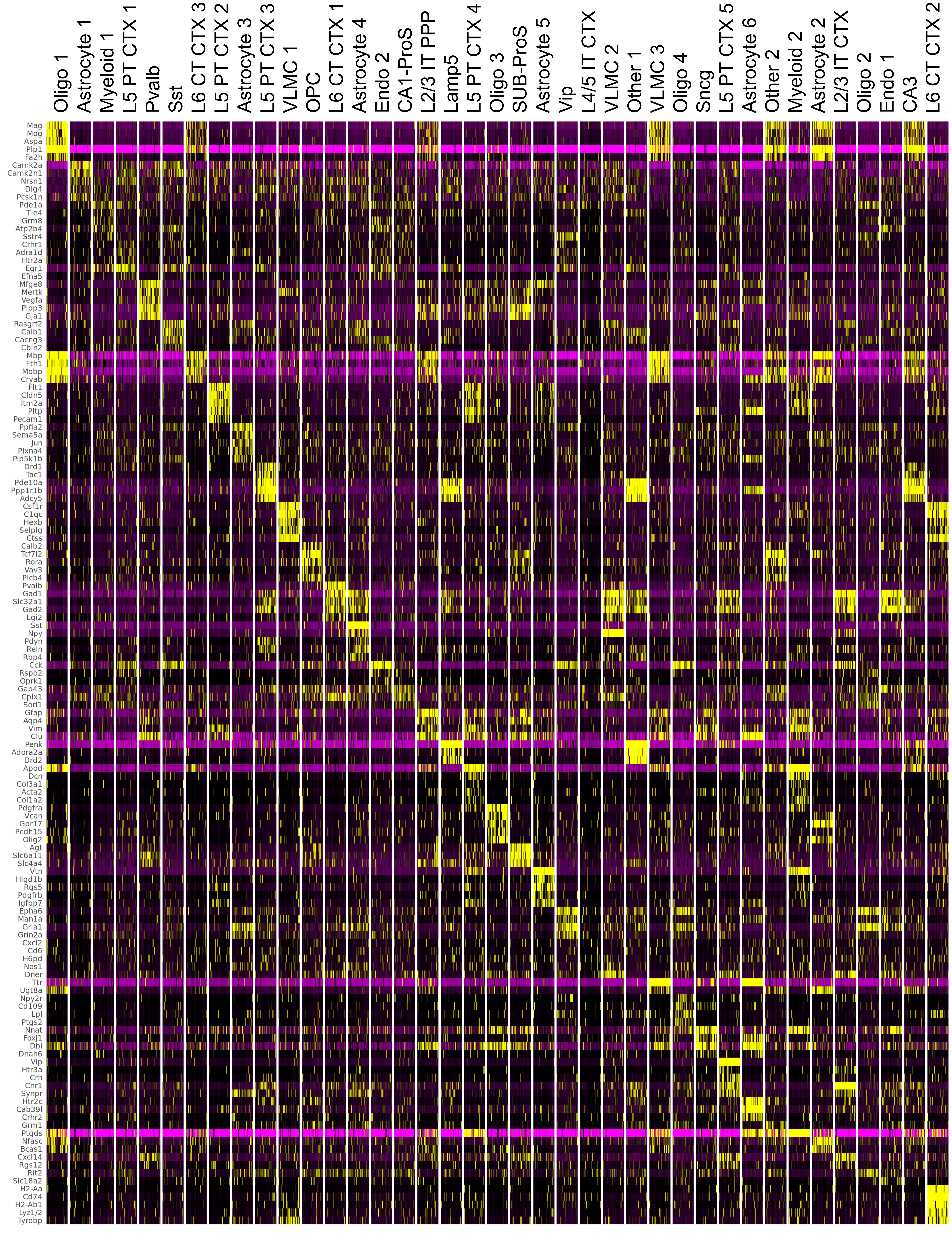

a) UMAP feature plot of canonical markers

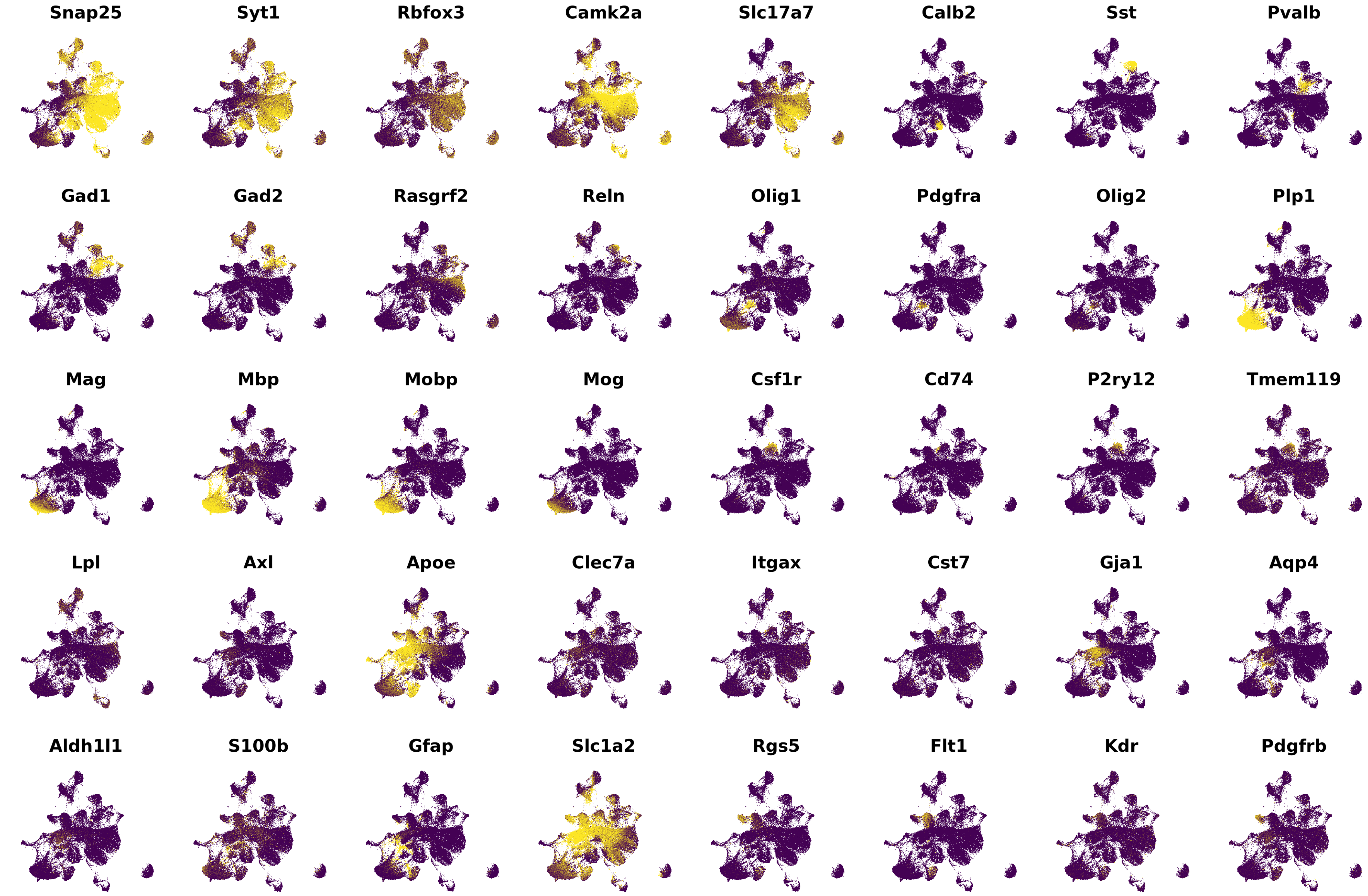


b) DEG scores, all subclusters

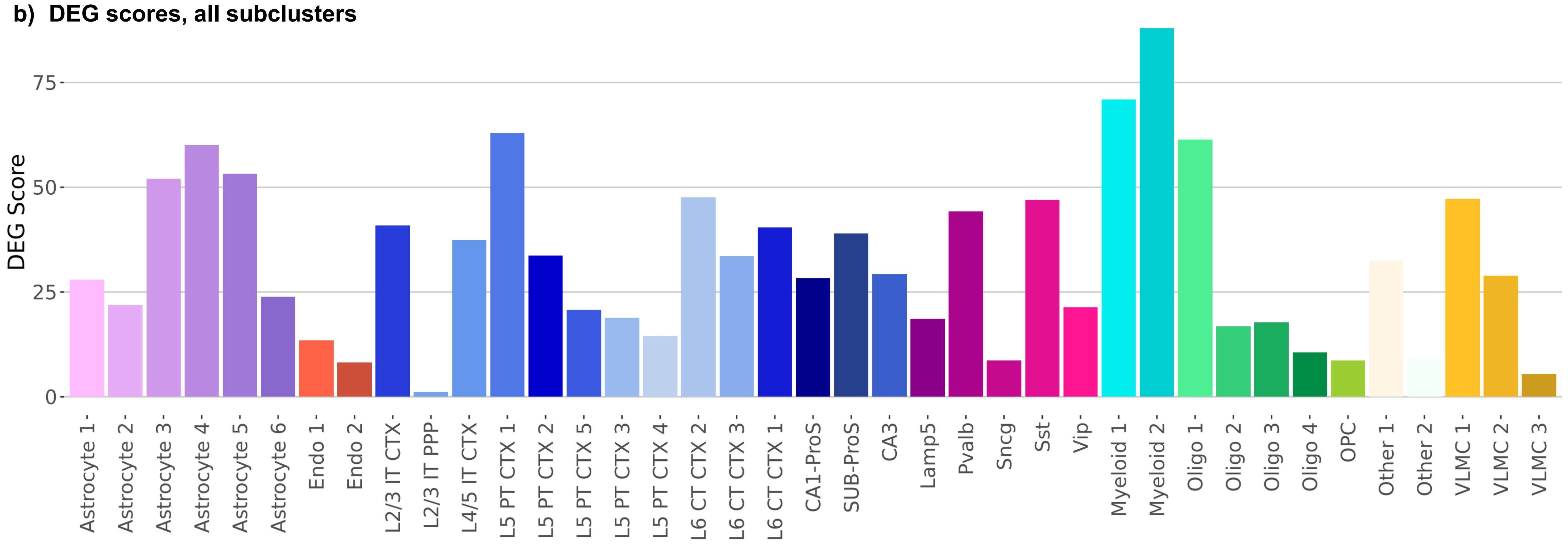

#### a) Volcano plots, *Hexb*$^{-/-}$ versus WT, all subclusters

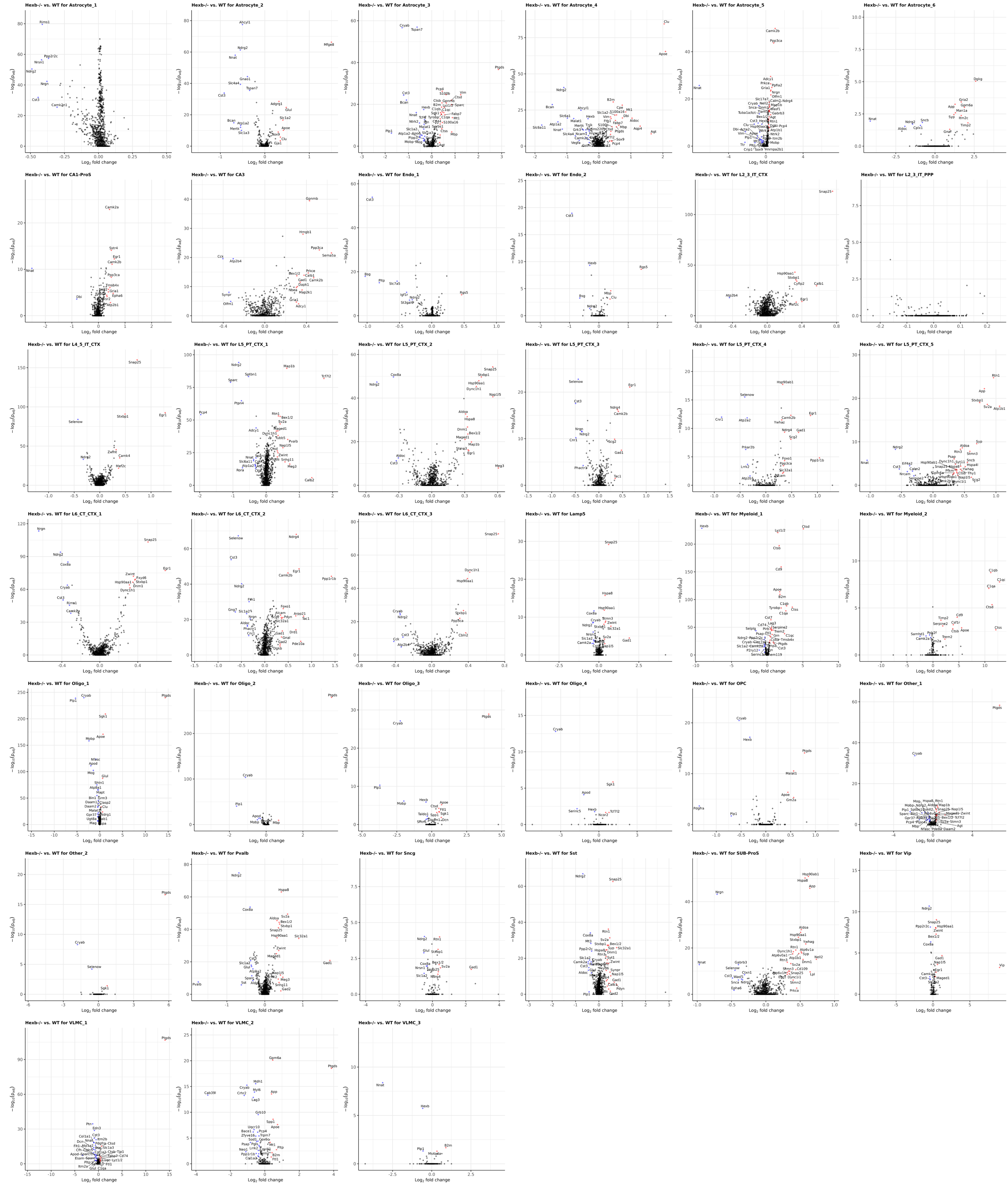

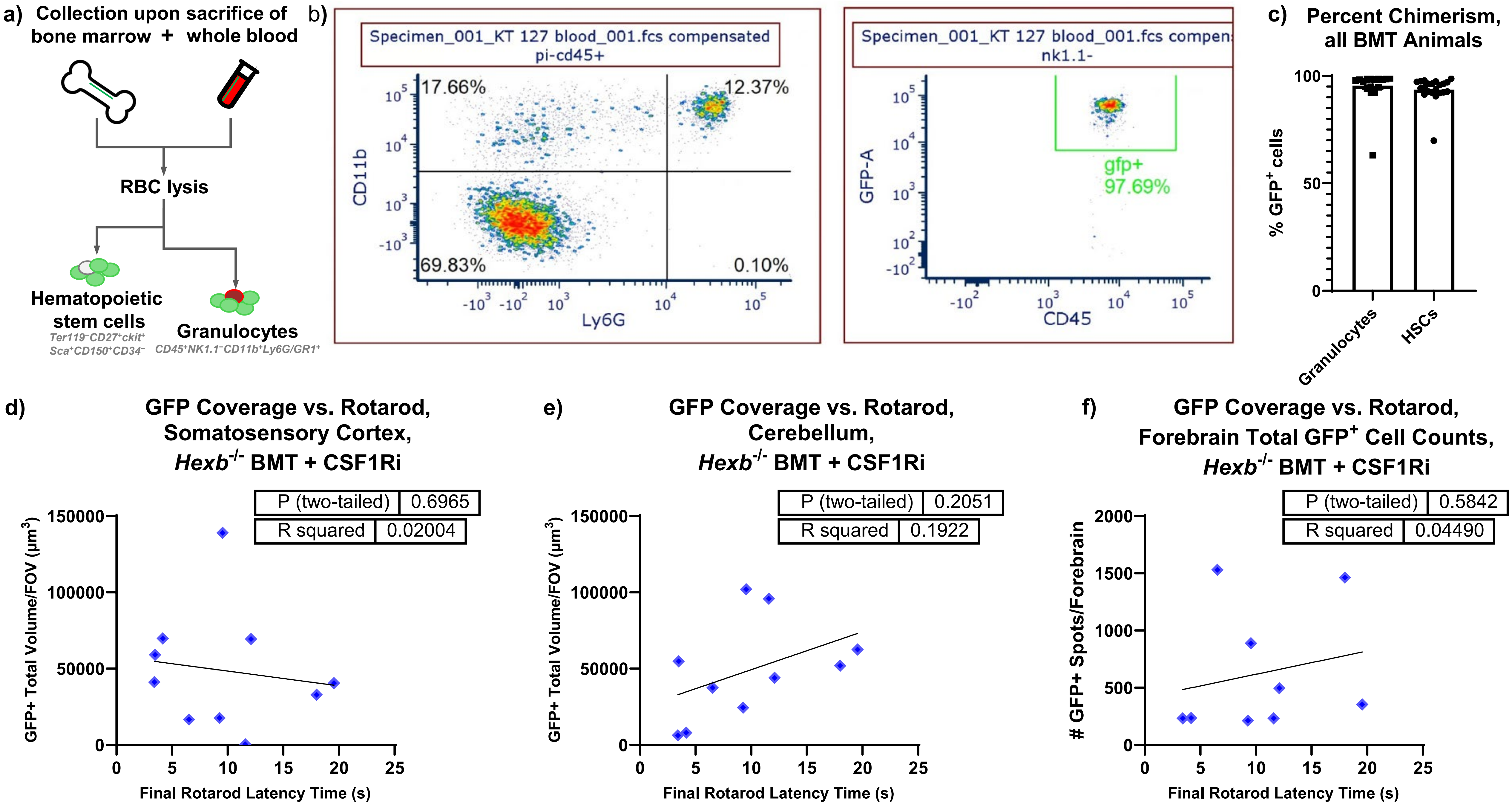

a) Collection upon sacrifice of
bone marrow + whole blood
RBC lysis
Hematopoietic stem cells
Ter119⁻CD27⁺ckit⁺ Sca⁺CD150⁺CD34⁻
Granulocytes
CD45⁺NK1.1⁻CD11b⁺Ly6G/GR1⁺
b)
Specimen_001_KT 127 blood_001.fcs compensated
pi-cd45+
17.66%
12.37%
69.83%
0.10%
CD11b
Ly6G
Specimen_001_KT 127 blood_001.fcs compen
nk1.1-
gfp+
97.69%
GFP-A
CD45
c) Percent Chimerism, all BMT Animals
% GFP⁺ cells
Granulocytes
HSCs
d) GFP Coverage vs. Rotarod, Somatosensory Cortex, Hexb⁻/⁻ BMT + CSF1Ri
P (two-tailed) 0.6965
R squared 0.02004
GFP+ Total Volume/FOV (µm³)
Final Rotarod Latency Time (s)
e) GFP Coverage vs. Rotarod, Cerebellum, Hexb⁻/⁻ BMT + CSF1Ri
P (two-tailed) 0.2051
R squared 0.1922
GFP+ Total Volume/FOV (µm³)
Final Rotarod Latency Time (s)
f) GFP Coverage vs. Rotarod, Forebrain Total GFP⁺ Cell Counts, Hexb⁻/⁻ BMT + CSF1Ri
P (two-tailed) 0.5842
R squared 0.04490
### GFP+ Spots/Forebrain
Final Rotarod Latency Time (s)

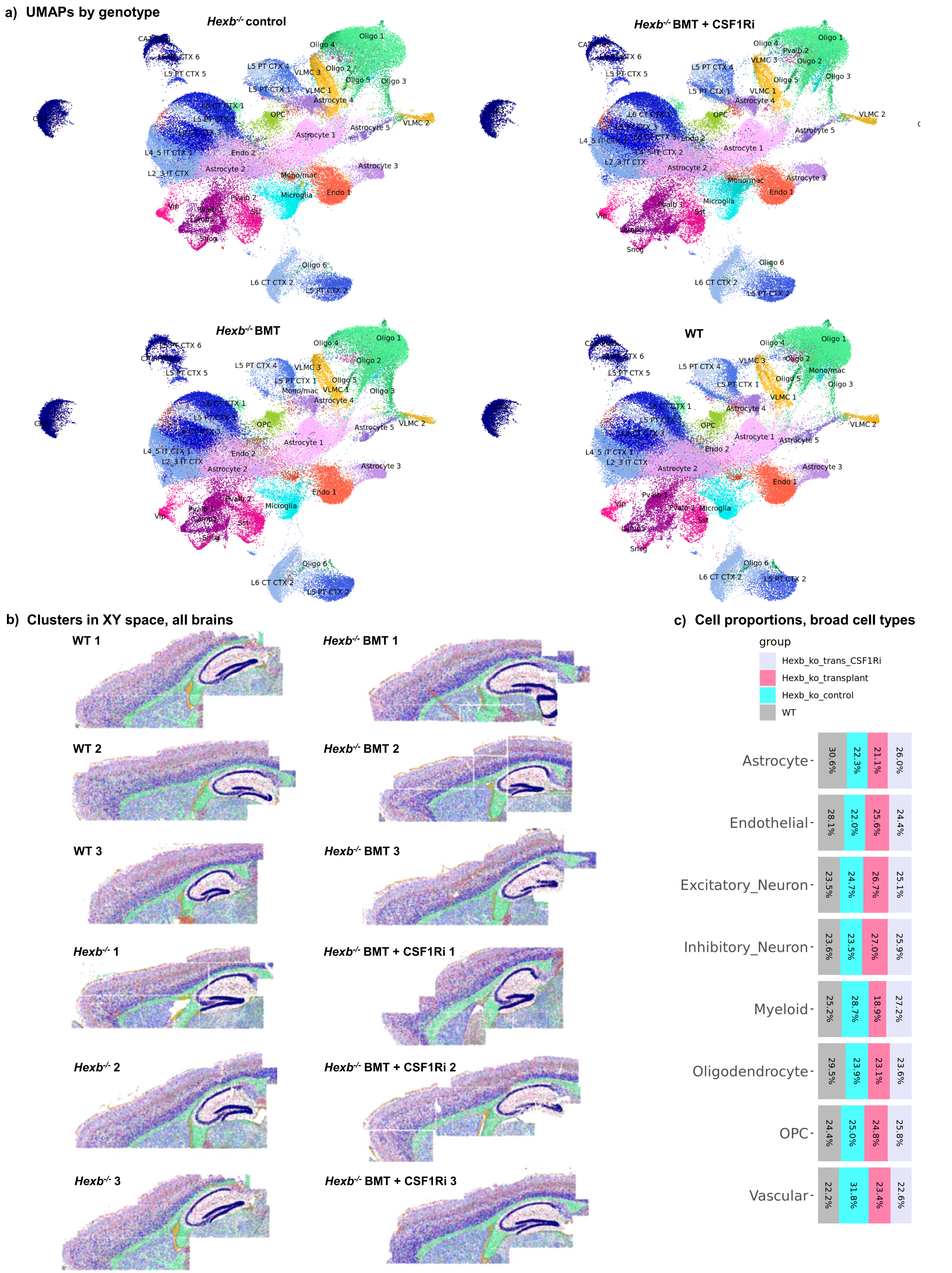
a) UMAPs by genotype
Hexb-/- control
Hexb-/- BMT + CSF1Ri
Hexb-/- BMT
WT
Oligo 1
Oligo 2
Oligo 3
Oligo 4
Oligo 5
Oligo 6
OPC
VLMC 1
VLMC 2
VLMC 3
Astrocyte 1
Astrocyte 2
Astrocyte 3
Astrocyte 4
Astrocyte 5
Endo 1
Endo 2
Mono/mac
Microglia
L5 PT CTX 1
L5 PT CTX 2
L5 PT CTX 4
L5 PT CTX 5
L6 CT CTX 1
L6 CT CTX 2
L4_5 IT CTX 1
L4_5 IT CTX 2
L2_3 IT CTX
Pvalb 1
Pvalb 2
Vip
Sst
Lamp5
Sncg
b) Clusters in XY space, all brains
WT 1
WT 2
WT 3
Hexb-/- 1
Hexb-/- 2
Hexb-/- 3
Hexb-/- BMT 1
Hexb-/- BMT 2
Hexb-/- BMT 3
Hexb-/- BMT + CSF1Ri 1
Hexb-/- BMT + CSF1Ri 2
Hexb-/- BMT + CSF1Ri 3
c) Cell proportions, broad cell types
group
Hexb_ko_trans_CSF1Ri
Hexb_ko_transplant
Hexb_ko_control
WT
Astrocyte 30.6% 22.3% 21.1% 26.0%
Endothelial 28.1% 22.0% 25.6% 24.4%
Excitatory_Neuron 23.5% 24.7% 26.7% 25.1%
Inhibitory_Neuron 23.6% 23.5% 27.0% 25.9%
Myeloid 25.2% 28.7% 18.9% 27.2%
Oligodendrocyte 29.5% 23.9% 23.1% 23.6%
OPC 24.4% 25.0% 24.8% 25.8%
Vascular 22.2% 31.8% 23.4% 22.6%

**a) Top 5 marker genes per subcluster**

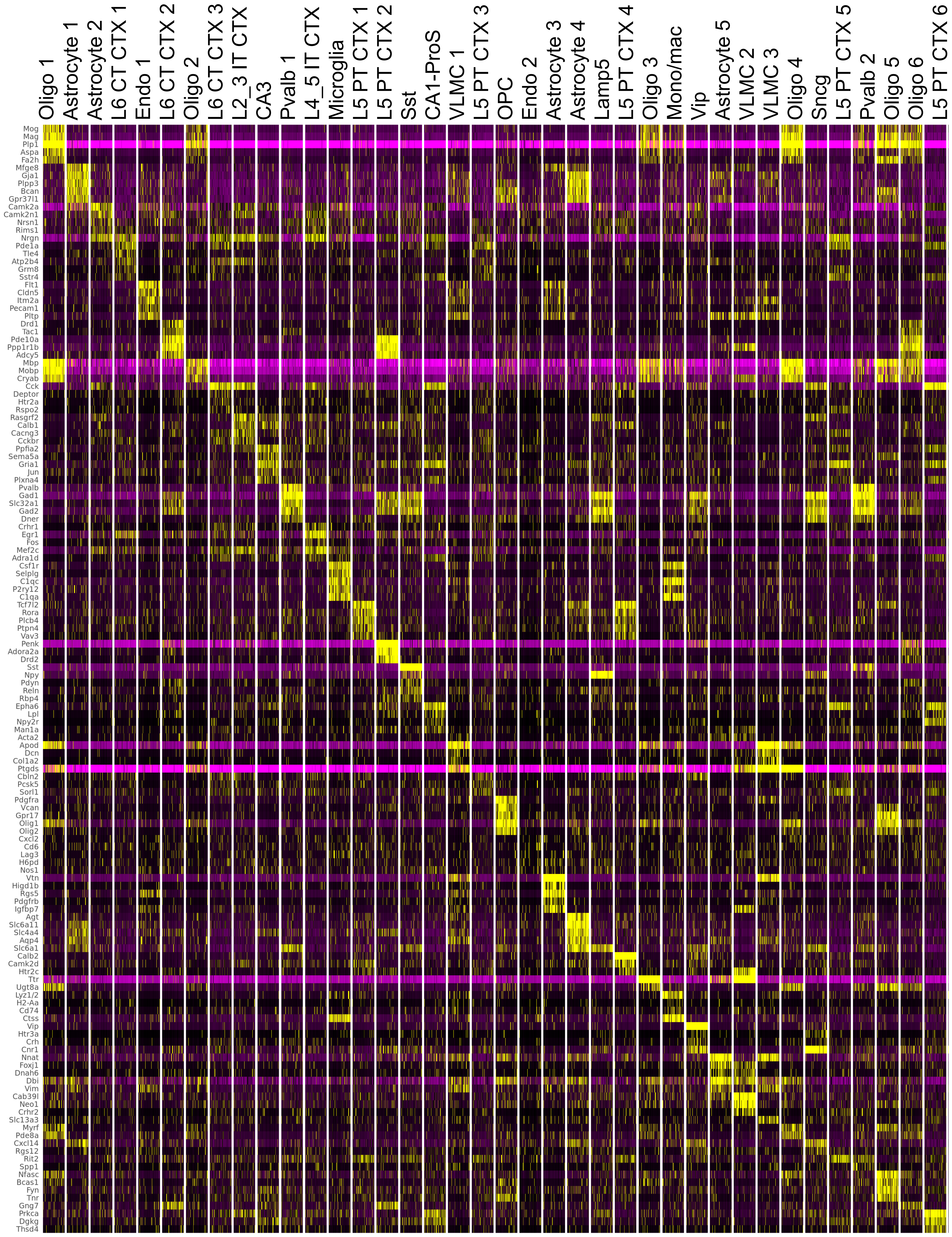

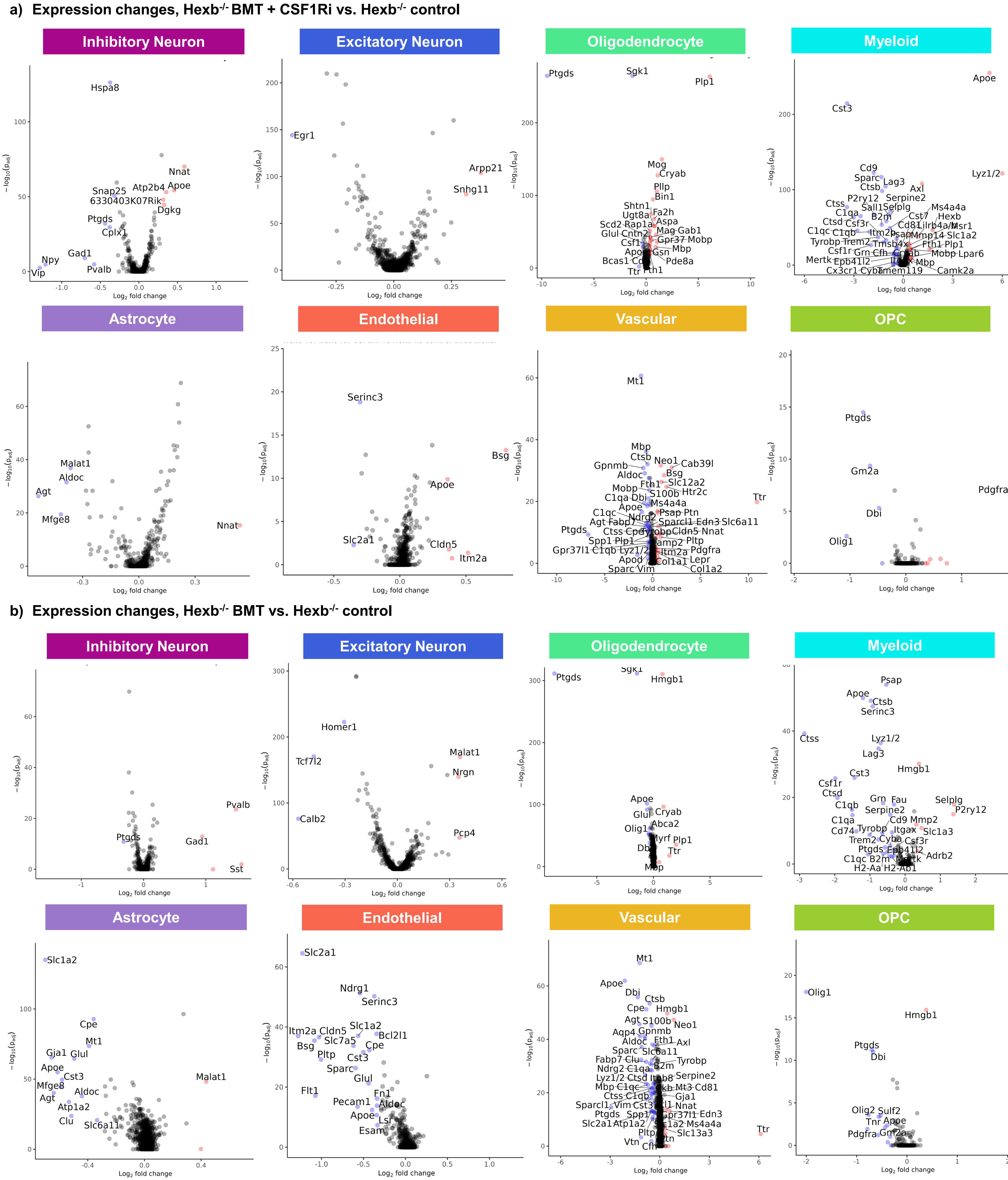

a) Expression changes, Hexb-/- BMT + CSF1Ri vs. Hexb-/- control
Inhibitory Neuron
Excitatory Neuron
Oligodendrocyte
Myeloid
Astrocyte
Endothelial
Vascular
OPC
b) Expression changes, Hexb-/- BMT vs. Hexb-/- control
Inhibitory Neuron
Excitatory Neuron
Oligodendrocyte
Myeloid
Astrocyte
Endothelial
Vascular
OPC
Log2 fold change
−log10(padj)

a) Expression changes, Hexb$^{-/-}$ BMT + CSF1Ri vs. WT control

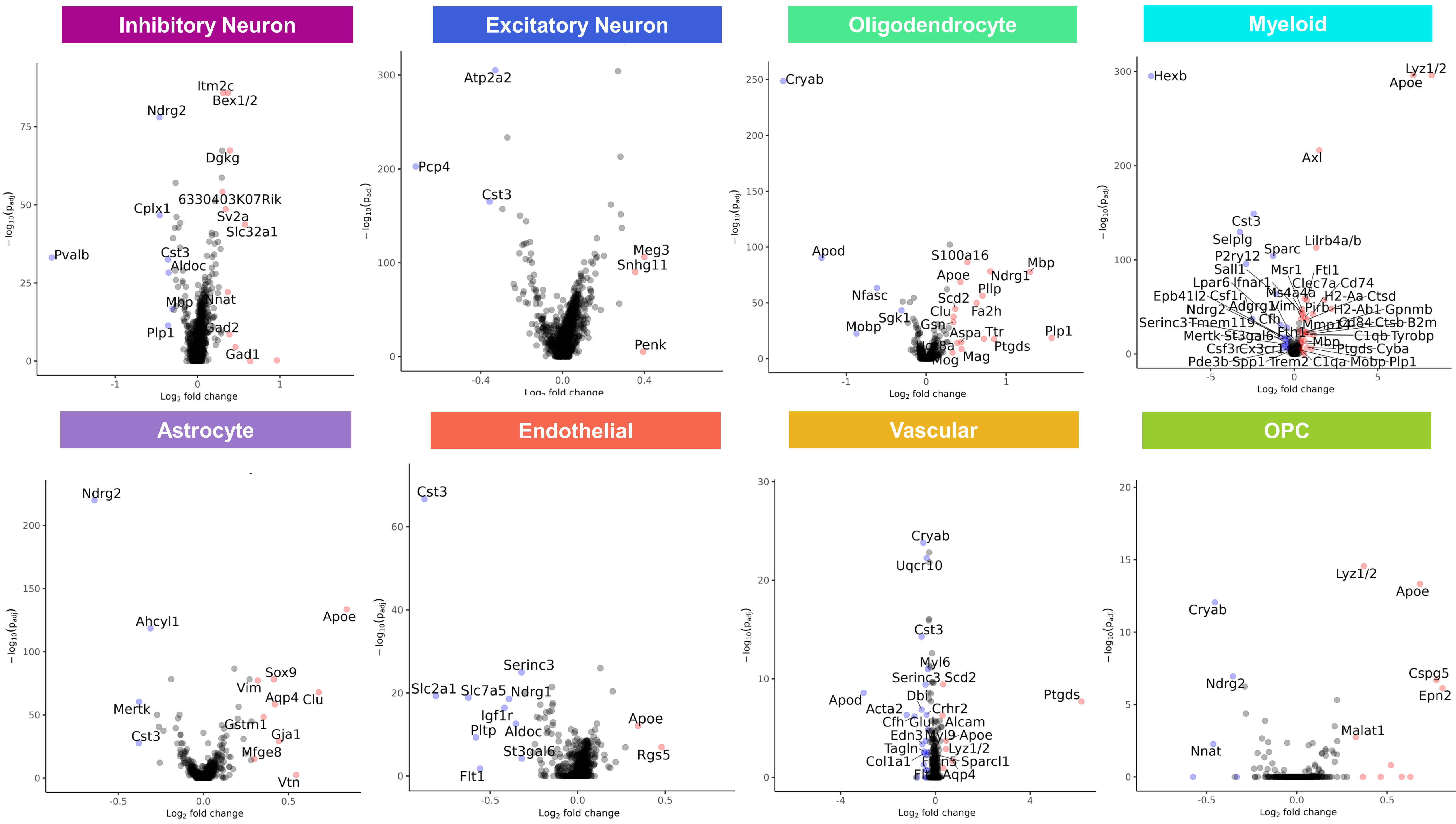


b) Expression changes, Hexb$^{-}$ BMT vs. WT control

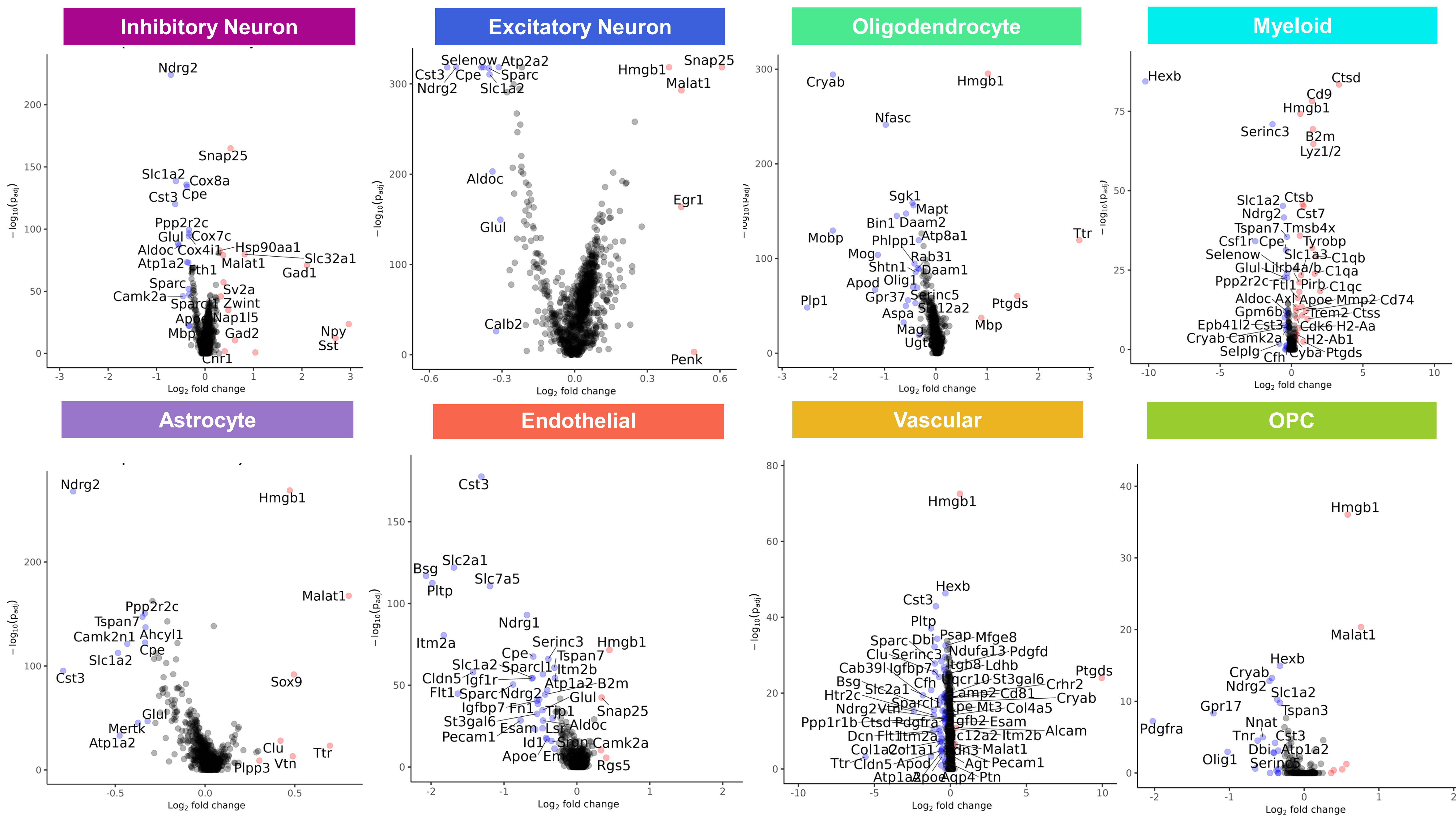

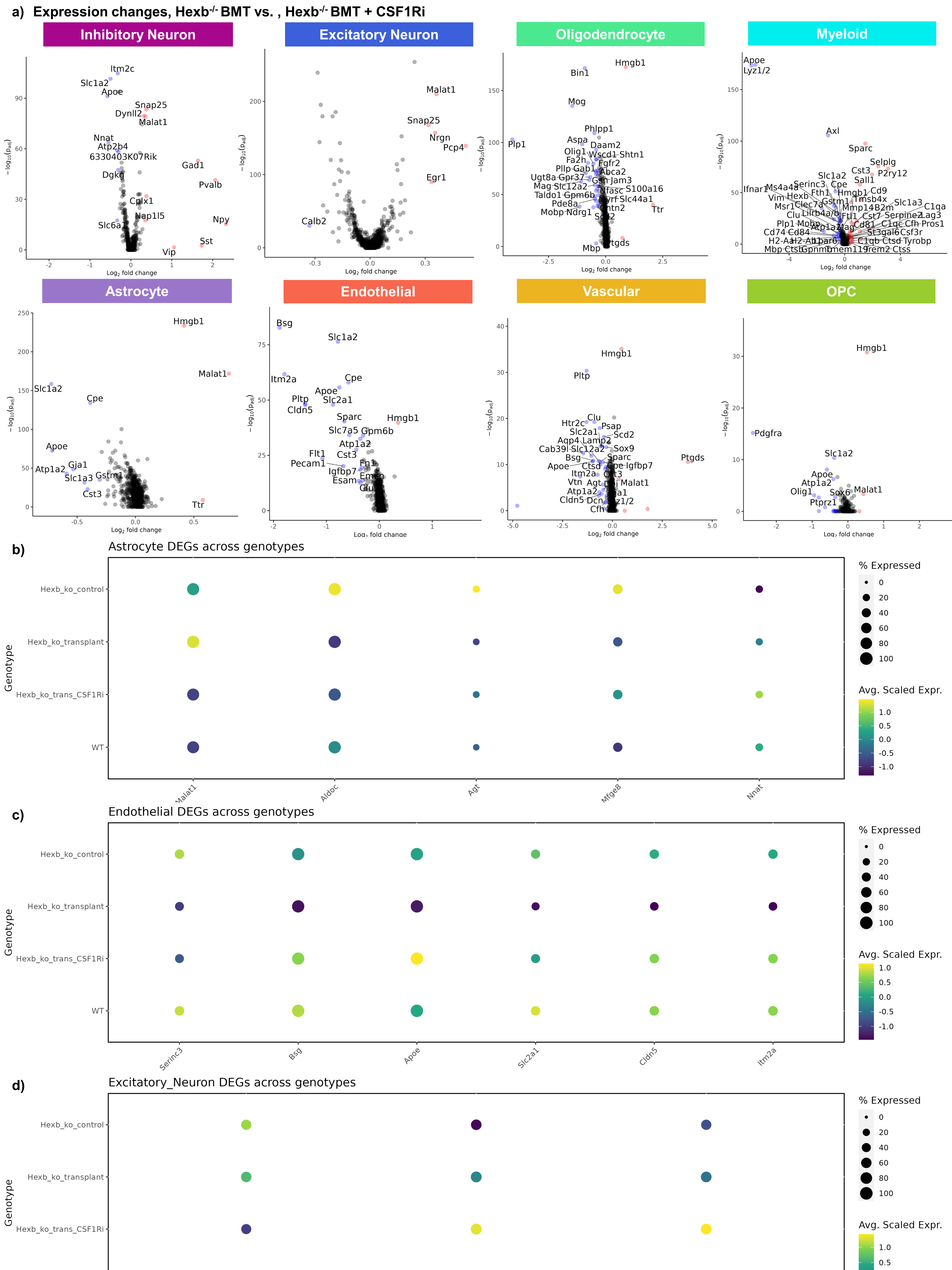
a) Expression changes, Hexb-/- BMT vs. , Hexb-/- BMT + CSF1Ri
Inhibitory Neuron
Excitatory Neuron
Oligodendrocyte
Myeloid
Astrocyte
Endothelial
Vascular
OPC
Log2 fold change
−log10(padj)
Itm2c
Slc1a2
Apoe
Snap25
Dynll2
Malat1
Nnat
Atp2b4
6330403K07Rik
Gad1
Dgkg
Pvalb
Cplx1
Nap1l5
Npy
Slc6a1
Sst
Vip
Snap25
Nrgn
Pcp4
Egr1
Calb2
Hmgb1
Bin1
Mog
Plp1
Phlpp1
Aspa
Daam2
Olig1
Wscd1
Shtn1
Fa2h
Fgfr2
Plp
Gab1
Abca2
Ugt8a
Gpr37
Gsn
Jam3
Mag
Slc12a2
Nfasc
S100a16
Taldo1
Gpm6b
Slc44a1
Pde8a
Ttr
Mobp
Ndrg1
Mbp
Lyz1/2
Axl
Sparc
Selplg
Cst3
P2ry12
Sall1
Cpe
Serinc3
Hexb
Fth1
Cd9
Slc1a3
C1qa
Lag3
Cd74
Cd84
Tyrobp
Ctss
Pdgfra
Olig1
Ptprz1
Sox6
Atp1a2
Gja1
Gstm1
Cst3
Bsg
Itm2a
Pltp
Cldn5
Slc2a1
Slc7a5
Gpm6b
Flt1
Pecam1
Fn1
Igfbp7
Esam
Clu
Htr2c
Psap
Scd2
Aqp4
Lamp2
Cab39l
Sox9
Ptgds
Ctsd
Igfbp7
Itm2a
Vtn
Agt
Dcn
Cfh
b) Astrocyte DEGs across genotypes
c) Endothelial DEGs across genotypes
d) Excitatory_Neuron DEGs across genotypes
Genotype
Hexb_ko_control
Hexb_ko_transplant
Hexb_ko_trans_CSF1Ri
WT
Malat1
Aldoc
Agt
Mfge8
Nnat
Serinc3
Bsg
Apoe
Slc2a1
Cldn5
Itm2a
% Expressed
0
20
40
60
80
100
Avg. Scaled Expr.
1.0
0.5
0.0
-0.5
-1.0

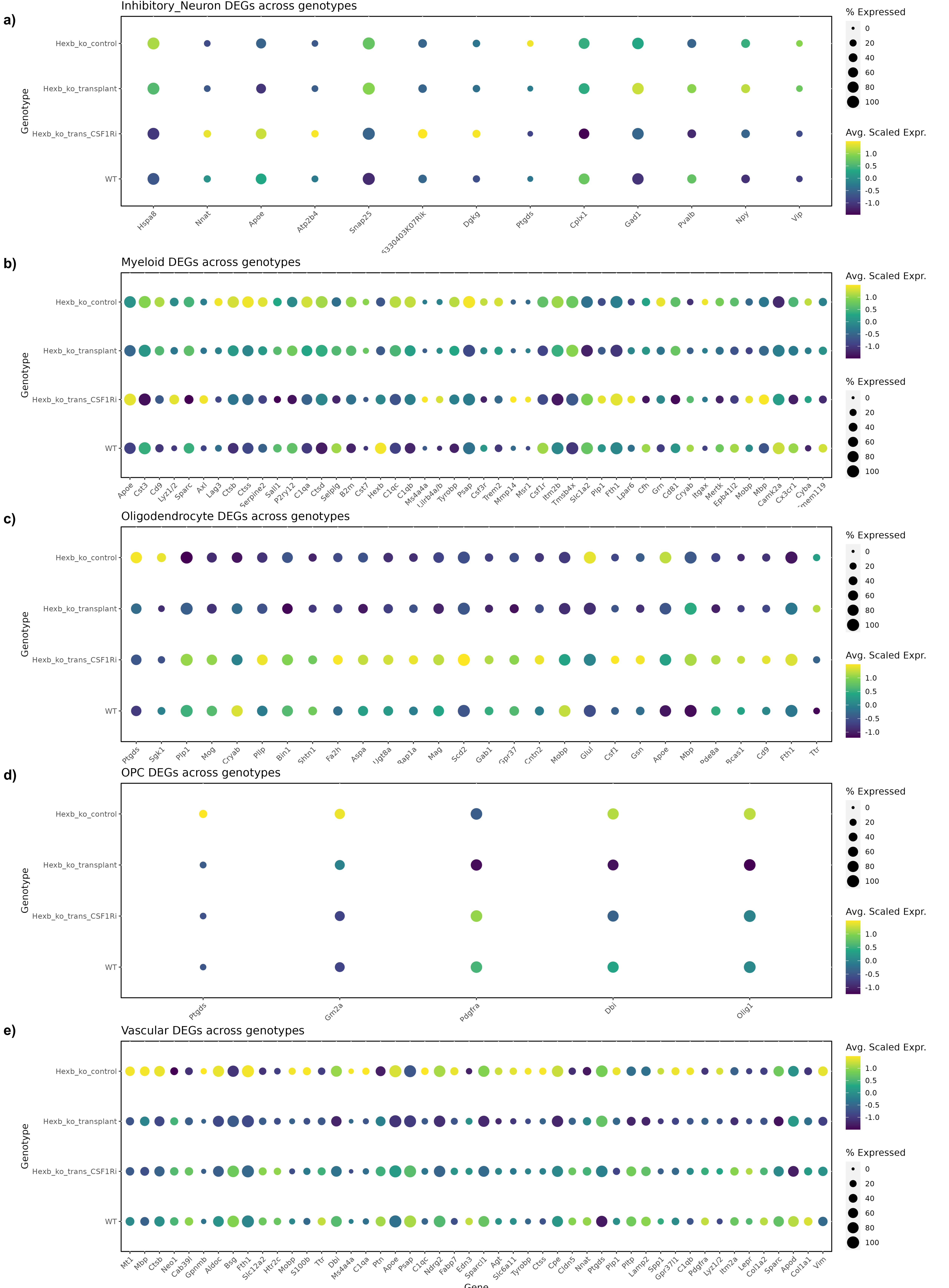
a)
Inhibitory_Neuron DEGs across genotypes
Genotype
Hexb_ko_control
Hexb_ko_transplant
Hexb_ko_trans_CSF1Ri
WT
Hspa8
Nnat
Apoe
Atp2b4
Snap25
Dgkg
Ptgds
Cplx1
Gad1
Pvalb
Npy
Vip
% Expressed
0
20
40
60
80
100
Avg. Scaled Expr.
1.0
0.5
0.0
-0.5
-1.0
b)
Myeloid DEGs across genotypes
Apoe
Cst3
Cd9
Lyz1/2
Sparc
Axl
Lag3
Ctsb
Ctss
Serpine2
Sall1
P2ry12
C1qa
Ctsd
Selplg
B2m
Cst7
Hexb
C1qc
C1qb
Ms4a4a
Tyrobp
Psap
Csf3r
Trem2
Mmp14
Msr1
Csf1r
Itm2b
Tmsb4x
Slc1a2
Plp1
Fth1
Lpar6
Cfh
Grn
Cd81
Cryab
Itgax
Mertk
Epb41l2
Mobp
Mbp
Camk2a
Cx3cr1
Cyba
c)
Oligodendrocyte DEGs across genotypes
Ptgds
Sgk1
Plp1
Mog
Cryab
Pllp
Bin1
Shtn1
Fa2h
Aspa
Ugt8a
Mag
Scd2
Gab1
Gpr37
Cntn2
Mobp
Glul
Csf1
Gsn
Apoe
Mbp
Bcas1
Cd9
Fth1
Ttr
d)
OPC DEGs across genotypes
Ptgds
Gm2a
Pdgfra
Dbi
Olig1
e)
Vascular DEGs across genotypes
Mt1
Mbp
Ctsb
Neo1
Cab39l
Gpnmb
Aldoc
Bsg
Fth1
Slc12a2
Htr2c
Mobp
S100b
Ttr
Dbi
Ms4a4a
C1qa
Ptn
Apoe
Psap
C1qc
Ndrg2
Fabp7
Edn3
Sparcl1
Agt
Slc6a11
Tyrobp
Ctss
Cpe
Cldn5
Nnat
Ptgds
Plp1
Pltp
Lamp2
Spp1
Gpr37l1
C1qb
Pdgfra
Lyz1/2
Itm2a
Lepr
Col1a2
Sparc
Apod
Col1a1
Vim
Gene

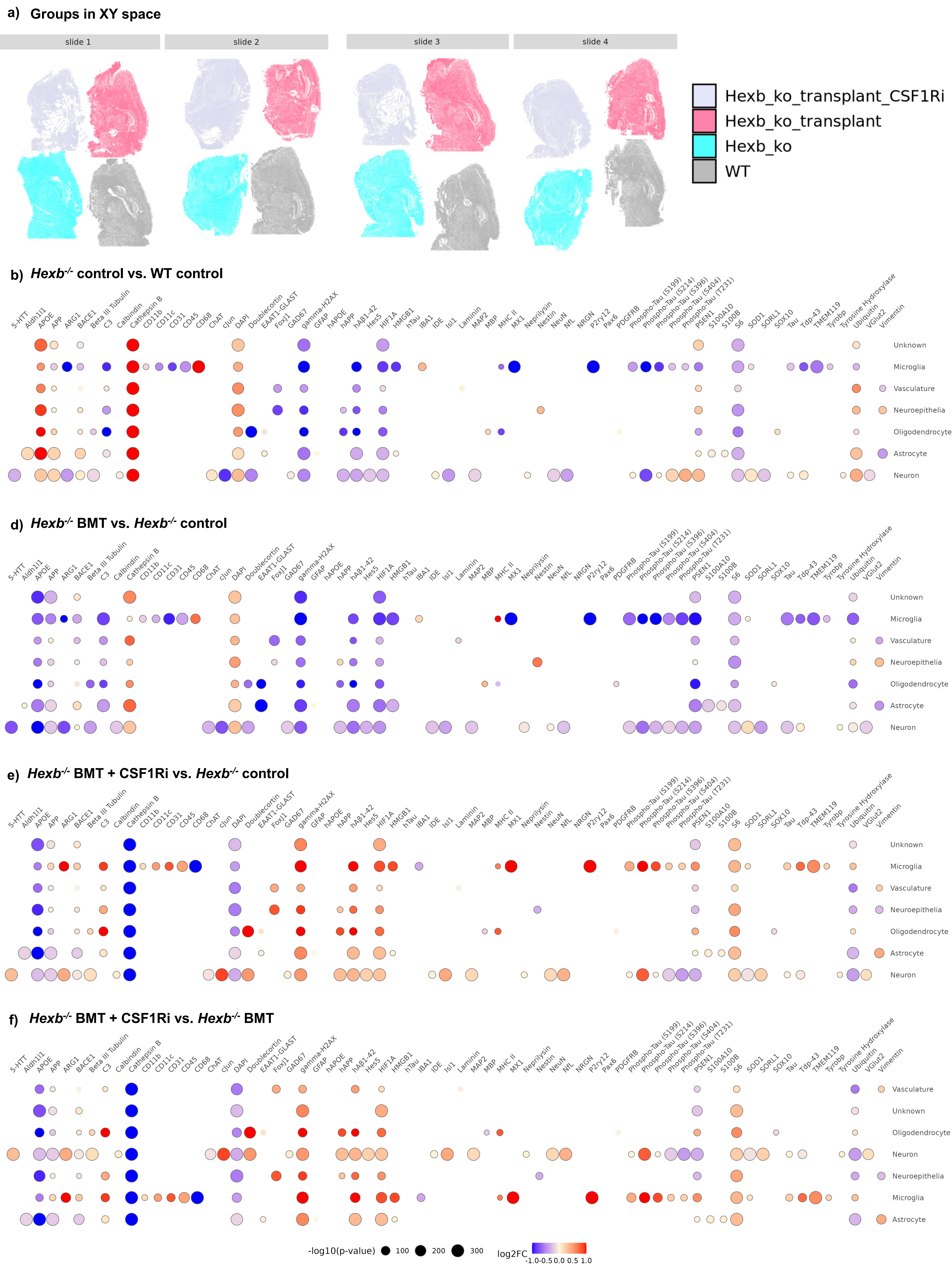

a) Groups in XY space
slide 1
slide 2
slide 3
slide 4
Hexb_ko_transplant_CSF1Ri
Hexb_ko_transplant
Hexb_ko
WT
b) Hexb-/- control vs. WT control
d) Hexb-/- BMT vs. Hexb-/- control
e) Hexb-/- BMT + CSF1Ri vs. Hexb-/- control
f) Hexb-/- BMT + CSF1Ri vs. Hexb-/- BMT
5-HTT
Aldh1l1
APOE
APP
ARG1
BACE1
Beta III Tubulin
C3
Calbindin
Cathepsin B
CD11b
CD11c
CD31
CD45
CD68
ChAT
cJun
DAPI
Doublecortin
EAAT1-GLAST
FoxJ1
GAD67
gamma-H2AX
GFAP
hAPOE
hAPP
hAβ1-42
Hes5
HIF1A
HMGB1
hTau
IBA1
IDE
Isl1
Laminin
MAP2
MBP
MHC II
MX1
Neprilysin
Nestin
NeuN
NfL
NRGN
P2ry12
Pax6
PDGFRB
Phospho-Tau (S199)
Phospho-Tau (S214)
Phospho-Tau (S396)
Phospho-Tau (S404)
Phospho-Tau (T231)
PSEN1
S100A10
S100B
S6
SOD1
SORL1
SOX10
Tau
Tdp-43
TMEM119
Tyrobp
Tyrosine Hydroxylase
Ubiquitin
VGlut2
Vimentin
Unknown
Microglia
Vasculature
Neuroepithelia
Oligodendrocyte
Astrocyte
Neuron
-log10(p-value)
100
200
300
log2FC
-1.0 -0.5 0.0 0.5 1.0
